# Ancient DNA reconstructs the genetic legacies of pre-contact Puerto Rico communities

**DOI:** 10.1101/765685

**Authors:** Maria A. Nieves-Colón, William J. Pestle, Austin W. Reynolds, Bastien Llamas, Constanza de la Fuente, Kathleen Fowler, Katherine M. Skerry, Edwin Crespo-Torres, Carlos D. Bustamante, Anne C. Stone

## Abstract

Indigenous peoples have occupied the island of Puerto Rico since at least 3000 B.C. Due to the demographic shifts that occurred after European contact, the origin(s) of these ancient populations, and their genetic relationship to present-day islanders, are unclear. We use ancient DNA to characterize the population history and genetic legacies of pre-contact Indigenous communities from Puerto Rico. Bone, tooth and dental calculus samples were collected from 124 individuals from three pre-contact archaeological sites: Tibes, Punta Candelero and Paso del Indio. Despite poor DNA preservation, we used target enrichment and high-throughput sequencing to obtain complete mitochondrial genomes (mtDNA) from 45 individuals and autosomal genotypes from two individuals. We found a high proportion of Native American mtDNA haplogroups A2 and C1 in the pre-contact Puerto Rico sample (40% and 44%, respectively). This distribution, as well as the haplotypes represented, support a primarily Amazonian South American origin for these populations, and mirrors the Native American mtDNA diversity patterns found in present-day islanders. Three mtDNA haplotypes from pre-contact Puerto Rico persist among Puerto Ricans and other Caribbean islanders, indicating that present-day populations are reservoirs of pre-contact mtDNA diversity. Lastly, we find similarity in autosomal ancestry patterns between pre-contact individuals from Puerto Rico and the Bahamas, suggesting a shared component of Indigenous Caribbean ancestry with close affinity to South American populations. Our findings contribute to a more complete reconstruction of pre-contact Caribbean population history and explore the role of Indigenous peoples in shaping the biocultural diversity of present-day Puerto Ricans and other Caribbean islanders.

## Introduction

Puerto Rico is the smallest of the Greater Antilles (Figure 1), the northernmost island grouping of the Caribbean archipelago. The archaeological record of the island’s prehistoric communities suggests a dynamic population history with multiple peopling events, frequent migration, and expanding population settlements (Rouse 1992; Rodríguez Ramos 2010). However, different lines of evidence produce conflicting results about the origins and number of these migrations and the role of genetic admixture in pre-contact Caribbean population history (Chanlatte Baik 2003; Rodríguez Ramos, et al. 2013).

**Figure 1.**
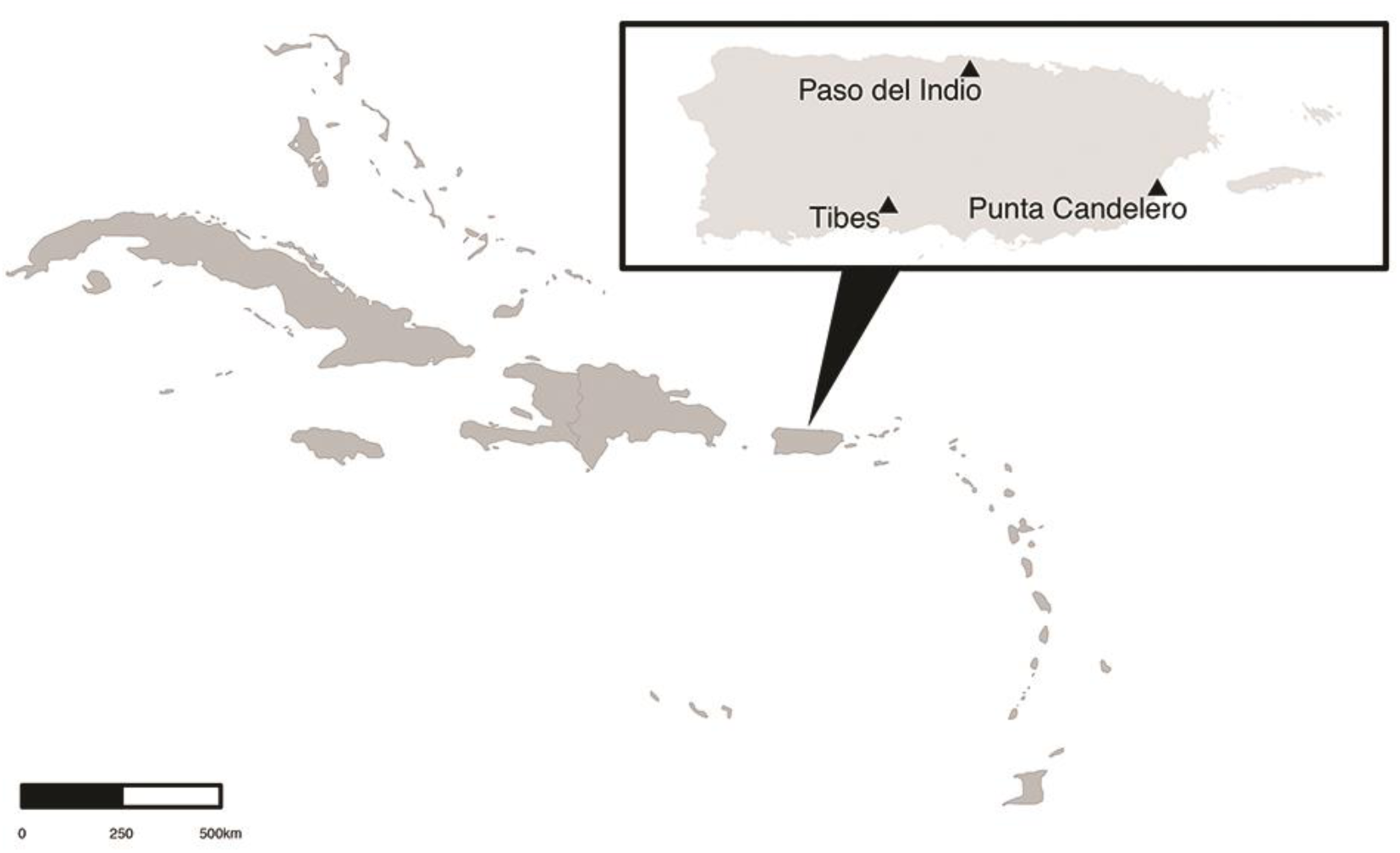
Map of Puerto Rico and the Antilles. Triangles are approximate location of pre-contact sites.

Archaeological evidence indicates humans first arrived in the Antilles by approximately 7000 B.C., reaching Puerto Rico before 3000 B.C. (Burney, et al. 1994; Rodríguez Ramos, et al. 2013). Due to the abundance of stone tools at these early sites, early settlers are known as the first representatives of the Caribbean Lithic Age, also referred to as Archaic or pre-Arawak. The origins of these first peoples and the route(s) they used to enter the Antilles are largely unknown. Based on similarities in material culture, Lithic Age populations may have originated in South America, Central America or the Isthmo-Colombian region (Rouse 1992; Rodríguez Ramos 2010; Rodríguez Ramos, et al. 2013).

Around 500 B.C., a new group of peoples with elaborate ceramic technology and large-scale agriculture entered the Antilles, arriving into Puerto Rico by 200 B.C. Archaeological and ethnohistoric evidence suggests these were likely Arawak-speakers from the Orinoco River delta region in South America (Chanlatte Baik 2003). Their arrival and the technological and cultural changes they introduced launched the Caribbean Ceramic Age (CA). Although it was initially thought these migrants displaced pre-existing Lithic/Archaic populations (Rouse 1992), more recent archaeological evidence suggests admixture occurred, and both groups contributed to the ancestry of Late Ceramic Age (LCA) peoples who expanded and diversified across the Antilles after A.D. 600 (Wilson 1999; Keegan and Rodriguez Ramos 2005). On Puerto Rico and other islands, the LCA was characterized by increased social stratification, the possible rise of organized chiefdoms, and the emergence of regional technological and artistic traditions (Rouse 1992; Chanlatte Baik 2003; Curet, et al. 2004). Material and isotopic evidence suggest that inter-island interaction and mobility expanded during the LCA (Laffoon and Hoogland 2012; Mol 2013). By the time of European contact in the 15^th^ century, multiple ethnic groups with varying levels of social and political complexity existed in the Antilles (Wilson 2007). In Puerto Rico, and other islands, these groups are known as the “Taino” (although see Curet (2014) for discussion of the problematic Taino ethnonym).

European colonialism altered the demography of Puerto Rico and the Caribbean. Forced relocations, disease, and slavery decimated native populations. The establishment of plantation and mining economies increased migration from Europe, and systems of forced labor brought peoples from Africa, Asia, and other parts of the Americas to the Antilles (Rogozinski 2008). Despite extensive ethnohistorical research (Anderson-Córdova 2005; Anderson-Córdova 2010), the extent to which Puerto Rico’s native Indigenous communities resisted, survived and were transformed by colonization is unclear. Their biocultural connection to present-day islanders is also disputed. Historical claims of extinction, based largely on colonial era censuses, are strongly opposed by islanders who assert cultural affiliation and direct descendance from native pre-contact communities (Haslip-Viera 2001; Forte 2006; Benn-Torres 2014).

Previous research has attempted to address these debates by characterizing Native American genome segments found in present-day, admixed Puerto Ricans (Martínez-Cruzado, et al. 2001; Martínez-Cruzado 2002; Martínez-Cruzado, et al. 2005; Martínez-Cruzado 2010; Gravel, et al. 2013; Moreno-Estrada, et al. 2013; Vilar, et al. 2014). Caribbean islanders have varying proportions of African, European, Native American, and Asian genetic ancestry. These patterns differ among populations and reflect the sex-biased nature of colonial and post-colonial demographic processes (Bryc, et al. 2010; Moreno-Estrada, et al. 2013). For instance, Puerto Ricans have high proportions of European and African ancestry in autosomal and Y-chromosome loci, but large proportions of Native American ancestry in the mitochondrial genome (Martínez-Cruzado, et al. 2005; Via, et al. 2011; Vilar, et al. 2014). This Native American ancestry component, presumed to originate from ancient island populations, has been used as a proxy to reconstruct pre-contact genetic variation (Martínez-Cruzado 2010; Gravel, et al. 2013).

However, present-day Caribbean genomes may not fully represent pre-contact genetic diversity. Evolutionary forces such as drift and natural selection affect lineage survival in descendant populations and bias reconstructions of ancient demography. Recent population replacements can also mask the genetic signal of ancient groups (Pickrell and Reich 2014). In Puerto Rico for example, historical documents indicate that Indigenous peoples from other parts of the Americas were imported as slave labor during the early 16^th^ century (Anderson-Córdova 2005; Anderson-Córdova 2010), yet their contribution to the gene pool of present-day Puerto Ricans is currently unknown (Martínez-Cruzado 2010). Many of these gaps could be filled with ancient DNA (aDNA), but poor preservation in pre-contact Caribbean archaeological contexts has limited aDNA research to partial fragments of mitochondrial DNA (mtDNA) (Lalueza-Fox, et al. 2001; Lalueza-Fox, et al. 2003; Mendisco, et al. 2015) or genome-wide data from few individuals (Schroeder, et al. 2015; Schroeder, et al. 2018).

To address these limitations, in this study we use paleogenomics to examine directly the genetic diversity of pre-contact populations from Puerto Rico. Using target enrichment and high-throughput sequencing, we recovered 45 complete mtDNA genomes and two partial autosomal genomes from 124 individuals sampled from three pre-contact sites. To draw inferences on the origins of these communities, and their relationship to coeval Caribbean populations, we compare these sequences to genetic data from ancient and present-day populations from the Caribbean islands and continental Americas. Contextualized within the framework of previous genetic and archaeological research, our findings prove instrumental for reconstructing the population history of pre-contact Puerto Rico and lead to a better understanding of the biocultural links between ancient Indigenous Caribbean communities and present-day islanders.

## Results

### Preservation and ancient DNA recovery

We sampled 124 human skeletal remains excavated from CA and LCA archeological contexts at three sites in Puerto Rico: Punta Candelero (n=34), Tibes (n=46) and Paso del Indio (n=44) (Figure 1; Figures S1-S2). Direct radiocarbon dates previously obtained for 81 of the individuals ranged between A.D. 500-1300 (Pestle 2010) (Table S1; Figure S3). The adverse preservation conditions of the Caribbean severely restricted aDNA recovery. Samples from Tibes had the worst mtDNA preservation (Figure S4). After enrichment and Illumina sequencing, complete mtDNA genomes with ≥5X average read depth and ≥98% genome coverage were recovered from 45 individuals (36%): Punta Candelero (n=16), Tibes (n=5) and Paso del Indio (n=24) (Table S2-S3). Five of these mtDNA genomes were obtained through multiple sequencing runs.

In addition to mitochondrial enrichment, 35 samples were shotgun sequenced and 22 of these were subjected to whole genome enrichment (WGE). Endogenous DNA content was extremely low for all samples even after WGE. Analyses of autosomal genotypes were performed solely on two samples from Paso del Indio with the highest average read depth and genome coverage after multiple rounds of enrichment and sequencing: PI-51 (0.16X, 13.89% genome coverage) and PI-420a (0.27X, 19.16% genome coverage) (Table S4-S5). Reads obtained for all 45 samples selected for mtDNA analyses and both samples selected for autosomal DNA analyses had damage patterns characteristic of authentic aDNA such as short fragments (<200 bp) and high rates of C-to-T and A-to-G transitions at 5’ and 3’ DNA fragment ends, respectively (Briggs, et al. 2007) (Figure S5-S6). For two mtDNA enriched samples with evidence of contamination (PC-443 and PC-448), endogenous reads were recovered after additional filtering with PMDtools (Skoglund, et al. 2014) (Table S6; Figure S7). Average DNA fragment length was approximately 65 bp and estimated contamination ranged between 1-9% for all analyzed samples.

### MtDNA diversity

Three of the five characteristic Native American mtDNA haplogroups were found in the pre-contact Puerto Rico (PC-PR) sample: A2, C1 and D1. When considering complete mtDNA variants, we identified 29 haplotypes in 45 individuals (Table S7). 84% were classified into haplogroups A2 and C1. The most frequent sub-haplogroup was C1b2, accounting for 33% of all mtDNA lineages (Figure 2). When considering only HVR-1 sequences, lineage variation is collapsed into 18 haplotypes.

**Figure 2.**
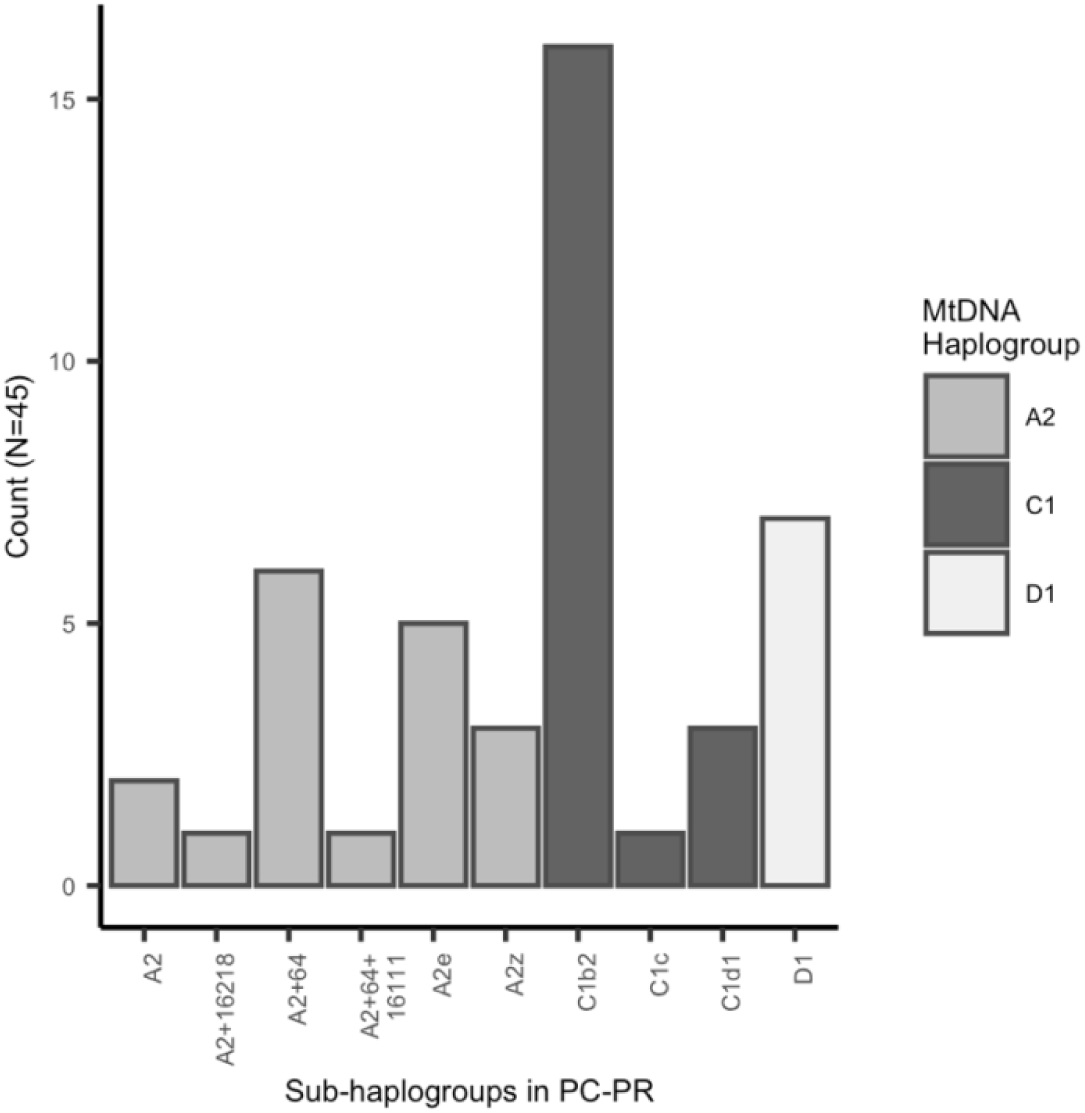
MtDNA sub-haplogroups in the PC-PR sample.

We tested for differences in complete mtDNA haplotype diversity between the three pre-contact sites by calculating an exact test of population differentiation and pairwise Φ_st_ measures. We found no significant differences (*p*>*0.05*), suggesting no genetic structure existed between site communities (Table S8-S9, Figure S8). We also found no significant relationship between genetic and temporal distance in the complete PC-PR sample (*z*=*182.65, p*=*0.880*) (Figure S9). Thus, subsequent analyses were conducted assuming a panmictic pre-contact island population.

Intra-population summary statistics calculated with complete mtDNA sequences indicate that the PC-PR sample has low haplotype (*Hd*= 0.902) and nucleotide diversity (π=0.001) compared to most reference populations, except for several groups with known low diversity such as the Surui and Karitiana of Amazonia (Wang, et al. 2007) (Table S10-S11, Figure S10A-B). A second test restricted to HVR-1 for comparison with available comparative datasets, found that PC-PR had low haplotype (*Hd*=0.942) and nucleotide diversity (π=0.009) relative to other Caribbean populations (Table S12, Figure S10C-D).

### MtDNA inter-population differentiation

We measured inter-population genetic distance and differentiation by comparing complete mtDNA haplotypes between PC-PR and 46 reference populations from the Americas (Table S10). Exact tests found significant differences in haplotype frequencies between PC-PR and 17 populations, but the null hypothesis of panmixia could not be rejected for the remaining 29 comparisons, including between PC-PR and present-day Puerto Ricans (Table S13). Pairwise comparisons of Φ_st_ measures calculated with complete mtDNA sequences found the lowest subdifferentiation values between PC-PR and Indigenous populations from northwest Amazonia and the Andes (Figure S11). For ten of these comparisons, we were unable to reject the null hypothesis of no differentiation (Table S14). Φ_st_ inter-population distances are visually represented with non-metric multidimensional scaling (MDS) in Figure 3. The MDS patterns are broadly recapitulated in the correspondence analysis plot in Figure 4 which clusters populations based on haplogroup frequency. PC-PR falls in the upper left quadrant of the plot, clustering with Amazonian populations carrying high frequencies of haplogroup C. We note that Φ_st_ distances between PC-PR and Puerto Ricans were low (Φ_st_ =*0.0441, p*=*0.018*), but significantly different. This suggests that although haplotype frequencies are similar between them, some differentiation exists.

**Figure 3.**
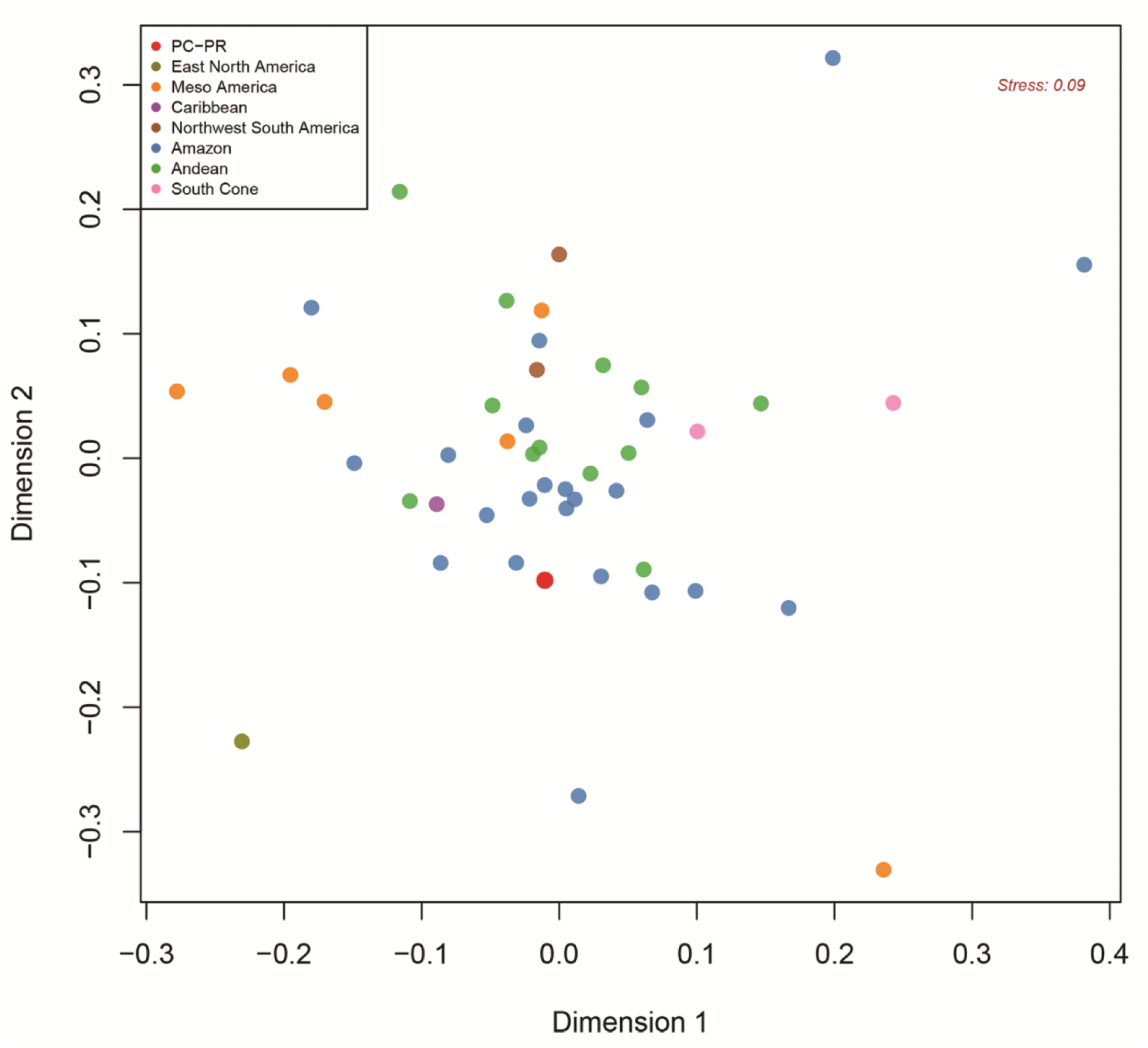
Non-metric MDS plot of complete mtDNA Φ_st_ distances between PC-PR and 46 ancient and present-day populations from the Americas.

**Figure 4.**
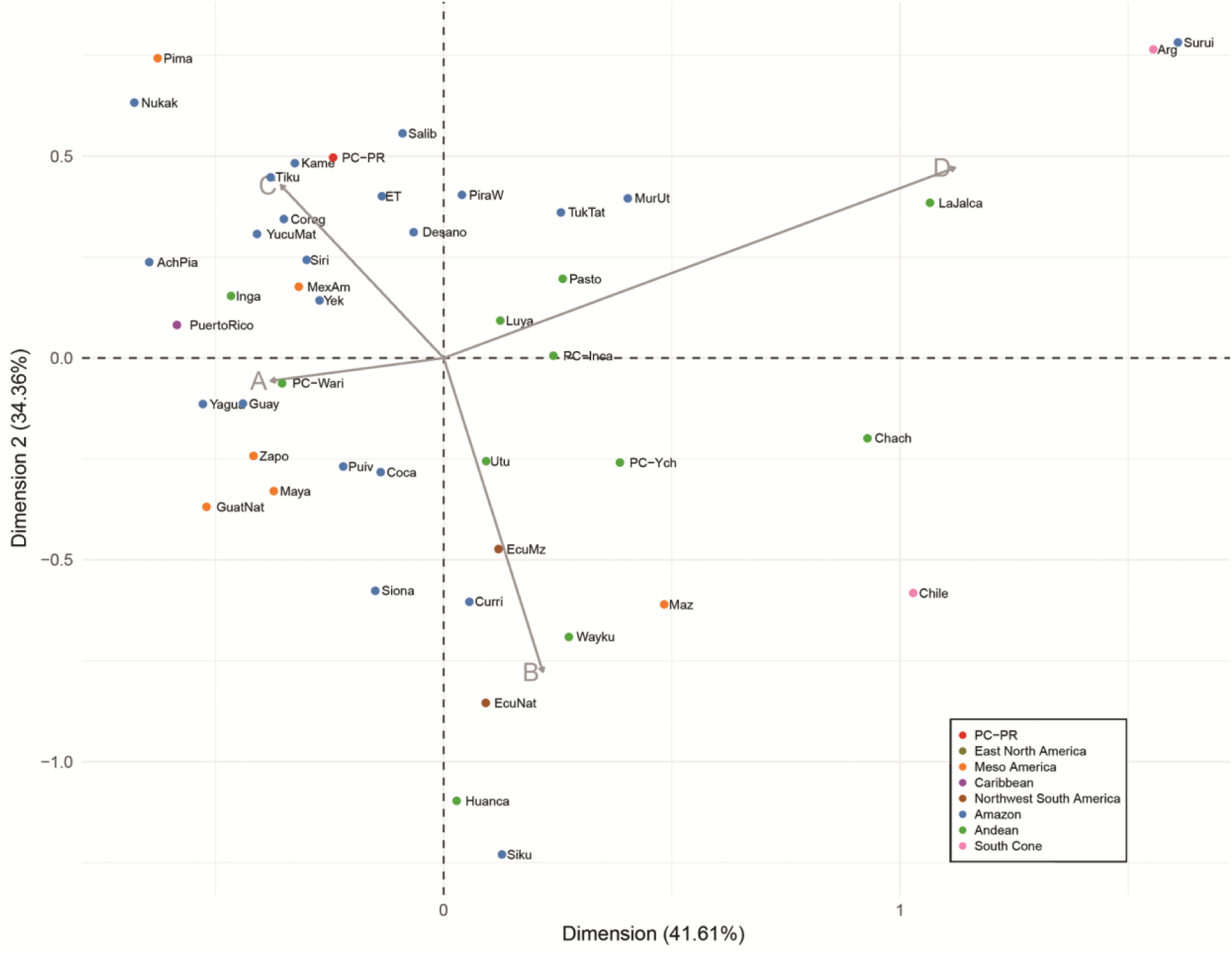
Correspondence analysis plot of haplogroup frequencies between PC-PR and 46 ancient and present-day populations from the Americas.

We repeated this analysis comparing PC-PR to eight Caribbean populations. The exact test found no significant difference in HVR-1 haplotype frequencies between PC-PR and most islanders, including Puerto Ricans, Cubans, Dominicans and the Trinidad First People’s Community (Table S15). However, pairwise comparisons of Φ_st_ measures calculated with direct haplotype sequences (Table S16) found the lowest distances were between PC-PR and PC-Guadeloupe, Puerto Ricans and the St. Vincent Garifuna (Native American lineages only) (Figure S12-S13; Table S15). When plotting two dimensions of non-metric MDS, PC-PR clusters closest to present-day Puerto Ricans (Figure 5). This pattern is repeated in the correspondence plot of haplogroup frequencies shown in Figure S14 where PC- PR falls closest to present-day Caribbean populations rather than other ancient Caribbean islanders. This clustering pattern likely stems from differences in haplogroup frequencies between pre-contact island groups. Specifically, we observe higher frequencies of haplogroups C1 and D1 in PC-Cuba and PC- Dominican Republic compared to PC-PR, PC-Guadeloupe and present-day islanders. Overall, these comparisons suggest that although Native American haplogroup frequencies are similar across Caribbean populations today, some differences may have existed between island communities in the past. Additionally, specific haplotypes differ between pre-contact and present-day populations, and among island groups.

**Figure 5.**
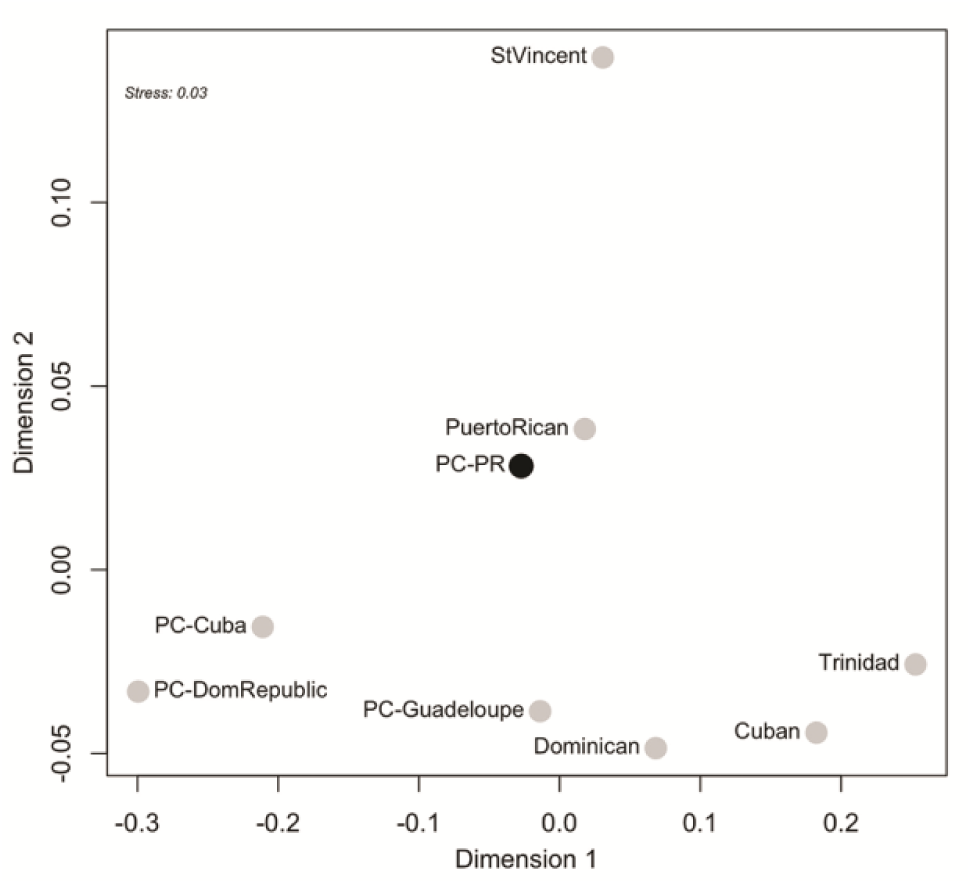
Non-metric MDS plot of HVR-1 Φ_st_ distances between PC-PR and eight ancient and present-day Caribbean populations.

### MtDNA network analysis

We constructed haplotype networks with complete mtDNA sequences from PC-PR, and reference populations from the Americas (Native American lineages only). We found that present-day Puerto Ricans were the only population that shared mtDNA haplotypes with PC-PR (Figure S15-S17). Thus, network analysis was repeated with only ancient and present-day Puerto Rican complete mtDNA haplotypes. We also conducted a second round of network analysis including HVR-1 haplotypes from the Caribbean. These networks suggest diverging histories for Caribbean A2 and C1 lineages.

The Puerto Rican complete mtDNA C1 network shows a large cluster of identical haplotypes found at high frequency in both the ancient and present-day populations (Figure 6). This clade is sub-haplogroup C1b2, the most common C1 lineage in both pre and post-contact Puerto Rico. C1b2 exhibits a star-like phylogenetic pattern, consistent with a history of lineage expansion and subsequent in-situ differentiation of derived haplotypes (Bandelt, et al. 1995). This pattern is mirrored in the HVR-1 Caribbean C1 network (Figure S18), although diversity is reduced, and previously distinct haplotypes are collapsed into a central C1 founder lineage. The C1 founder, and derived lineages, are found at high frequencies in most Caribbean populations (Lalueza-Fox, et al. 2001; Lalueza-Fox, et al. 2003; Mendizabal, et al. 2008; Martínez-Cruzado 2010; Vilar, et al. 2014; Benn-Torres, et al. 2015; Mendisco, et al. 2015). This pattern of reduced diversity is consistent with a strong founder effect of pre-contact C1 lineages in the peopling of the Caribbean islands and has been noted previously (Martínez-Cruzado 2010; Vilar, et al. 2014).

**Figure 6.**
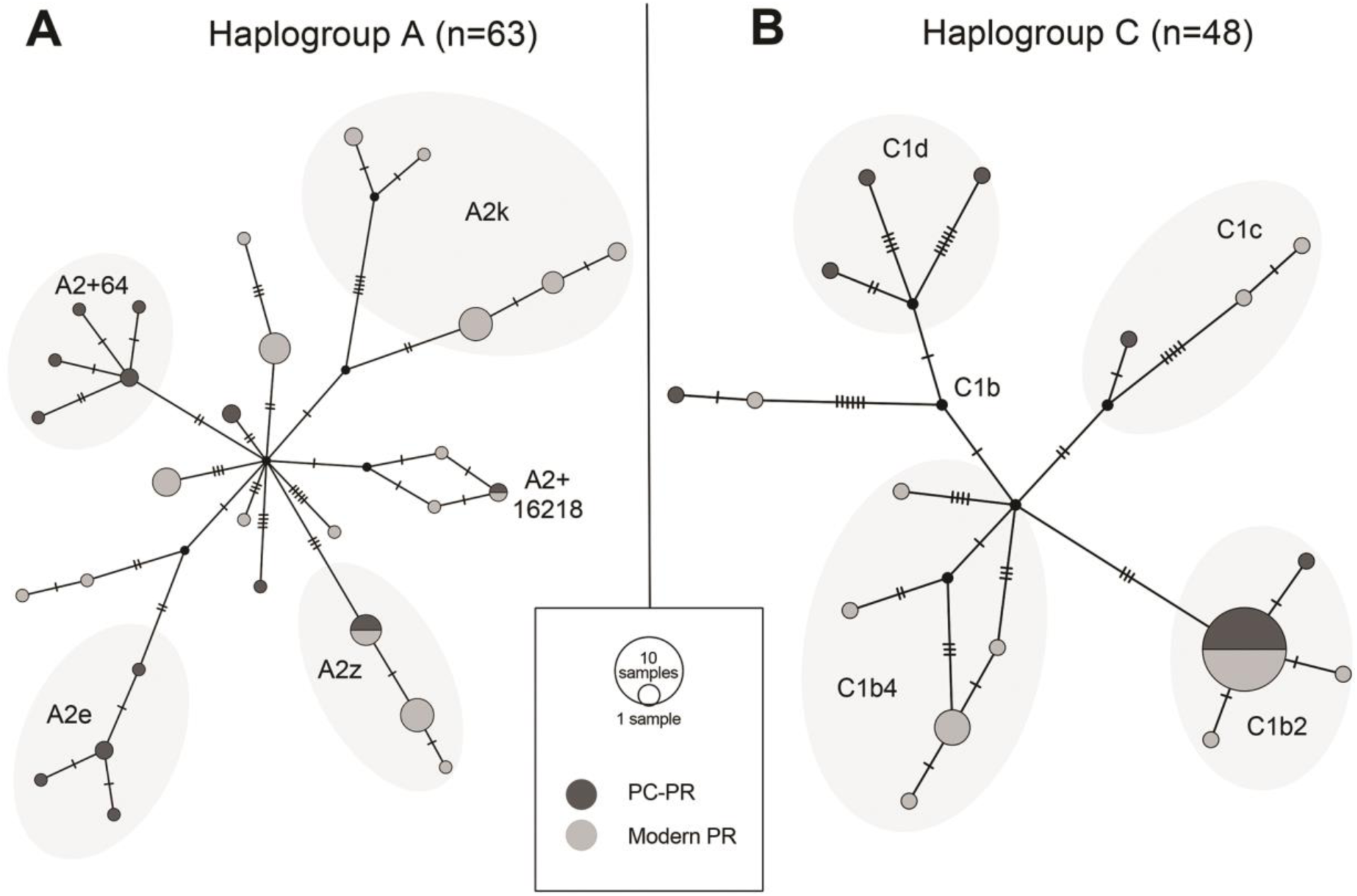
Median joining network of complete mtDNA haplotypes in PC-PR and present-day Puerto Rico. A) Haplogroup A, B) Haplogroup C. Major sub-haplogroup clades are labeled.

In contrast, the complete mtDNA Puerto Rican A2 network exhibits a diversity of low and mid-frequency haplotype clusters (Figure 6). Five sub-haplogroups (A2, A2+16218, A2+64, A2e, A2z) are represented in 14 PC-PR individuals. Two haplotypes belonging to sub-haplogroups A2+16218 and A2z are found at low frequencies in both PC-PR and present-day Puerto Ricans. Additionally, four HVR-1 haplotypes seen in PC-PR are also present in other Caribbean populations, including Cubans, Dominicans, and pre-contact individuals from Guadeloupe (Figure S18). However, many low frequency, derived A2 haplotypes have a restricted distribution, found on only one island or two neighboring islands. This topology is consistent with patterns noted in previous research that suggest multiple independent introductions of A2 lineages into Puerto Rico and the Caribbean, with subsequent expansion of some derived haplotypes across island populations (Martínez-Cruzado 2010; Vilar, et al. 2014).

Lastly, the HVR-1 network for haplogroup D1 demonstrates the high diversity of this clade despite its low frequency in the Antilles (e.g. six unique haplotypes in six PC-PR individuals) (Figure S18). This topology is inconsistent with expansion of a single founder and instead suggests derived D1 haplotypes may have also arrived independently to the Antilles. No complete mtDNA sequences matching PC-PR D1 haplotypes were found in comparative datasets, or in PhyloTree build 17 (van Oven and Kayser 2009).

### MtDNA demographic models and estimation of effective population size

The pre-contact demographic history of Puerto Rico and the Caribbean was further analyzed through an approximate Bayesian computation (ABC) approach. We used Bayesian Serial SimCoal (BayeSSC) to simulate samples based on the available HVR-1 data for pre-contact individuals from Cuba, Dominican Republic, Puerto Rico and Guadeloupe, under eight possible demographic scenarios (M1-M8 with several variants a,b,c, and d, as shown in Figure S19). In each scenario, we varied the amount and direction of migration and gene flow. The best goodness-of-fit value (AIC = 80.59) and highest relative likelihood value (ω=0.87) were associated with model M3b, which represented the peopling of the Antilles as an initial population split followed by low levels of inter-island migration in all directions (Table S17). All other models had low relative likelihood values, including those that varied the direction of gene flow. Therefore, we find that with the resolution provided by the available data, we cannot confidently estimate the direction or timing of pre-contact Caribbean migrations. We also used BayeSSC to test if neutral processes of drift and mutation sufficiently explain the observed genetic distances between pre and post-contact Native American mtDNA lineages in Puerto Rico. We simulated 10,000 mtDNA genome datasets with samples sizes matching the observed data under a model of population continuity (Reynolds, et al. 2015). Comparing our empirical genetic distances to the simulated distribution suggests we cannot reject the null hypothesis of population continuity over time (*p*=0.0804).

We estimated female effective population size (*Ne*) over time in Puerto Rico by constructing an extended Bayesian Skyline Plot (eBSP) in BEAST using 127 complete mtDNA sequences from ancient and present-day individuals as input (Native American lineages only) (Figure 7). The eBSP shows an increase in female *Ne* around 17,000 years before present (YBP) corresponding to population expansion associated with the settlement of the Americas (Mulligan, et al. 2008; Llamas, et al. 2016). After the peopling of Puerto Rico, approximately 5,000 YBP, *Ne* declines slowly until European contact, about 500 YBP, when a sharp reduction occurs. Similar contractions are seen in the demographic histories of other Indigenous populations impacted by European colonization (O’Fallon and Fehren-Schmitz 2011; Lindo, et al. 2016; Llamas, et al. 2016). We also detected some evidence of different demographic histories for the two major Puerto Rican mtDNA clades: A2 and C1 (Figure S20-S24). Specifically, we were unable to reject the null hypothesis of constant population size for the Puerto Rican C1 clade, but we detect a slight decline in *Ne* over time for haplogroup A2. Although these results suggest possible divergent evolutionary trajectories for these two clades, we note that the larger sample of available A2 sequences (A2: n=63, C1: n=47) likely results in increased power for detecting slight changes in Ne for clade A2 versus clade C1. The BEAST tree in Figure S21 shows an acceleration of the molecular rate of evolution for C1b2 compared to other Puerto Rican clades (Table S18-S19).

**Figure 7.**
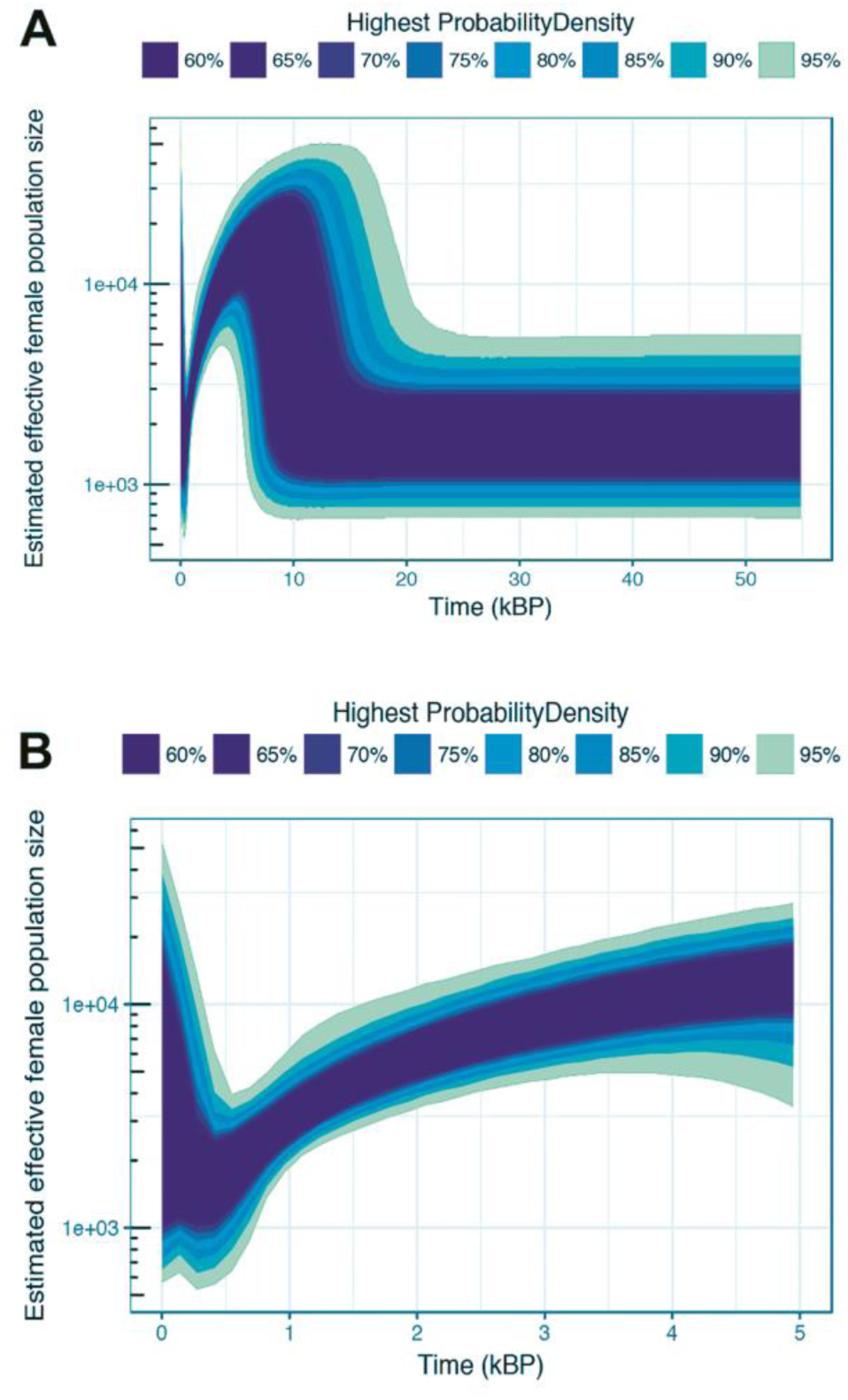
Extended Bayesian skyline plot (eBSP) of female effective population size, based on a generation time of 25 years. Bottom panel is zoomed in to 5,000 YBP.

### Autosomal genotypes

The two newly sequenced partial genomes recovered from Paso del Indio were analyzed alongside the genome of a pre-contact individual from the Preacher’s Cave site in the Bahamas: PC537, previously sequenced by Schroeder, et al. (2018). The ancient samples were intersected with a reference panel of 597,573 genome-wide SNPs collected from 967 present-day and ancient individuals in 37 worldwide populations (Table S20). Initial analyses conducted with all overlapping sites suggested the PC-PR individuals had a component of non-Native American ancestry (Figure S25-S26). However, this pattern disappeared after removing transitions, suggesting it was an artifact of post-mortem damage (Briggs, et al. 2007). At the best fit value of K=7, ADMIXTURE analysis conducted on the no-transitions dataset shows that the two PC-PR individuals have ancestry proportions similar to PC537, and to present-day Amazonian populations such as the Yukpa, Piapoco and Surui (Figure 8). Similarly, in the Principal Components Analysis (PCA), ancient Caribbean individuals cluster with South American populations from Amazonia and the Andes (Figure S27). These results echo the findings of our mtDNA analyses and suggest a close relationship between PC-PR communities and Indigenous South American populations.

**Figure 8.**
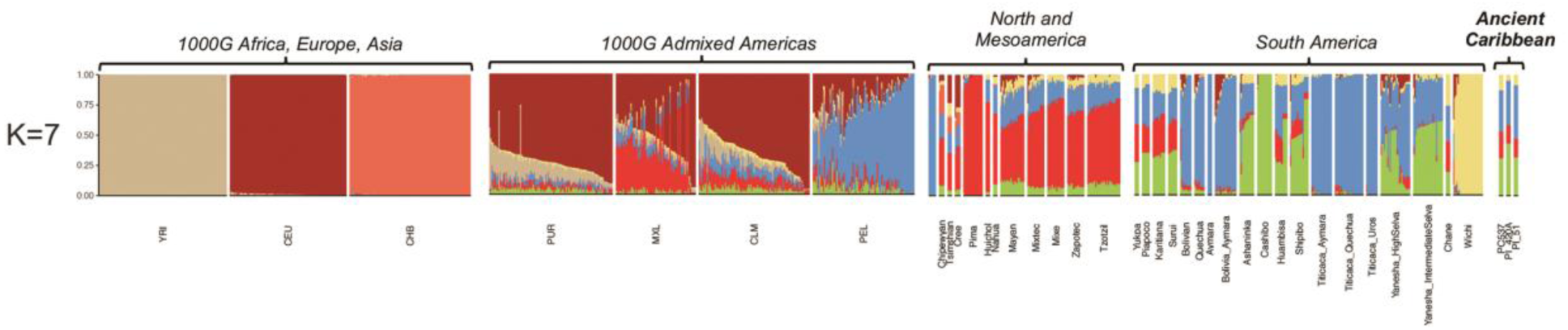
ADMIXTURE analysis at K=7 including autosomal genotypes from two pre-contact individuals from PC-PR (PI-420A, PI-51), one pre-contact individual from the Bahamas (PC537), and 37 worldwide reference population.

To determine the genome-wide affinities of these individuals, Outgroup *f*_3_ statistics were calculated in the form *f*_3_(ancient, X; Yoruba) (Raghavan, et al. 2014), where ‘ancient’ represented one of the two PC-PR ancient genomes and X was a set of 22 Americas populations or the PC537 individual (Figure S28). However, results were inconsistent and affected by the low number of overlapping sites recovered in the PC-PR samples (Table S21). In our first analysis including all overlapping positions, the two PC-PR individuals were closest first to PC537, and second to South American populations. But, upon removing transitions the closest similarities were between PC-PR and both South American and Mesoamerican groups. Thus, with the low resolution provided by the available data, we cannot confidently estimate genome-wide affinities. Additional analysis with higher coverage genomes is necessary to draw further conclusions about the autosomal ancestry and affinities of PC-PR communities.

## Discussion

### Preservation and ancient DNA recovery in Puerto Rican archaeological sites

Endogenous aDNA was poorly preserved in the samples included in this study, consistent with previous Caribbean paleogenomics research (Schroeder, et al. 2015; Nieves-Colón, et al. 2018; Schroeder, et al. 2018). Shotgun sequencing demonstrated that endogenous DNA was found at low quantities in the PC-PR sample. But, through target enrichment, we successfully recovered medium to high coverage mtDNA genomes and low coverage partial autosomal genomes. Thus, our findings show that target enrichment and high-throughput sequencing are essential approaches for maximizing aDNA recovery in Caribbean and tropical archaeological contexts. We also found that DNA preservation varied across sites, with Tibes having the worst preservation. This is consistent with previous reports of poor organic preservation at Tibes relative to other sites in Puerto Rico (Pestle and Colvard 2012). Our results suggest that site-specific processes may play a larger part in aDNA decay than island or region-wide environmental conditions. Future human paleogenomics research at Tibes may benefit from assessing endogenous aDNA preservation in dense skeletal tissues, such as the petrous bone (Gamba, et al. 2014).

### MtDNA diversity in pre-contact Puerto Rico communities

Genetic relationships at an intra-island scale were evaluated by testing for significant differences in mtDNA diversity between individuals from the three studied sites. Archaeological evidence indicates a trend towards cultural differentiation and regionalization across Caribbean communities during the LCA. Within Puerto Rico, this is visible in distinctive material culture, settlement patterns and ceramic traditions that differentiated communities along an East-West divide after A.D. 600 (Rouse 1992; Curet, et al. 2004). In this study, we found no evidence for genetic structure or significant differences in mtDNA diversity between PC-PR communities. This suggests that cultural diversification during the LCA may not have been accompanied by genetic isolation or inter-site restrictions on female-mediated gene flow. This interpretation contrasts with (Martínez-Cruzado, et al. 2005) who reported a geographic gradient in the distribution of C1 mtDNA lineages among present-day Puerto Ricans. However, our results are consistent with subsequent work conducted with higher-resolution markers (Gravel, et al. 2013; Vilar, et al. 2014). Thus, we infer that biogeographic differentiation patterns in the Native American ancestry of present-day Puerto Ricans do not date to the pre-contact period, but arose later, due to recent demographic processes. However, we acknowledge that these results may be biased by small sample sizes per site (especially for Tibes) and that future research may reveal more complex patterns of intra-island diversity.

Archaeological and isotopic evidence indicate that inter-island interaction and mobility increased in the Antilles during the LCA (Laffoon and Hoogland 2012; Mol 2013). But, dental biodistance studies found few shared morphological traits among pre-contact island groups, suggesting that little gene flow occurred between them (Coppa, et al. 2008). Here, we find genetic evidence for both migration and contact, as well as isolation and differentiation in the population history of the ancient Antilles. With the available HVR-1 data, we identified a shared mtDNA component among pre-contact groups, mainly represented by high frequencies of shared C1 haplotypes. But, we also observed several private or island-specific mtDNA haplotypes, which differentiate groups in inter-island comparisons. We further found larger differences in haplogroup frequencies between PC-PR and several pre-contact island groups than between PC-PR and present-day islanders. For example, while PC-PR communities carried high frequencies of A2 and C1 mtDNA lineages, ancient communities from Cuba and Dominican Republic had high frequencies of C1 and D1 (Lalueza-Fox, et al. 2001; Lalueza-Fox, et al. 2003). This distribution contrasts with mtDNA haplogroup patterns reported for PC-Guadeloupe and for most present-day islanders, including Cubans and Dominicans (Martínez-Cruzado, et al. 2005; Benn-Torres, et al. 2007; Mendizabal, et al. 2008; Vilar, et al. 2014; Benn-Torres, et al. 2015; Schurr, et al. 2016). Differences in mtDNA diversity patterns between the Greater and Lesser Antilles have been reported previously and may reflect distinct island founder populations or the effects of genetic drift and relative isolation between ancient island communities (Vilar, et al. 2014; Benn-Torres, et al. 2015). We find support for the latter scenario through our best fit demographic model in which pre-contact island populations diverge from each other after initial settlement with limited subsequent gene flow. These findings suggest that female-mediated gene flow and matrilineal kinship was not essential for the maintenance of Pan-Caribbean interaction networks during the CA and LCA. Therefore, inter-island connectivity and mobility patterns may respond primarily to other factors such as trade, residency patterns or patrilineal kin networks (Keegan and Maclachlan 1989; Laffoon and Hoogland 2012; Mol 2013). Future research with a more comprehensive genomic sampling of the pre-contact Antilles is needed to study this in more detail.

### MtDNA lineages and the origins of Ceramic Age Caribbean populations

MtDNA haplogroup distribution in PC-PR fits a broader Caribbean-wide pattern of high frequencies of haplogroups A2 and C1 and low frequencies of D1 (Martínez-Cruzado, et al. 2005; Benn-Torres, et al. 2007; Mendizabal, et al. 2008; Vilar, et al. 2014; Benn-Torres, et al. 2015; Schurr, et al. 2016). Similar distributions in several ancient Caribbean populations suggest this pattern originated in the pre-contact era (Lalueza-Fox, et al. 2003; Mendisco, et al. 2015). We did not find haplogroup B2 in the PC-PR sample. This is consistent with previous research which suggests that B2 was rare in the pre-contact period and that most B2 lineages in present-day Puerto Rico arrived during the final centuries of the LCA or after European contact (Martínez-Cruzado 2010; Vilar, et al. 2014; Benn-Torres, et al. 2015; Mendisco, et al. 2015). However, PC537, the pre-contact individual from the Bahamas, carried a B2 haplotype (Schroeder, et al. 2018), so this pattern may not extend to other Antilles.

We found similarities in mtDNA variation and haplotype frequencies between PC-PR and Indigenous populations from northwest Amazonia and the Andes. Specifically, the closest relationships were observed between PC-PR and Eastern Tukanoan speakers from the regions surrounding the Orinoco and Rio Negro rivers. These include groups such as the Siriano, Desano and Wanano. We also found close similarities between PC-PR, Yekuana and Kamentsa groups living in Venezuela and Colombia; as well as between PC-PR, Pasto and Quechua speakers from the Andean foothills of Colombia and northern Peru (Barbieri, et al. 2011; Lee and Merriwether 2015; Arias, et al. 2018). These findings are consistent with previous genetics research (Lalueza-Fox, et al. 2001; Lalueza-Fox, et al. 2003; Martínez-Cruzado, et al. 2005; Martínez-Cruzado 2010; Gravel, et al. 2013; Moreno-Estrada, et al. 2013; Vilar, et al. 2014; Schroeder, et al. 2018), biodistance studies (Ross 2004; Coppa, et al. 2008) and archaeological evidence (Rouse 1992; Chanlatte Baik 2003; Rodríguez Ramos, et al. 2013) all of which suggest that Caribbean CA populations originated in the Orinoco River delta region of northern South America.

The high frequency of sub-haplogroup C1b2 we found in PC-PR (33%) also supports a strong genetic link with South America. Coalescent analyses indicate C1b2 arose in that continent approximately 2,000 YBP (Perego, et al. 2010; Gomez-Carballa, et al. 2015). It has been identified in Amazonian populations including the Yanomamö, Kraho, Marikitare and Pasto from Colombia, Brazil and Venezuela, and in communities of Indigenous-descent from Uruguay (Torroni, et al. 1993; Williams, et al. 2002; Noguera-Santamaría, et al. 2015; Sans, et al. 2015). C1b2 is also the most frequent C1 lineage found in present-day Puerto Ricans (Martínez-Cruzado 2010; Vilar, et al. 2014). Previous research estimated that it likely arrived in Puerto Rico after the CA expansions, between 647 ± 373 YBP (estimated with mtDNA control region sequences) and 1,195 ± 690 YBP (estimated with HVR-1 sequences) (Martínez-Cruzado 2010; Vilar, et al. 2014). In our network analysis, the Puerto Rican C1b2 clade exhibits a star-like signature suggestive of a strong founder effect with subsequent expansion and differentiation. This is also supported by the BEAST tree for haplogroup C1, which shows that the Puerto Rican C1b2 clade has an abnormally high rate of evolution relative to other clades. Given the large number of identical C1b2 haplotypes we identified, and its consistently high frequency over time, we infer that this lineage has had a continuous presence on the island from the pre-contact period until the present-day. HVR-1 polymorphisms typed in most other Caribbean populations lack sufficient resolution to distinguish C1b2 from the C1 New World founder lineage. Thus, we cannot conclusively determine its frequency over time in other islands. However, C1 and C1b haplotypes are found at high frequency throughout the region (Lalueza-Fox, et al. 2001; Lalueza-Fox, et al. 2003; Mendizabal, et al. 2008; Benn-Torres, et al. 2015; Mendisco, et al. 2015; Schurr, et al. 2016). Taken together, these data suggest that the Caribbean C1b2 clade arrived originally from South America during the expansion of Arawak speaking populations into the Antilles. We thus support the conclusions of previous research and designate this clade as a characteristic lineage of CA Antillean populations (Martínez-Cruzado 2010; Vilar, et al. 2014).

Haplogroup A2 was the second most common haplogroup in our sample, accounting for 40% of all PC-PR sequences. In contrast with the patterns observed for lineage C1b2, the topology of haplotype A2 networks suggest multiple, independent lineage introductions into Puerto Rico and other Antilles during the pre-contact period. Previous research proposed that some of these lineages may have originated in Mesoamerica and arrived in the Caribbean during the Lithic Age (Martínez-Cruzado 2002, 2010; Vilar, et al. 2014). This hypothesis is supported by similarities in material culture between Lithic Age Caribbean groups and coeval populations in Belize and Honduras (Wilson and Hester 1998). Additionally, Mesoamerican populations have the highest frequency and diversity of A2 haplotypes (Perego, et al. 2010; Gonzalez-Martin, et al. 2015) and similarities have been identified between Caribbean A2 HVR-1 types and continental lineages (Vilar, et al. 2014). However, none of the A2 haplotypes in PC-PR were represented in a reference database of over 1,600 complete mtDNA genomes from the Americas, or in broad surveys of Mesoamerican mtDNA diversity (Kumar, et al. 2011; Gorostiza, et al. 2012; Perego, et al. 2012; Mizuno, et al. 2014; Gonzalez-Martin, et al. 2015; Söchtig, et al. 2015). Thus, we cannot trace a direct genetic link between PC-PR and Mesoamerican populations.

Lastly, we find that some mtDNA lineages in the PC-PR sample, are not found outside of the Antilles and as such may represent locally differentiated variation. For instance, three PC-PR individuals had haplotypes belonging to sub-haplogroups A2+16218 and A2z. A2+16218 was first reported in self-identified communities of Indigenous descent from Indiera Alta, Maricao, Puerto Rico (Martínez-Cruzado, et al. 2001; Martínez-Cruzado 2010). Afterwards, it was found in broad samples of Cubans and Puerto Ricans (Mendizabal, et al. 2008; Vilar, et al. 2014) and in one individual from Grande Anse, a pre-contact site in Guadeloupe (Mendisco, et al. 2015). Martinez-Cruzado (2010) proposed that A2+16218 could be a derived Caribbean-specific lineage, with a proximate origin in pre-contact communities from Mona Island, an island off the southwestern coast of Puerto Rico. Similarly, A2z may also be a Caribbean lineage, with a possible origin in Cuba, where it is found at high frequencies today (Vilar, et al. 2014).

That said, genetic drift and the population bottlenecks caused by European contact have led to loss of mtDNA diversity in Native American populations. Genetic discontinuity between pre-contact populations and their descendants has been observed previously in aDNA studies in the Americas (Lindo, et al. 2016; Llamas, et al. 2016). Additionally, some sub-haplogroups identified in PC-PR, such as C1d, C1c, D1 and basal A2 types, are New World founding lineages and have a Pan-American distribution (Perego, et al. 2010; Kumar, et al. 2011). These lineages are not informative for tracing sub-continental origins within the Americas. Thus, with the available data, we cannot exclude potential genetic contributions from Mesoamerica or other regions of the Americas to the ancestry of pre-contact Puerto Rican communities.

### MtDNA diversity and effective population size

MtDNA diversity in PC-PR is low relative to most comparative populations; with the exception of several Amazonian groups such as the Surui and Karitiana. Low genetic diversity has previously been reported for Amazonian and Eastern South American Indigenous groups due to small historical effective population sizes, isolation and repeated genetic bottlenecks (Lewis, et al. 2007; Hunley, et al. 2016). Similarly, low diversity in PC-PR communities may stem from genetic drift and serial founder effects during the original peopling of the Americas, the initial peopling of the Antilles or the CA expansions. Previous studies of mtDNA and autosomal loci in Puerto Ricans noted a pattern consistent with strong effects of drift and at least one pre-contact population bottleneck (Martínez-Cruzado, et al. 2005; Gravel, et al. 2013). Estimates of long-term historical *Ne* gleaned from autosomal genome fragments suggest that the Native American ancestors of present-day Puerto Ricans had an effective breeding population size of approximately 1,922 individuals; 32 times smaller than the estimated size of coeval populations in Mexico (Gravel, et al. 2013). Strong effects of genetic drift have also been described in previous studies of mtDNA diversity in ancient and present-day Cuban and Dominican populations (Lalueza-Fox, et al. 2001; Lalueza-Fox, et al. 2003; Mendizabal, et al. 2008). In contrast, based on the ancient genome of one individual, Schroeder, et al. (2018) estimated a relatively high *Ne* of around 1,600 breeding individuals for pre-contact communities in the Bahamas. The results of our diversity estimates, haplotype network and eBSP analyses are consistent with low mtDNA *Ne* for PC-PR. Our findings support a scenario where a reduced number of mtDNA haplotypes became isolated from their ancestral population after settling in Puerto Rico. This led to a reduction in female *Ne*, increased susceptibility to genetic drift and loss of mtDNA diversity over time, with the largest reduction occurring after European contact.

### Autosomal ancestry of PC-PR communities

Here we report the first recovery of human autosomal genomes from pre-contact Puerto Rico. Our analyses of these data show similar ancestry patterns between two PC-PR individuals and PC537, a pre-contact individual from the Bahamas. Schroeder, et al. (2018) found the genome-wide ancestry of PC537 to be closely related to present-day Arawakan speakers from the Amazon and Orinoco regions of Northern South America, in agreement with archaeological and genetic evidence. The results of our ADMIXTURE analysis are broadly consistent with these findings as we identify similar ancestry proportions between PC-PR individuals and Indigenous populations from the Colombian, Venezuelan and Brazilian Amazon, including the Yukpa, Piapoco, Karitiana and Surui. These similarities suggest that pre-contact Caribbean populations had a shared component of autosomal ancestry with close affinity to South American populations. However, due to poor aDNA preservation and limited overlap between our samples and available reference panels, our genome-wide data lack resolution for confident estimation of allele frequency-based statistics such as Outgroup *f*_3_, and for direct estimation of sub-continental origins. Thus, we refrain from drawing further inferences regarding the autosomal ancestries or genome-wide affinities of PC-PR communities. Future research with higher coverage genomes and finer-scale analyses of genetic structure is necessary to further address this question.

### Indigenous genetic legacies in Puerto Rico

Through demographic modeling and lineage sharing analysis, we find evidence of direct mtDNA ancestry and partial genetic continuity between pre-contact Indigenous communities and present-day Puerto Ricans. Specifically, three mtDNA haplotypes from PC-PR persist in the modern population. Martínez-Cruzado (2010) predicted that nine Native American mtDNA lineages found in Puerto Ricans had their proximate origin in pre-contact Caribbean populations. We find three of these lineages in PC-PR: C1b2, A2z, A2+16218. However, we also identified multiple haplotypes that are not shared between pre and post contact Puerto Rico. This differentiation reflects both the neutral processes of drift and lineage loss, and the demographic shifts brought by European contact.

Extensive historical evidence documents the decline of Indigenous Caribbean populations during the first decades of European colonization. Just before contact, in the 15^th^ century, estimates place the population of Puerto Rico between 30,000 to 70,000 individuals (Anderson-Córdova 2005; Anderson-Córdova 2010). By the census of 1530, the native population was reportedly 1,543 individuals. This count included native communities as well as Indigenous people forcibly relocated from other islands or from the Circum-Caribbean basin (Anderson-Córdova 2005; Wilson 2007). Despite this decline, in a database of over 1,600 complete mitochondrial genomes from the Americas, the only identical matches to PC-PR haplotypes were found in Puerto Rico. This indicates that admixed Puerto Rican genomes are at least a partial reservoir of pre-contact mtDNA diversity. As publicly available genomic datasets become more representative of the diversity of Native American and Latino populations (Bustamante, et al. 2011), it is possible we may find other populations, in the Caribbean or elsewhere, that are also closely related to pre-contact Puerto Rican communities. Beyond its anthropological importance, better characterization of the genetic diversity of ancient populations that contributed to present-day Puerto Rican ancestry may guide future efforts at rare variant discovery in Caribbean biomedical cohorts (Belbin, et al. 2017).

Our findings do not support historical narratives of complete population replacement or genetic extinction of Indigenous communities in Puerto Rico. Indigenous heritage is an important component of Puerto Rican national and ethnic identity (Haslip-Viera 2001; Veran 2003). However, cultural claims of indigeneity and Native American ancestry coexist with narratives of Indigenous extinction (Benn-Torres 2014). For the most part, these narratives are rooted in interpretations of the written historical record and colonial era censuses. But these documents under-represent the number of Indigenous people living in Puerto Rico and other islands during Spanish occupation (Anderson-Córdova 2010; Benn-Torres 2014). Other historical sources as well as ethnographic research note that Indigenous peoples in Puerto Rico and the Antilles resisted European colonization and persisted, albeit in small numbers, well into the 16^th^ century (Anderson-Córdova 2010). Moreover, oral histories of Indigenous survival are common in Puerto Rico and other islands (Forte 2006). Some of these oral narratives are reinforced by genetic research which has found reservoirs of Native American genetic ancestry in communities that self-identify with Indigenous or Maroon descent in Puerto Rico, Jamaica and other islands (Martínez-Cruzado, et al. 2001; Madrilejo, et al. 2015; Schurr, et al. 2016; Fuller and Benn Torres 2018; Benn Torres, et al. 2019). Furthermore, self-identified and government-recognized Indigenous communities are found throughout the Caribbean and the international Caribbean diaspora (Forte 2006).

## Conclusions

In conclusion, this research characterized the genetic diversity of pre-contact communities in Puerto Rico and tested hypotheses about their origins and relationships to other Native American and Caribbean populations. Our findings support a primarily South American contribution to the genetic ancestry of pre-contact Puerto Rican peoples, in agreement with previous genetics and archaeological research. However, we cannot reject the possibility that additional migrations from other parts of the Americas also contributed to the peopling of Puerto Rico. Future research with more ancient genomes from the Antilles, and higher coverage genome-wide data will provide added resolution for detecting ancient admixture events in the Caribbean and elucidating the genetic relationships between island communities and continental Native American populations. We also found evidence that at least some of the mtDNA diversity of present-day Puerto Ricans, can be directly traced to pre-contact Puerto Rican communities. Thus, our study adds to a growing corpus of research documenting the persistence of cultural and biological elements from pre-contact Indigenous Caribbean peoples into the present day. We hope our findings lead to a critical and interdisciplinary reassessment of historical narratives of Indigenous extinction in Puerto Rico, while informing future study of Indigenous responses to European colonization, and of the complex role of native peoples in shaping the biocultural diversity of the Antilles.

## Materials and Methods

### Sampling, DNA extraction and library preparation

The human skeletal remains included in this study are patrimony of the people of the Commonwealth of Puerto Rico. Permits for destructive sampling and DNA analysis were obtained from three government agencies in Puerto Rico (Figure S1). We sampled 124 individuals excavated from three pre-contact sites: Punta Candelero (n=34), Tibes (n=46) and Paso del Indio (n=44) (Figure 1; Figure S2). Direct radiocarbon dates for 81 individuals were previously obtained by (Pestle 2010) (Table S1; Figure S3). DNA was extracted from tooth roots, bone or dental calculus using silica-based extraction methods optimized for aDNA (Table S2) (Rohland and Hofreiter 2007; Dabney, et al. 2013; Nieves-Colón, et al. 2018). DNA libraries were constructed, double-indexed and amplified following (Meyer and Kircher 2010; Kircher, et al. 2012; Seguin-Orlando, et al. 2015). The optimal number of PCR cycles was determined by real-time PCR (qPCR). Libraries were purified with the Qiagen® MinElute PCR kit. DNA fragment sizes were assessed with the Agilent 2100 Bioanalyzer.

### Target enrichment and Illumina sequencing

Targeted enrichment for the complete mtDNA genome was performed following (Maricic, et al. 2010; Ozga, et al. 2016). MtDNA enriched libraries were sequenced on multiple runs of the Illumina MiSeq. Thirty-five libraries were additionally screened by shotgun sequencing on several runs of the Illumina NextSeq 500 and HiSeq 2500. Twenty-two of these libraries were selected for WGE performed following (Carpenter, et al. 2013), with slight modifications (see Supplementary Materials & Methods) or with the MYbaits Human Whole Genome Capture Kit (Arbor Biosciences), following manufacturer’s instructions. After WGE, libraries were amplified, and purified as detailed above, then sequenced on several runs of the Illumina NextSeq 500 and HiSeq 2500.

### Sequence read processing

Illumina sequence reads were merged, and adapters trimmed with SeqPrep (https://github.com/jstjohn/SeqPrep). Mapping was performed using BWA v.0.7.5 with seed disabled (Li and Durbin 2009; Schubert, et al. 2012). For mtDNA enriched libraries, reads were mapped to the revised Cambridge Reference Sequence (rCRS) (Andrews, et al. 1999). For shotgun and WGE libraries, reads were mapped to the GRCh37 (hg19) assembly with the mitochondrial sequence replaced by the rCRS. BAM files from samples sequenced over multiple sequencing runs were merged with SAMtools v.0.1.19 (Li, et al. 2009). Filtering and duplicate removal were also performed with SAMtools, keeping reads with quality ≥Q30 and no multiple mappings. Damage patterns were characterized and read quality scores rescaled with mapDamage v.2.0.2 (Figure S5-S7) (Jónsson, et al. 2013). Contamination estimates were generated for mtDNA reads with contamMix (Fu, et al. 2013) and schmutzi (Renaud, et al. 2015). Reads were further contamination filtered with PMDtools (Skoglund, et al. 2014) (Tables S3-S4).

### MtDNA variant calling, haplogroup assignment and data analyses

MtDNA variants were called using SAMtools *mpileup* on the rescaled BAM files. Haplogroup assignment was performed in HaploGrep 2.0 (Weissensteiner, et al. 2016) and confirmed manually with reference to Phylotree mtDNA tree Build 17 (van Oven and Kayser 2009). MtDNA consensus sequences were generated with schmutzi and curated manually in Geneious v.7.0.6 (Biomatters). MtDNA reference data collected from the literature included 1,636 complete mtDNA genomes from ancient and present-day individuals from the Americas and Caribbean. For comparative analyses, this dataset was restricted to 1,403 sequences grouped into 47 populations. A second comparative dataset included 391 mtDNA HVR-1 sequences from ancient and present-day Caribbean islanders (see Table S10).

We performed a Mantel test to evaluate the relationship between temporal and genetic distance in the PC-PR sample. For radiocarbon dated individuals (n=35), this test compared a Euclidean distance matrix of median calibrated radiocarbon dates and a Tamura-Nei genetic distance matrix. Intra-population diversity measures such as number of haplotypes (*h),* number of segregating sites (*S*), nucleotide (π) and haplotype diversity (*Hd*) were calculated for PC-PR and all reference populations. We calculated exact tests of genetic differentiation to compare haplotype frequencies between the three PC-PR sites, between PC-PR and continental Native American populations (complete mtDNA) and between PC-PR and Caribbean populations (HVR-1). We also calculated pairwise population Φ_st_ measures using complete mtDNA and HVR-1 sequences (Excoffier, et al. 1992). The resulting matrix was used as input for non-metric MDS scaling. Population haplogroup frequencies (e.g. A, B, C, D, X) were estimated by direct counting and used as input for correspondence analysis. These analyses were performed and plotted using R v. 3.6.1 (R Core Team). For additional details see Supplementary Materials & Methods. Lastly, median joining haplotype networks were constructed in popART with default parameters (Bandelt, et al. 1999; Leigh and Bryant 2015). Complete mtDNA and HVR-1 networks were constructed per haplogroup (A2, C1, D1) comparing PC-PR with populations from the Americas and Caribbean.

We used BayeSSC (Excoffier, et al. 2000; Anderson, et al. 2005) to model eight possible demographic scenarios (Figure S19: M1-M8) that could explain the HVR-1 mtDNA diversity reported for ancient Caribbean populations (Lalueza-Fox, et al. 2001; Lalueza-Fox, et al. 2003; Mendisco, et al. 2015). To determine which model most likely explained the observed data, we calculated Euclidean distances between simulated and empirical datasets following (Beaumont, et al. 2002; Duggan, et al. 2017). Goodness of fit was determined using the Akaike information criterion (AIC) (Akaike 1974). The model with the lowest AIC value and the highest relative likelihood value was chosen as the best fit. We also used BayeSSC simulations to evaluate if the observed *Fst* genetic distances between the complete mtDNA genomes collected from ancient and present-day Puerto Rico (Native American haplotypes only) were consistent with a model of population continuity. This model was simulated 10,000 times to produce a distribution of possible *Fst* values for comparison with observed *Fst* values (Reynolds, et al. 2015).

Demographic history was further reconstructed with eBSP in BEAST 1.8.4 (Drummond, et al. 2012) using partitioned complete mtDNA sequences from ancient and present-day Puerto Rico as input. PartitionFinder v.1.1.1 (Lanfear, et al. 2012) was used to determine the best partitioning scheme and substitution model for the data. Median calibrated radiocarbon date estimates were used as tip calibrations to reconstruct the phylogeny. Individuals without radiocarbon dates were assigned a prior date range based on archaeological context, and posterior dates were estimated based on the empirically calculated molecular rate (Table S19). Analyses were conducted under a strict clock model, which could not be rejected after testing other models. A total of 3 chains of 100 million generations were executed for three datasets: All sequences, Haplogroup A, and Haplogroup C. Parameters were sampled every 10,000 generations, with the initial samples discarded as burn-in. Generation time was set as 25 years.

### Autosomal genome genotype calling and data analysis

Two PC-PR samples with the highest read depth and endogenous content (PI-420a and PI-51) were sequenced across multiple runs to increase genome coverage. Reads were combined, and filtering was repeated as detailed above. Chromosomal sex was estimated following (Skoglund, et al. 2013) and X-chromosome contamination was estimated with ANGSD (Korneliussen, et al. 2014) (Table S5). Autosomal genotypes from PC-PR were analyzed alongside the genome of a pre-contact individual from the Bahamas: PC537 (Schroeder, et al. 2018). These samples were intersected with a reference panel of 597,573 SNPs from 37 worldwide populations compiled from the literature (N=967), which included 656 individuals from the Americas (see Table S20). Haploid genotype calls in the ancient Caribbean individuals were generated by randomly sampling one read per overlapping positions with the reference panel. PCA was performed with Eigensoft 6.0.1 (Patterson, et al. 2006) using the *lsqproject* option. Outgroup-*f*_3_ analysis was performed with *qp3pop* within Admixtools (Patterson, et al. 2012) in the form *f*_3_(ancient, X; Yoruba) (Raghavan, et al. 2014), where ‘ancient’ represented one of the ancient genomes from PC-PR and X was a reference population or individual. For admixture analysis, a genotype likelihood approach was implemented with FastNGSadmix (Jørsboe, et al. 2017). Genotype likelihoods in the ancient samples were estimated for all reference panel overlapping positions using ANGSD. We then conducted ten runs of ADMIXTURE for each value of K3 to K7 (Alexander, et al. 2009) using only the reference panel individuals. We retained the run with the highest likelihood per K to calculate the proportion of the ancestral components in the ancient genomes (Figure S26). Analyses were performed both with and without transitions to account for aDNA damage.

## Acknowledgements

We thank L. Antonio Curet, Alexandra Adams, Meredith Carpenter, Rosa Fregel, Kelly Blevins, the Crabbe family, Irma Zayas and the staff of the *Centro Ceremonial Indígena de Tibes* for support and assistance. This work was supported by the National Science Foundation (BCS-1622479 to M.N.C. and BCS-0612727 to W.J.P.), the Rust Family Foundation Grant for Archaeological Research (RFF-2016-08 to M.N.C.), Sigma Xi (G2012161222 and G201510151642390 to M.N.C.) and pilot grant programs from the ASU School of Human Evolution and Social Change, School of International Letters and Cultures and Graduate and Professional Student Association. Sequence data generated through this study are available in the NCBI Short Read Archive (SRA) under BioProject accession PRJNA557308.

## References

Akaike H. 1974. A new look at the statistical model identification. IEEE T Automat Contr 19:716–723.

Alexander DH, Novembre J, Lange K. 2009. Fast model-based estimation of ancestry in unrelated individuals. Genome Res 19:1655–1664.

Anderson CN, Ramakrishnan U, Chan YL, Hadly EA. 2005. Serial SimCoal: a population genetics model for data from multiple populations and points in time. Bioinformatics 21:1733–1734.

Anderson-Córdova K. 2005. The Aftermath of Conquest: The Indians of Puerto Rico during the Early Sixteenth Century. In: Siegel P, editor. Ancient Borinquen: Archaeology and ethnohistory of native Puerto Rico. Tuscaloosa: University of Alabama Press. p. 337–352

Anderson-Córdova K. 2010. Surviving Spanish Conquest: Indian Fight, Flight, and Cultural Transformation in Hispaniola and Puerto Rico. Tuscaloosa: University of Alabama Press.

Andrews RM, Kubacka I, Chinnery PF, Lightowlers RN, Turnbull DM, Howell N. 1999. Reanalysis and revision of the Cambridge reference sequence for human mitochondrial DNA. Nat Genet 23:147–147.

Arias L, Barbieri C, Barreto G, Stoneking M, Pakendorf B. 2018. High-resolution mitochondrial DNA analysis sheds light on human diversity, cultural interactions, and population mobility in Northwestern Amazonia. Am J Phys Anthropol 165:238–255.

Bandelt H-J, Forster P, Rohl A. 1999. Median-Joining Networks for Inferring Intraspecific Phylogenies. Mol Biol Evol 16:37–48.

Bandelt H-J, Forster P, Sykes BC, Richards MB. 1995. Mitochondrial portraits of human populations using median networks. Genetics 141:743–753.

Barbieri C, Heggarty P, Castri L, Luiselli D, Pettener D. 2011. Mitochondrial DNA variability in the Titicaca basin: Matches and mismatches with linguistics and ethnohistory. Am J Hum Biol 23:89–99.

Beaumont MA, Zhang W, Balding DJ. 2002. Approximate Bayesian computation in population genetics. Genetics 162:2025–2035.

Belbin GM, Odgis J, Sorokin EP, Yee MC, Kohli S, Glicksberg BS, Gignoux CR, Wojcik GL, Van Vleck T, Jeff JM, et al. 2017. Genetic identification of a common collagen disease in puerto ricans via identity-by-descent mapping in a health system. Elife 6.

Benn Torres J, Martucci V, Aldrich MC, Vilar MG, MacKinney T, Tariq M, Gaieski JB, Bharath Hernandez R, Browne ZE, Stevenson M, et al. 2019. Analysis of biogeographic ancestry reveals complex genetic histories for indigenous communities of St. Vincent and Trinidad. Am J Phys Anthropol 169:482–497.

Benn-Torres J. 2014. Prospecting the past: Genetic perspectives on the extinction and survival of indigenous poeples of the Caribbean. New Genetics and Society 33:21–41.

Benn-Torres J, Kittles RA, Stone AC. 2007. Mitochondrial and Y chromosome diversity in the English-speaking Caribbean. Ann Hum Genet 71:782–790.

Benn-Torres J, Vilar MG, Torres GA, Gaieski JB, Bharath Hernandez R, Browne ZE, Stevenson M, Walters W, Schurr TG, Genographic C. 2015. Genetic Diversity in the Lesser Antilles and Its Implications for the Settlement of the Caribbean Basin. PLoS One 10:e0139192.

Briggs AW, Stenzel U, Johnson Philip LF, Green RE, Kelso J, Prüfer K, Meyer M, Krause J, Ronan MT, Lachmann M, et al. 2007. Patterns of damage in genomic DNA sequences from a Neandertal. Proc Natl Acad Sci U S A 104:14616–14621.

Bryc K, Velez C, Karafet T, Moreno-Estrada A, Reynolds A, Auton A, Hammer M, Bustamante CD, Ostrer H. 2010. Colloquium paper: genome-wide patterns of population structure and admixture among Hispanic/Latino populations. Proc Natl Acad Sci U S A 107 Suppl 2:8954–8961.

Burney D, Burney LP, McPhee RDE. 1994. Holocene Charcoal Stratigraphy from Laguna Tortuguero, Puerto Rico, and the Timing of Human Arrival on the Island. J Archaeol Sci 21:273–281.

Bustamante CD, De La Vega FM, Burchard EG. 2011. Genomics for the world. Nature 475:163–165.

Carpenter ML, Buenrostro JD, Valdiosera C, Schroeder H, Allentoft ME, Sikora M, Rasmussen M, Gravel S, Guillen S, Nekhrizov G, et al. 2013. Pulling out the 1%: whole-genome capture for the targeted enrichment of ancient DNA sequencing libraries. Am J Hum Genet 93:852–864.

Chanlatte Baik LA. 2003. Agricultural societies in the Caribbean: The Greater Antilles, and the Bahamas. In. General History of the Caribbean, Vol. I. Autochtonous Societies. London: UNESCO Publishing. p. 228–258

Coppa A, Cucina A, Hoogland MLP, Lucci M, Luna Calderón F, Panhuysen RGAM, Tavarez Maria G, Valcárcel Rojas R, Vargiu R. 2008. New evidence of two different migratory waves in the Circum-Caribbean area during the pre-Columbian period from the analysis of dental morphological traits. In: Hofman CL, Hoogland MLP, van Gijn AL, editors. Crossing the borders: New methods and techniques in the study of archaeological materials from the Caribbean. Tuscaloosa: University of Alabama Press. p. 195–213

Curet LA. 2014. The Taino: Phenomena, Concepts, and Terms. Ethnohistory 61:467–495.

Curet LA, Torres J, Rodríguez M. 2004. Political and social history of Eastern Puerto Rico: the Ceramic age. In: Hofman CL, Delpuech A, editors. Late Ceramic Age Societies in the Eastern Caribbean. Oxford: Paris Monographs in American Archaeology 14. p. 59–86

Dabney J, Knapp M, Glocke I, Gansauge MT, Weihmann A, Nickel B, Valdiosera C, Garcia N, Pääbo S, Arsuaga JL, et al. 2013. Complete mitochondrial genome sequence of a Middle Pleistocene cave bear reconstructed from ultrashort DNA fragments. Proc Natl Acad Sci U S A 110:15758–15763.

Drummond AJ, Suchard MA, Xie D, Rambaut A. 2012. Bayesian Phylogenetics with BEAUti and the BEAST 1.7. Mol Biol Evol 29:1969–1973.

Duggan AT, Harris AJT, Marciniak S, Marshall I, Kuch M, Kitchen A, Renaud G, Southon J, Fuller B, Young J, et al. 2017. Genetic Discontinuity between the Maritime Archaic and Beothuk Populations in Newfoundland, Canada. Curr Biol 27:3149–3156 e3111.

Excoffier L, Novembre J, Schneider S. 2000. SIMCOAL: a general coalescent program for the simulation of molecular data in interconnected populations with arbitrary demography. J Hered 91:506–509.

Excoffier L, Smouse PE, Quattro JM. 1992. Analysis of molecular variance inferred from metric distances among DNA haplotypes: application to human mitochondrial DNA restriction data. Genetics 131:479.

Forte MC. 2006. Indigenous resurgence in the contemporary Caribbean: Amerindian survival and revival. New York: Peter Lang Publishing.

Fu Q, Mittnik A, Johnson PL, Bos K, Lari M, Bollongino R, Sun C, Giemsch L, Schmitz R, Burger J, et al. 2013. A revised timescale for human evolution based on ancient mitochondrial genomes. Curr Biol 23:553–559.

Fuller H, Benn Torres J. 2018. Investigating the “Taíno” ancestry of the Jamaican Maroons: a new genetic (DNA), historical, and multidisciplinary analysis and case study of the Accompong Town Maroons. Canadian Journal of Latin American and Caribbean Studies 43.

Gamba C, Jones ER, Teasdale MD, McLaughlin RL, Gonzalez-Fortes G, Mattiangeli V, Domboroczki L, Kovari I, Pap I, Anders A, et al. 2014. Genome flux and stasis in a five millennium transect of European prehistory. Nat Commun 5:5257.

Gomez-Carballa A, Catelli L, Pardo-Seco J, Martinon-Torres F, Roewer L, Vullo C, Salas A. 2015. The complete mitogenome of a 500-year-old Inca child mummy. Sci Rep 5:16462.

Gonzalez-Martin A, Gorostiza A, Regalado-Liu L, Arroyo-Pena S, Tirado S, Nuno-Arana I, Rubi-Castellanos R, Sandoval K, Coble MD, Rangel-Villalobos H. 2015. Demographic History of Indigenous Populations in Mesoamerica Based on mtDNA Sequence Data. PLoS One 10:e0131791.

Gorostiza A, Acunha-Alonzo V, Regalado-Liu L, Tirado S, Granados J, Sámano D, Rangel-Villalobos H, González-Martín A. 2012. Reconstructing the History of Mesoamerican Populations through the Study of the Mitochondrial DNA Control Region. PLoS One 7:e44666.

Gravel S, Zakharia F, Moreno-Estrada A, Byrnes JK, Muzzio M, Rodriguez-Flores JL, Kenny EE, Gignoux CR, Maples BK, Guiblet W, et al. 2013. Reconstructing Native American migrations from whole-genome and whole-exome data. PLoS Genet 9:e1004023.

Haslip-Viera G. 2001. Taino Revival: Critical prespectives on Puerto Rican identity and cultural politics. Princeton: Markus Weiner Publishers.

Hunley K, Gwin K, Liberman B. 2016. A Reassessment of the Impact of European Contact on the Structure of Native American Genetic Diversity. PLoS One 11:e0161018.

Jónsson H, Gonolhac A, Schubert M, Johnson P, Orlando L. 2013. mapDamage2.0: fast approximate Bayesian estimates of ancient DNA damage parameters. Bioinformatics 29:1682–1684.

Jørsboe E, Hanghøj K, Albrechtsen A. 2017. fastNGSadmix: admixture proportions and principal component analysis of a single NGS sample. Bioinformatics 33:3148–3150.

Keegan WF, Maclachlan MD. 1989. The Evolution of Avunculocal Chiefdoms: A reconstruction of Taino Kinship and politics. Am Anthropol 91:613–630.

Keegan WF, Rodriguez Ramos R. 2005. Archaic origins of the Classic Tainos. Proceedings of the 21st International Congress for Caribbean Archaeology; Trinidad. p. 211–217

Kircher M, Sawyer S, Meyer M. 2012. Double indexing overcomes inaccuracies in multiplex sequencing on the Illumina platform. Nucleic Acids Res 40:e3.

Korneliussen TS, Albrechtsen A, Nielsen R. 2014. ANGSD: Analysis of next generation sequencing data. BMC Bioinform 15.

Kumar S, Bellis C, Zlojutro M, Melton PE, Blangero J, Curran JE. 2011. Large scale mitochondrial sequencing in Mexican Americans suggests a reappraisal of Native American origins. BMC Evol Biol 11.

Laffoon JE, Hoogland MLP. 2012. Migration and mobility in the circum-Caribbean: Integrating archaeology and isotopic analysis. In: Kaiser E, Burger J, Schier W, editors. Population Dynamics in Prehistory and Early History: New Approaches Using Stable Isotopes and Genetics. Berlin: De Gruyter. p. 337–353

Lalueza-Fox C, Calderón FL, Calafell F, Morera B, Bertranpetit J. 2001. MtDNA from extinct Tainos and the peopling of the Caribbean. Ann Hum Genet 65:137–151.

Lalueza-Fox C, Gilbert MTP, Martinez-Fuentes AJ, Calafell F, Bertranpetit J. 2003. Mitochondrial DNA from pre-Columbian Ciboneys from Cuba and the prehistoric colonization of the Caribbean. Am J Phys Anthropol 121:97–108.

Lanfear R, Calcott B, Ho SY, Guindon S. 2012. Partitionfinder: combined selection of partitioning schemes and substitution models for phylogenetic analyses. Mol Biol Evol 29:1695–1701.

Lee EJ, Merriwether DA. 2015. Identification of Whole Mitochondrial Genomes from Venezuela and Implications on Regional Phylogenies in South America. Hum Biol 87:29–38.

Leigh J, Bryant D. 2015. PopART: Full-feature software for haplotype network construction. Methods Ecol Evol 6:1110–1116.

Lewis CM, Jr., Lizarraga B, Tito RY, Lopez PW, Iannacone GC, Medina A, Martinez R, Polo SI, De La Cruz AF, Caceres AM, et al. 2007. Mitochondrial DNA and the peopling of South America. Hum Biol 79:159–178.

Li H, Durbin R. 2009. Fast and accurate short read alignment with Burrows–Wheeler transform. Bioinformatics 25:1754–1760.

Li H, Handsaker B, Wysoker A, Fennell T, Ruan J, Homer N, Marth G, Abecasis G, Durbin R, Genome Project Data Processing S. 2009. The Sequence Alignment/Map format and SAMtools. Bioinformatics 25:2078–2079.

Lindo J, Huerta-Sanchez E, Nakagome S, Rasmussen M, Petzelt B, Mitchell J, Cybulski JS, Willerslev E, DeGiorgio M, Malhi RS. 2016. A time transect of exomes from a Native American population before and after European contact. Nat Commun 7:13175.

Llamas B, Fehren-Schmitz L, Valverde G, Soubrier J, Mallick S, Rohland N, Nordenfelt S, Valdiosera C, Richards SM, Rohrlach A, et al. 2016. Ancient mitochondrial DNA provides high-resolution time scale of the peopling of the Americas. Sci Adv 2:e1501385.

Madrilejo N, Lombard H, Torres JB. 2015. Origins of marronage: Mitochondrial lineages of Jamaica’s Accompong Town Maroons. Am J Hum Biol 27:432–437.

Maricic T, Whitten M, Paabo S. 2010. Multiplexed DNA sequence capture of mitochondrial genomes using PCR products. PLoS One 5:e14004.

Martínez-Cruzado JC. 2010. The History of Amerindian mitochondrial DNA lineages in Puerto Rico. In: Fitzpatrick S, Ross A, editors. Island shores, distant pasts: Archaeological and biological approaches to the Pre-Columbian settlement of the Caribbean. Gainesville: University Press of Florida. p. 54–80

Martínez-Cruzado JC. 2002. The Use of Mitochondrial DNA to Discover Pre-Columbian Migrations to the Caribbean: Results for Puerto Rico and Expectations for the Dominican Republic. J Caribb Amerind Hist Anthro:1–11.

Martínez-Cruzado JC, Toro-Labrador G, Ho-Fung V, Estevez-Montero M, Lobaina-Manzanet A, Padovani-Claudio DA, Sanchez-Cruz H, Ortiz-Bermudez P, Sanchez-Crespo A. 2001. Mitochondrial DNA Analysis Reveals Substantial Native American Ancestry in Puerto Rico. Hum Biol 73:491–511.

Martínez-Cruzado JC, Toro-Labrador G, Viera-Vera J, Rivera-Vega MY, Startek J, Latorre-Esteves M, Roman-Colon A, Rivera-Torres R, Navarro-Millan IY, Gomez-Sanchez E, et al. 2005. Reconstructing the population history of Puerto Rico by means of mtDNA phylogeographic analysis. Am J Phys Anthropol 128:131–155.

Mendisco F, Pemonge MH, Leblay E, Romon T, Richard G, Courtaud P, Deguilloux MF. 2015. Where are the Caribs? Ancient DNA from ceramic period human remains in the Lesser Antilles. Philos Trans R Soc Lond B Biol Sci 370:20130388.

Mendizabal I, Sandoval K, Berniell-Lee G, Calafell F, Salas A, Martinez-Fuentes A, Comas D. 2008. Genetic origin, admixture, and asymmetry in maternal and paternal human lineages in Cuba. BMC Evol Biol 8:213.

Meyer M, Kircher M. 2010. Illumina sequencing library preparation for highly multiplexed target capture and sequencing. Cold Spring Harb Protoc 6:1–10.

Mizuno F, Gojobori J, Wang L, Onis K. 2014. Complete mitogenome analysis of indigenous populations in Mexico: its relevance for the origin of Mesoamericans. J Hum Genet 59:359–367.

Mol AA. 2013. Studying Pre-Columbian Interaction Networks: Mobility and Exchange. In: Keegan WF, Hofman CL, Rodriguez Ramos R, editors. The Oxford Handbook of Caribbean Archaeology. Oxford: Oxford University Press. p. 329–346

Moreno-Estrada A, Gravel S, Zakharia F, McCauley JL, Byrnes JK, Gignoux CR, Ortiz-Tello PA, Martinez RJ, Hedges DJ, Morris RW, et al. 2013. Reconstructing the population genetic history of the Caribbean. PLoS Genet 9:e1003925.

Mulligan CJ, Kitchen A, Miyamoto MM. 2008. Updated three-stage model for the peopling of the Americas. PLoS One 3:e3199.

Nieves-Colón MA, Ozga AT, Pestle WJ, Cucina A, Tiesler V, Stanton TW, Stone AC. 2018. Comparison of two ancient DNA extraction protocols for skeletal remains from tropical environments. Am J Phys Anthropol 166:824–836.

Noguera-Santamaría MC, Edlund Anderson C, Uricoechea D, Durán C, Briceño-Balcázar I, Bernal Villegas J. 2015. Mitochondrial DNA analysis suggests a Chibchan migration into Colombia. Univ. Sci. 20:261–278.

O’Fallon BD, Fehren-Schmitz L. 2011. Native Americans experienced a strong population bottleneck coincident with European contact. Proc Natl Acad Sci U S A 108:20444–20448.

Ozga AT, Nieves-Colón MA, Honap TP, Sankaranarayanan K, Hofman CA, Milner GR, Lewis CM, Jr., Stone AC, Warinner C. 2016. Successful enrichment and recovery of whole mitochondrial genomes from ancient human dental calculus. Am J Phys Anthropol 160:220–228.

Patterson N, Moorjani P, Luo Y, Mallick S, Rohland N, Zhan Y, Genschoreck T, Webster T, Reich D. 2012. Ancient Admixture in Human History. Genetics 192:1065.

Patterson N, Price AL, Reich D. 2006. Population structure and eigenanalysis. PLoS Genet 2:e190.

Perego UA, Angerhofer N, Pala M, Olivieri A, Lancioni H, Hooshiar Kashani B, Carossa V, Ekins JE, Gomez-Carballa A, Huber G, et al. 2010. The initial peopling of the Americas: a growing number of founding mitochondrial genomes from Beringia. Genome Res 20:1174–1179.

Perego UA, Lancioni H, Tribaldos M, Angerhofer N, Ekins JE, Olivieri A, Woodward SR, Pascale JM, Cooke R, Motta J, et al. 2012. Decrypting the mitochondrial gene pool of modern Panamanians. PLoS One 7:e38337.

Pestle WJ. 2010. Diet and Society in Prehistoric Puerto Rico: An Isotopic Approach. [Ph.D. Thesis]: University of Illinois at Chicago.

Pestle WJ, Colvard M. 2012. Bone collagen preservation in the tropics: a case study from ancient Puerto Rico. J Archaeol Sci 39:2079–2090.

Pickrell JK, Reich D. 2014. Toward a new history and geography of human genes informed by ancient DNA. Trends Genet 30:377–389.

Raghavan M, Skoglund P, Graf KE, Metspalu M, Albrechtsen A, Moltke I, Rasmussen S, Stafford TW, Jr., Orlando L, Metspalu E, et al. 2014. Upper Palaeolithic Siberian genome reveals dual ancestry of Native Americans. Nature 505:87–91.

Renaud G, Slon V, Duggan AT, Kelso J. 2015. Schmutzi: estimation of contamination and endogenous mitochondrial consensus calling for ancient DNA. Genome Biol 16:224.

Reynolds AW, Raff JA, Bolnick DA, Cook DC, Kaestle FA. 2015. Ancient DNA from the Schild site in Illinois: Implications for the Mississippian transition in the Lower Illinois River Valley. Am J Phys Anthropol 156:434–448.

Rodríguez Ramos R. 2010. Caribbean Archaeology and Ethnohistory: Rethinking Puerto Rican Precolonial History. Tuscaloosa: University of Alabama Press.

Rodríguez Ramos R, Pagán Jiménez J, Hofman CL. 2013. The humanization of the insular Caribbean. In: Keegan WF, Hofman CL, Rodríguez Ramos R, editors. The Oxford Handbook of Caribbean Archaeology. Oxford: Oxford University Press. p. 126–140

Rogozinski J. 2008. A Brief History of the Caribbean: From the Arawak and Carib to the Present. New York: Penguin Books.

Rohland N, Hofreiter M. 2007. Ancient DNA extraction from bones and teeth. Nat Protoc 2:1756–1762.

Ross AH. 2004. Cranial evidence of pre-contact multiple population expansions in the Caribbean. Caribbean Journal of Science 40:291–298.

Rouse I. 1992. The Tainos: Rise & decline of the people who greeted Columbus: Yale University Press.

Sans M, Figueiro G, Hughes CE, Lindo J, Hidalgo PC, Malhi RS. 2015. A South American Prehistoric Mitogenome: Context, Continuity, and the Origin of Haplogroup C1d. PLoS One 10:e0141808.

Schroeder H, Ávila-Arcos MC, Malaspinas A-S, Poznik GD, Sandoval-Velasco M, Carpenter ML, Moreno-Mayar JV, Sikora M, Johnson PLF, Allentoft ME, et al. 2015. Genome-wide ancestry of 17th-century enslaved Africans from the Caribbean. Proc Natl Acad Sci U S A 112:3669–3673.

Schroeder H, Sikora M, Gopalakrishnan S, Cassidy LM, Maisano Delser P, Sandoval Velasco M, Schraiber JG, Rasmussen S, Homburger JR, Avila-Arcos MC, et al. 2018. Origins and genetic legacies of the Caribbean Taino. Proc Natl Acad Sci U S A 115:2341–2346.

Schubert M, Ginolhac A, Lindgreen S, Thompson JF, Al-Rasheid KA, Willerslev E, Krogh A, Orlando L. 2012. Improving ancient DNA read mapping against modern reference genomes. BMC Genomics 13.

Schurr TG, Benn Torres J, Vilar MG, Gaieski JB, Melendez C. 2016. An emerging history of indigenous Caribbean and circum-Caribbean populations: Insights from archaeological, ethnographic, genetic and historical studies. In: Zuckerman M, Martin DL, editors. New Directions in Biocultural Anthropology. New York: John Wiley & Sons.

Seguin-Orlando A, Hoover CA, Vasiliev SK, Ovodov ND, Shapiro B, Cooper A, Rubin EM, Willerslev E, Orlando L. 2015. Amplification of TruSeq ancient DNA libraries with AccuPrime Pfx: consequences on nucleotide misincorporation and methylation patterns. Sci Tech Arch Resear 1:1–9.

Skoglund P, Northoff BH, Shunkov MV, Derevianko AP, Paabo S, Krause J, Jakobsson M. 2014. Separating endogenous ancient DNA from modern day contamination in a Siberian Neandertal. Proc Natl Acad Sci U S A 111:2229–2234.

Skoglund P, Storå J, Götherström A, Jakobsson M. 2013. Accurate sex identification of ancient human remains using DNA shotgun sequencing. J Archaeol Sci 40:4477–4482.

Söchtig J, Álvarez-Iglesias V, Mosquera-Miguel A, Gelabert-Besada M, Gómez-Carballa A, Salas A. 2015. Genomic insights on the ethno-history of the Maya and the ‘Ladinos’ from Guatemala. BMC Genomics 16:131.

Torroni A, Schurr TG, Cabell MF, Brown MD, Neel JV, Larsen M, Smith DG, Vullo CM, Wallace DC. 1993. Asian affinities and continental radiation of the four founding Native American mtDNAs. Am J Hum Genet 53:563–590.

van Oven M, Kayser M. 2009. Updated comprehensive phylogenetic tree of global human mitochondrial DNA variation. Hum Mutat 30:E386–E394.

Veran C. 2003. Born Puerto Rican, born (again) Taino? A resurgence of indigenous identity among Puerto Ricans has sparked debates over the island’s tri-racial history. In. Colorlines Magazine. California. p. 23–25

Via M, Gignoux CR, Roth LA, Fejerman L, Galanter J, Choudhry S, Toro-Labrador G, Viera-Vera J, Oleksyk TK, Beckman K, et al. 2011. History shaped the geographic distribution of genomic admixture on the island of Puerto Rico. PLoS One 6:e16513.

Vilar MG, Melendez C, Sanders AB, Walia A, Gaieski JB, Owings AC, Schurr TG, Genographic C. 2014. Genetic diversity in Puerto Rico and its implications for the peopling of the Island and the West Indies. Am J Phys Anthropol 155:352–368.

Wang S, Lewis CM, Jakobsson M, Ramachandran S, Ray N, Bedoya G, Rojas W, Parra MV, Molina JA, Gallo C, et al. 2007. Genetic variation and population structure in native Americans. PLoS Genet 3:e185.

Weissensteiner H, Pacher D, Kloss-Brandstätter A, Forer L, Specht G, Bandelt H-J, Kronenberg F, Salas A, Schönherr S. 2016. HaploGrep 2: mitochondrial haplogroup classification in the era of high-throughput sequencing. Nucleic Acids Res 44:W58–W63.

Williams SR, Chagnon NA, Spielman RS. 2002. Nuclear and mitochondrial genetic variation in the Yanomamo: a test case for ancient DNA studies of prehistoric populations. Am J Phys Anthropol 117:246–259.

Wilson SM. 2007. The archaeology of the Caribbean. Cambridge: Cambridge University Press.

Wilson SM. 1999. Cultural pluralism and the emergence of complex society in the Greater Antilles. Proceedings of the 18th International Congress for Caribbean Archaeology; Grenada. p. 7–12

Wilson SM, Hester TR. 1998. Preceramic Connections between Yucatan and the Caribbean. Lat Am Antiq 9:342–352.

