## SupplementalMaterialsMethodsFigures for "Ancient DNA reconstructs the genetic legacies of pre-contact Puerto Rico communities"

### Supplementary Materials and Methods

**Ethics statement.** The human skeletal remains included in this study are considered the patrimony of the people of the Commonwealth of Puerto Rico. Permits for destructive sampling were obtained from the *Consejo para la Protección del Patrimonio Arqueológico Terrestre de Puerto Rico*, the *Autoridad de Carreteras y Transportación* (which has specific jurisdiction over the archaeological remains excavated from Paso del Indio), and the *Secretaría de Cultura y Turismo del Municipio Autónomo de Ponce* (which has jurisdiction over the Tibes site) (Figure S1). Permits were obtained by Dr. L. Antonio Curet (National Museum of the American Indian), Dr. William J. Pestle (University of Miami) and Dr. Edwin Crespo-Torres (University of Puerto Rico, Río Piedras Campus) as part of project NSF BCF-0612727 and extend into this project. Compliance with Native American Graves Protection and Repatriation Act regulations was not required for this research because no federally recognized tribes claim cultural affiliation to the remains, and the law has no jurisdiction in Puerto Rico (Hutt, et al. 1999; Ousley, et al. 2005; Siegel 2011). The skeletal remains were documented prior to destructive analysis (Figure S2).

**Bioarchaeological Context.** The site of Punta Candelero is located on a coastal peninsula in southeastern Humacao, Puerto Rico (Figure 1). It was identified and excavated between 1986-1989. The site had two successive occupation periods: 350 B.C. to A.D. 210, and A.D. 660 to 1010 (Figure S3). Each period is associated with distinct site usage patterns, ceramic and lithic assemblages, all of which suggest a multicomponent occupation. Household structures and general activity areas, including a central plaza built during the later period, have been identified at the site. No human burials were recovered from the early period, but 106 skeletal remains were found in late period strata, 78 of which have been identified as human (Fontanez 1991; Rodríguez López 1991; Crespo-Torres 2000).

The site of Paso del Indio is located in the alluvial plain of the Río Indio in north-central Vega Baja, Puerto Rico. It was identified and excavated between 1993-1995 due to construction of a major highway. The site was non-continuously occupied from 2690 B.C. to A.D. 1440. Post molds identified at the site indicate that many household structures, and potentially one central plaza were built at Paso del Indio throughout its occupation. One-hundred and thirty eight human skeletal remains were recovered from the site during excavations (Walker 2005). At the time of sample collection, skeletal remains from Paso del Indio and Punta Candelero were housed by Dr. Edwin Crespo-Torres at the Forensic Anthropology and Bioarchaeology Laboratory of the University of Puerto Rico. For this research, 44 individuals were sampled from Paso del Indio and 34 from Punta Candelero.

The site of Tibes is located in an alluvial terrace near the Río Portugués in southern Ponce, Puerto Rico. The site was first identified and excavated between 1975-1981. In 1982 an archaeological park and museum, the *Centro Ceremonial Indígena de Tibes* was built at the site. Research and excavations resumed in 1995 and continue to the present day. Tibes was continuously occupied between A.D. 300 – 1200. In addition to habitation areas it has several monumental structures including five stone lined plazas. To date, 126 human skeletal remains have been recovered from the site (Curet and Stringer 2010). Skeletal remains from Tibes are currently housed at the *Centro Ceremonial Indígena de Tibes* museum in Ponce, Puerto Rico. For this study, 46 individuals were sampled from Tibes.

In total, 124 human skeletal remains were sampled for destructive analysis across all three sites (Table S1). Direct radio carbon dates previously obtained for 81 of these remains indicate that most individuals ranged between A.D. 500-1300 (Pestle and Colvard 2012). Morphological sex assessment for individuals with sufficient macroscopic preservation was previously determined by a bioarcheologist (Crespo-Torres 2000; Pestle 2010).

**Sampling and DNA extraction.** Tooth, long bone fragments and dental calculus were collected from the human skeletal remains included in this study. Sample processing and DNA extractions were conducted at the Arizona State University (ASU) Ancient DNA Laboratory, a Class 10,000 clean-room facility. Before destructive sampling, remains were photographed and documented (Figure S3). To eliminate surface contaminants and inhibitors, tooth and long bone fragments were cleaned with a 1% sodium hypochlorite solution and the outer surface was mechanically removed with a circular Dremel wheel (Rohland and Hofreiter 2007). Samples were UV irradiated for 5 minutes on each side in a UVP CL-1000 Ultraviolet Crosslinker. Teeth were sliced transversally at the cemento-enamel junction using the Dremel. The roots were covered in aluminum foil and pulverized by blunt force with a hammer as in (Schuenemann, et al. 2011). Long bone fragments were pulverized in a Spex CertiPrep 8000M Mixer/Mill. When present, dental calculus was sampled from the tooth surface with a dental scaler following (Warinner, et al. 2014). Throughout the sampling process strict controls were implemented to minimize potential sample contamination. These included single use of Dremel wheels, UV irradiation of tools and work area before and between uses, full body coverings and bleach decontamination, among others (Cooper and Poinar 2000; Gilbert, et al. 2006; Llamas, et al. 2017). Samples were extracted using several silica-based extraction methods optimized for ancient DNA (aDNA) (Rohland and Hofreiter 2007; Dabney, et al. 2013; Nieves-Colón, et al. 2018). Between 30-100 mg of dentine or bone powder were used for tooth and long bone extractions. Calculus extractions were performed with all the calculus obtained, usually between 2-20 mg. DNA yields in ng/uL were measured using 1 $\mu$l of each extract through fluorometric quantification with the Qubit 2.0 High Sensitivity assay (Simbolo, et al. 2013). Extraction blanks were included throughout this process. Some samples were extracted more than once to obtain more DNA (Table S2).

**Library preparation.** Double stranded libraries were produced with 20 μl of each extract following the protocol by (Meyer and Kircher 2010) using the Qiagen MinElute PCR purification kit instead of SPRI beads. Extraction blanks were also converted into libraries. A negative library control was included per library batch. 1:100 dilutions of each library were quantified using qPCR. Reactions were run in triplicate for each library in final volumes of 20 μl with the following conditions: 10 μl of 2X Dynamo SYBR Green qPCR Master Mix with 0.3x ROX (Thermo Scientific), 1 μl of primer IS7 (5'-ACACTCTTTCCCT ACACGAC-3') at 10 μM, 1 μl of primer IS8 (5'-GTGACTGGAGTTCAGACGTGT-3') at 10 μM, 7 μl of $ddH_20$, and 1 μl of the library dilution. Reactions were heated to 95°C for 10 minutes, then 40 cycles of 95°C for 15 seconds and 60°C for 1 minute for 40 cycles. A final disassociation stage was added at the end of these cycles: 95°C for 15 seconds, 60°C for 15 seconds and 95°C for 15 seconds. Quantification was performed using an ABI7900HT thermocycler, and results were analyzed with SDS software. Analysis of qPCR data focused on cycle threshold values (Ct), which represent the number of qPCR cycles required for a fluorescent signal to exceed background levels. Mean Ct values were averaged across all replicates per library. Non-template controls (NTC), which have no DNA, were also included in the reaction to monitor background fluorescent levels.

All libraries were double indexed and amplified following (Kircher, et al. 2012). The Illumina specific indexing primers used were P5 (5'-AATGATACGGCGACCACCGAGATCTACACxxxxxx ACACTCTTTCCCTACACGACGCTCTT-3') and P7 (5'-CAAGCAGAAGACGGCATACGAGAT xxxxxx GTGACTGGAGTTCAGACGTGT-3'). Unique index combinations for all samples (represented by x above) are provided in Table S2. To increase library complexity, four 100 μl indexing reactions were performed per library. Samples processed between 2012-2014 were amplified with *Pfu*Turbo enzyme (Agilent) for 10 cycles with the following reaction conditions: 10 μl of 10X *Pfu*Turbo Buffer, 2.50 μl of 10 mM dNTPs, 1.50 μl of 10 mg/ml Bovine Serum Albumin, 2 μl of P5 indexing primer at 10000 nM, 2 μl of P7 indexing primer at 10000 nM, 1 μl of *Pfu*Turbo enzyme. Samples processed after 2014 were indexed for 15-20 cycles with AmpliTaq Gold® enzyme (Life Technologies) following recommendations by (Seguin-Orlando, et al. 2015). Reaction conditions were as follows: 9.27 μl of 10X PCR Buffer II, 3.68 μl of 10 mM dNTPs, 2.21 μl of 10 mg/ml Bovine Serum Albumin, 9.27 μl of 25 mM Gold $MgCl_2$ solution, 2 μl of P5 indexing primer at 10000 nM, 2 μl of P7 indexing primer at 10000 nM, 61.09 μl of $ddH_20$, 1.48 μl of AmpliTaq Gold® enzyme and 9 μl of DNA library. All reactions irrespective of enzyme were heated to 95°C for 15 minutes for initial denaturation. Further denaturation, annealing and elongation were performed at 95°C for 30 seconds, 58°C for 30 seconds and 72°C for 45 seconds for the determined amount of cycles. Final extension was performed at 72°C for 10 minutes, and reactions were kept at 10°C. All four aliquots of each indexed library were combined, and the library was purified with the Qiagen MinElute PCR purification kit following manufacturer instructions with the following modification: EB buffer was preheated to 65°C, and elution was performed in 30 $\mu$l. 1 $\mu$l of each library was used for quantification with the Qubit 2.0 Broad Range assay. Purified libraries were further diluted to a factor of 1:1000 and quantified using the KAPA Library Quantification kit (Kapa Biosystems) following manufacturer instructions on the ABI7900HT thermocycler. Fragment analysis of the indexed libraries was performed using the DNA 1000 assay on the Agilent 2100 Bioanalyzer.

Libraries initially indexed for 10 cycles which had low post purification DNA concentrations were re-amplified to obtain sufficient DNA for mitochondrial genome (mtDNA) capture (300-500 ng). Libraries were divided into four 100 $\mu$L aliquots as explained above. Re-amplification conditions were: 10 $\mu$l of 10X Accuprime™ *Pfx* reaction mix, 3 $\mu$l of IS5 primer at 10 $\mu$M, 3 $\mu$l of IS6 primer at 10 $\mu$M, 76 $\mu$l of $ddH_20$, 1 $\mu$l of Accuprime™ *Pfx* enzyme and 7 $\mu$l of DNA library. Reactions were heated to 95°C for 2 minutes for initial denaturation; further denaturation, annealing and elongation were performed at 95°C for 15 seconds, 60°C for 30 seconds and 68°C for 1 minute for 7-13 cycles. Final extension was performed at 68°C for 5 minutes, and reactions were then kept at 4°C. Subsequent purification and quantification was performed as above.

**Mitochondrial target enrichment capture and Illumina sequencing.** In-solution and targeted enrichment for the complete mitochondrial genome was performed as in (Maricic, et al. 2010) with modifications as in (Ozga, et al. 2016). DNA libraries were pooled in equimolar amounts up to 2 $\mu$g per capture pool. After enrichment, libraries were amplified, purified and quantified using the same conditions as above. Enriched libraries (including blanks) were sequenced on multiple runs of the Illumina MiSeq (2 x150 bp) at the DNASU Sequencing Core at ASU. Some enriched libraries were captured and sequenced more than once to increase read depth and genomic coverage (Table S2-S3).

**Shotgun sequencing and whole genome enrichment.** Thirty-five ancient samples were shotgun sequenced on several runs of the Illumina HiSeq 2500 (2 x 100 bp) at the Yale Center for Genomic Analysis Core and the NextSeq 500 (2 x 75 bp) at Stanford University (see below). Based on analyses of the shotgun data (Table S4), 22 samples were selected for whole-genome enrichment. Two whole-genome enrichment methods were used in this study. The first enrichment was performed in 2014 in Dr. Carlos D. Bustamante's genetics laboratory at Stanford University, as part of a pilot experiment. Ten libraries were screened by shotgun sequencing on the Illumina NextSeq 500 (2 x 75 bp). The seven samples with the highest endogenous content were chosen for whole genome in-solution capture (WISC) which was performed as in (Carpenter, et al. 2013) with the following modifications. RNA bait libraries were created from a pool of human genomic DNA from three male individuals from the Coriell Hapmap populations (MKK, JPT and CEU). These individuals were chosen to maximize sequence diversity and ensure inclusion of Y-chromosome baits. Axygen® beads (1.8x volume) were used for bait library purification and xGEN® LNA Adaptor blockers for Illumina TruSeq were used to prevent non-specific binding to the baits. WISC enriched libraries were sequenced on all four lanes of the NextSeq (2 x 75 bp). All subsequent whole genome capture experiments were performed between 2016-2017 in the Molecular Anthropology Laboratory at ASU. Fifteen samples were enriched using the MYbaits Human Whole Genome Capture Kit (Arbor Biosciences), following manufacturer instructions. Enriched libraries were subsequently amplified with the 2X Kapa HiFi Hot Start Ready Mix (Kapa Biosystems) following manufacturer instructions. Purification and quantification were performed as detailed above. For these samples, sequencing was performed on several runs of the NextSeq 500 (2x 75 bp) and Illumina HiSeq 2500 (2 x 100 bp) at the University of Arizona Genetics Core, and one run of the Illumina HiSeq 2500 (2 x 100 bp) at the Yale Center for Genomic Analysis Core. Some enriched libraries were captured and sequenced more than once to increase read depth and genomic coverage (Table S4).

**MtDNA sequence read mapping and processing.** Illumina sequence reads were merged, and adapters trimmed using SeqPrep (https://github.com/jstjohn/SeqPrep) with a minimum overlap of 11 base pairs (bp) and a minimum length threshold of 30 bp. Read quality was assessed pre and post-merging with FastQC v.0.11.3 (http://www.bioinformatics.babraham.ac.uk/projects/fastqc/). For mitochondrial enriched libraries, reads were mapped to the revised Cambridge Reference Sequence (rCRS NCBI Reference Sequence: NC_012920.1). Sequence read mapping was performed using BWA v. 0.7.5 (Li and Durbin 2009) with seeding disabled (-1 1000) and edit distance increased (-n 0.01) to improve mapping accuracy (Schubert, et al. 2014). Reads were filtered with SAMtools v. 0.1.19 (Li, et al. 2009). for mapping quality ≥ Q30, duplicates were removed with the *rmdup* option and reads mapping to more than one location were discarded by controlling for XA, XT and X0 tags (Table S3). Damage patterns were characterized (Figure S5-S4) and quality scores were rescaled with mapDamage v.2.0.2 (Ginolhac, et al. 2011; Jónsson, et al. 2013). Summary statistics such as mean read depth, standard deviation of read depth and percent of reference sequence covered were estimated from the rescaled BAM files using Qualimap v.2.2.1 (Okonechnikov, et al. 2016). BAM files were visualized with Tablet v.1.16.09.06 (Milne, et al. 2013) and Geneious v.7.0.6 (Biomatters). Scripts used for processing mtDNA sequence reads and estimating contamination are publicly available on GitHub: https://github.com/mnievesc/Ancient_mtDNA_Pipeline Reads from enriched libraries sequenced over multiple runs were combined after duplicate removal using SAMtools (Table S3). Before merging, read group information was added to each sample replicate using the *AddOrReplace ReadGroups* module from Picard v.2.01 (http://broadinstitute.github.io/picard). After merging duplicate removal was repeated and damage patterns as well as summary statistics were recalculated. For all mtDNA enriched libraries, mitochondrial SNP variants were called using SAMtools *mpileup* on the rescaled BAM files with ploidy was set to 1 with the sample option. Variant calls were output in VCF format using bcftools. MtDNA haplogroup assignment was performed in HaploGrep 2.0 (Weissensteiner, et al. 2016). Confirmation of haplogroup defining mutations and Haplogrep assignments was performed manually with reference to Phylotree mtDNA tree Build 17 (van Oven and Kayser 2009).

**MtDNA contamination controls.** Contamination was estimated using the mitochondrial enriched reads with contamMix (Fu, et al. 2013) and schmutzi (Renaud, et al. 2015). ContamMix provides a Bayesian estimate of the proportion of contaminant DNA present in the sample reads and the proportion of authentic aDNA. A set of 311 complete human mtDNA genomes were used as potential contaminant sources. Insufficient read depth or coverage of the reference sequence can result in inaccurate contamination estimates. Therefore, only samples with ≥3x read depth and ≥90% mtDNA genome coverage were included in this assessment. We further estimated mitochondrial contamination using schmutzi by comparing our test samples to a reference database of 197 potential contaminant mitochondrial allele frequencies (provided with the program). Only samples with ≥5x read depth and ≥98% genome coverage were included in this analysis (Table S3).

Nine sequenced mtDNA enriched libraries (8 samples and one extraction blank) were identified as having potential contamination based on low proportions of estimated authentic reads, high average contamination estimates, unusually large average fragment lengths, unusual damage patterns and/or non-Native American haplogroups (Table S3). BAM files for these samples were filtered using PMDtools (Skoglund, et al. 2014) to retain only reads showing signs of aDNA damage (threshold: PMD > 3). Summary statistics were recalculated, and damage pattern assessment and haplogroup assignment were repeated for these samples after filtering (Table S6; Figure S7). Post-filtering, two samples (PC-443 and PC-448) retained sufficient endogenous reads to include in population genetics analyses.

**MtDNA consensus files.** Forty-five enriched mtDNA samples with ≥5x read depth and ≥98% coverage of the mitochondrial genome were selected for population genetics analyses. Reads for five of these samples were combined across multiple sequencing runs. For two samples, endogenous reads were recovered after contamination filtering with PMDtools. Quality filtered (>Q20) mtDNA consensus sequences for the 45 selected samples were produced with schmutzi. Consensus sequences were revised manually in Geneious and exported in FASTA format. Variant sites with no coverage were designated as N. Consensus sequences are provided in Supporting File S1. Unique haplotypes were determined by direct counting (Table S7).

**MtDNA comparative data and DNA alignments.** A reference dataset of 1,636 complete mtDNA genome sequences was assembled from the literature including data from ancient and modern Native Americans, as well as admixed individuals of Native American descent (Ingman, et al. 2000; Tamm, et al.

2007; Achilli, et al. 2008; Fagundes, et al. 2008; Perego, et al. 2009; Perego, et al. 2010; Barbieri, et al. 2011; Kashani, et al. 2011; Kumar, et al. 2011; Bodner, et al. 2012; Cardoso, et al. 2012; de Saint Pierre, et al. 2012; Achilli, et al. 2013; Cui, et al. 2013; Lippold, et al. 2014; Tascón Peñaranda 2014; Vilar, et al. 2014; 1000 Genomes Project, et al. 2015; Lee and Merriwether 2015; Söchtig, et al. 2015; Llamas, et al. 2016; Ozga, et al. 2016; Mizuno, et al. 2017; Arias, et al. 2018; Brandini, et al. 2018). A second dataset of 391 control region sequences was assembled including mtDNA data collected from present-day Caribbean individuals with Native American mtDNA ancestry from Cuba, Dominican Republic, Puerto Rico, the First People's Community (FPC) of Trinidad and the Garifuna of St. Vincent (Mendizabal, et al. 2008; Nieves-Colón 2012; Tascón Peñaranda 2014; Vilar, et al. 2014; Benn-Torres, et al. 2015). This dataset also includes aDNA sequences from individuals from pre-contact sites in Cuba, Dominican Republic and Guadeloupe (Lalueza-Fox, et al. 2001; Lalueza-Fox, et al. 2003; Mendisco, et al. 2015) (Table S10). Multiple alignments were prepared to account for variables sequence lengths of control region datasets. For comparative analyses including HVR-1 data from Caribbean populations, sequences were trimmed to a common region between positions 16056-16391, except when noted. Phylogenetically uninformative sites such as indels or poly-C stretches at positions 309, 315, 515-522, 3107, 16182-16183 or 16193 and mutational hotspot 16519 were excluded from analysis as recommended by (van Oven and Kayser 2009). Multiple sequence alignments were performed with the command line version of MAFFT v7.244 (Katoh and Standley 2013). Alignment trimming was done in Geneious v.7.0.6 (Biomatters). Format conversions were done with PGD Spider (Lischer and Excoffier 2012).

**MtDNA sequence and statistical analyses.** To detect potential diachronic changes in genetic diversity in pre-contact Puerto Rico (PC-PR), we examined the relationship between temporal and genetic distance for all radiocarbon dated remains (n=37) using a Mantel test. The test compared a Euclidean distance matrix of median calibrated A.D. radiocarbon dates and a Tamura-Nei genetic distance matrix calculated with gamma correction of 0.26 (Meyer, et al. 1999). The test was performed using the *mantel.test* function in the *ape* R package (Paradis, et al. 2004). Significance *($p<0.05$)* was evaluated after 10,000 permutations (Figure S9).

To characterize patterns of genetic variation, intra-population diversity measures such as number of haplotypes (*h*), number of segregating sites (*S*), nucleotide ($\pi$) and haplotype diversity (*Hd*) (Nei 1987; Nei and Miller 1990) were calculated using the *haplotype*, *seg.sites*, *hap.div* and *nuc.div* functions from the *pegas* R package (Paradis 2010). Diversity measures obtained for the full PC-PR sample were compared to: (1) a subset of 1,403 ancient and modern complete mtDNA sequences classified into 47 population groups (Table S10), and (2) HVR-1 sequences for three ancient and five modern Caribbean populations (Table S11). All measures were calculated using default parameters and considering base ambiguities. Default parameter exclude indels as segregating sites. Thus, haplotypes defined by indels are collapsed in this analysis.

We measured inter-population genetic differentiation by using exact tests to compare haplotype frequencies between the three PC-PR sites (Table S8), between PC-PR and continental reference populations (complete mtDNA; Table S13) and between PC-PR and Caribbean populations (HVR-1 data only; Table S15). The exact tests were performed using the *test_diff* function from the *genepop* R package (Rousset 2008) using a Markov chain with 10,000 steps, 100 batches and 5,000 iterations. Switch values >1000 indicate appropriate convergence of the MC chain. Significant results (*p<0.05)* reject the null hypothesis that haplotypes are randomly distributed across all populations (panmixia). Additionally, mtDNA sequences were used to calculate inter-population pairwise $\Phi_{st}$ genetic distance measures using the *pairPhiST* function in the haplotypes R package (Excoffier, et al. 1992; Aktas 2019). Significant comparisons after 1,000 permutations (*p<0.05)* reject the null hypothesis of no difference between populations (Tables S9, S14; Figures S). Heat maps visually representing $\Phi_{st}$ distances were plotted with the *lattice* R package (Sarkar 2008) (Figures S8, S11-12). Pairwise $\Phi_{st}$ distance matrices were used as input for non-metric multidimensional scaling (MDS) calculated with the *metaMDS* function in the *vegan* R package (Oksanen, et al. 2018) (Figures 3, 5, S13). Lastly, haplogroup (e.g. A,B,C,D,X) frequencies per population were estimated by direct counting and correspondence analysis was performed and visualized using the *CA* function from the *FactoMineR* R package, and other functions from the *corrplot, factoextra, gridExtra, gplots* and *ggplot2* packages (Figure 4, S14) (Wickham 2009; Warnes, et al. 2016; Auguie 2017; Kassambara and Mundt 2017; Wei and Simko 2017).

To examine mtDNA lineage sharing across the pre and post-contact Caribbean, and between PC-PR and continental Native American populations, haplotype networks were constructed in popART (Bandelt, et al. 1999; Leigh and Bryant 2015) with default parameters. Indels and sites with >5% missing data were excluded from network reconstruction. FASTA files were converted to NEXUS format using PGD Spider (Lischer and Excoffier 2012). Complete mtDNA and HVR1 networks were constructed comparing PC-PR lineages belonging to haplogroups A2, C1 and D1 to comparative populations from across the Americas and the Caribbean. A second round of analysis included only complete mtDNA lineages from haplogroups A2 and C1 found in PC-PR and modern Puerto Ricans. No D1 lineages were present in the complete mtDNA data available from present-day Puerto Ricans (Figures 6, S15-18).

**Demographic modeling and test of continuity (mtDNA).** The pre-contact demographic history of Puerto Rico and the Caribbean was analyzed using an approximate Bayesian computation (ABC) approach. For this analysis, aDNA HVR-1 sequences were trimmed to a common region between 16056-16400 present in all populations. We used the Bayesian Serial SimCoal (BayeSSC) program (Excoffier, et al. 2000; Anderson, et al. 2005) to simulate the available HVR-1 data under eight possible demographic scenarios (M1-M8) with several variants (a,b,c,d), shown in (Figure S19). Scenario M1 models the ancient HVR-1 data from Guadalupe, Cuba, Puerto Rico, and the Dominican Republic, as a single panmictic population. In scenario M2, each island population split from the others at some point in the past (between 1,000-2,500 years ago) followed by a period of no migration between islands. Scenario M3 builds on M2 by adding varying levels of migration (0.01, 0.05, 0.1, 0.2) between all of the islands each generation. Scenario M4 limits this migration so that it only occurs between the islands closest to each other geographically. Scenarios M5 and M6 model single migration events of varying sizes (0.01, 0.05, 0.1, 0.25) from West to East and East to West respectively, after the initial peopling of the islands. Scenarios M7 and M8 model the peopling of the islands as a stepping stone event from West to East and East to West, respectively.

Following Duggan, et al. (2017), each model was parameterized using prior estimates of population size and timing of the population expansion into the Americas. All simulations were performed using an ancestral population size of 1,000 that expanded exponentially between 15,000 and 14,000 years ago (ya) followed by a period of stable population size, while allowing for local population dynamics in more recent time periods (between 1,000-2.500 ya). The following parameters of sequence evolution were used for all simulations: generation time of 25 years, a fixed mutation rate of $1.6427333 \times 10^{7}$ (Soares, et al. 2009), transition:transversion ratio of 0.9841 (Kimura 1980), and a gamma distribution of rates with shape parameters of 0.205 (theta) and 10 (kappa) (Ho and Endicott 2008). Modern population sizes were drawn from a uniform distribution between 2,000 and 1,0000 individuals. For models where island populations diverge from one another in the past (all but M1), growth rates following this divergence were determined by the population size at the time of the divergence, the timing of the divergence, and the modern population size. 500,000 genealogies were simulated for each model.

To determine which of our models most likely explain the observed HVR-1 data, we compared our simulated datasets to the empirical data using Euclidean distances (Beaumont, et al. 2002). Euclidean distances were calculated from summary statistics calculated for each simulated dataset and those calculated from the empirical data. Private haplotypes, haplotype diversity, and *Fst* were used in this analysis, following the recommendations of (Duggan, et al. 2017). The fraction of simulations with the smallest Euclidian distance to the empirical summary statistics was retained (1%) to construct posterior distributions of population parameters using the functions from the R package included with BayeSSC (available at http://web.stanford.edu/group/ hadlylab/ssc/) to select the "maximum credible" version of each model. The prior distribution used in each model was replaced with maximum-likelihood estimation values from the posterior distributions, and each model was run for 1,000 genealogies. The goodness of fit for the different models was compared using the Akaike information criterion (AIC) (Akaike 1974) and Akaike weights x (Posada and Buckley 2004). The model with the lowest goodness-of-fit (AIC) value and the highest relative likelihood value is the best fit for the available data (Table S17).

We also used BayeSSC simulations to evaluate whether the observed genetic distance (*Fst*) between the complete mtDNA genomes collected from ancient and present-day Puerto Rico samples were consistent with a model of population continuity on the island, where the only evolutionary forces acting on the population were drift and mutation. The model used a per base substitution rate of $1.23 \times 10^{-8}$ substitutions/site/year for the entire mtDNA genomes (Soares, et al. 2009), a mitochondrial sequence length of 16,565 bp, and a gamma distribution of rates with shape parameters of 0.205 (theta) and 10 (kappa) (Ho and Endicott 2008). This model was simulated 10,000 times to produce a distribution of possible *Fst* values for comparison with our observed *Fst*. Results suggest that we cannot reject the null hypothesis of continuity between the ancient and modern Puerto Rico samples ($p$=0.0804).

**Estimation of effective population size and demographic history (mtDNA).** Effective population size (*Ne*) was estimated with BEAST 1.8.4 (Drummond, et al. 2012). This analysis included 127 aligned complete mtDNA sequences: 45 from PC-PR (this study), 81 from present-day Puerto Ricans of Native American ancestry included in the 1000 Genomes Phase 3 (1000 Genomes Project, et al. 2015), and 12 present-day Puerto Ricans sequenced by Vilar, et al. (2014). Common indels and mutation hotspots at nucleotide positions 309.1C(C), 315.1C, AC indels at 515–522, 16182, 16183, 16193.1C(C), and 16519 were excluded from the alignment as recommended by (van Oven and Kayser 2009). The complete mtDNA sequences were partitioned into five concatenated regions: D-loop, rRNA, tRNA, genes on the heavy strand, and ND6 on the light strand (Dloop+Coding+ND6+rRNA+tRNA). PartitionFinder v.1.1.1 (Lanfear, et al. 2012) was used to determine the best substitution model for three different datasets: (1) All sequences: TrN+G, (2) Haplogroup A: HKY+G, and (3) Haplogroup C: HKY+G.

BEAST was used with tip date calibrations (median cal A.D. radiocarbon dates) to reconstruct the phylogeny. No outgroup was included. Instead we constrained monophyly for haplogroups A, B, C, D, A+B and C+D, to obtain a more reliable skyline plot without violating the assumption of panmixia. Given that accounting for uncertainty in tip dates has little impact on posterior estimates (Molak, et al. 2013; Molak, et al. 2015), we used point estimates for tip calibrations. Individuals that did not have an associated radiocarbon date were assigned a prior date range based on the archaeological context, and posterior dates were estimated based on the molecular rate calculated empirically (Table S18-S19).

A strict clock could not be rejected according to preliminary analysis with uncorrelated lognormal relaxed clock, and so was used for all subsequent analyses. Three Markov Chain Monte Carlo (MCMC) chains of 100 million generations were performed for each dataset (All sequences (n=127), Haplogroup A (n=63), Haplogroup C (n=47), with sampling of parameters every 10,000 generations. Parameter traces were monitored in Tracer v1.6 to ensure convergence of the MCMC chains and effective sample sizes higher than 200. The initial 10 million (40 million for Haplogroup C) steps were discarded as burn-in before sampled trees and parameter traces from the three independent chains were combined to summarize the results.

Demographic history was reconstructed using the extended Bayesian skyline method (Heled and Drummond 2008), which infers *Ne* through time and also estimates the number of demographic changes from the data. The skyline plot was reconstructed using a Java program (O'Fallon and Fehren-Schmitz 2011) and visualised with *ggplot2*, assuming a generation time of 25 years (Figures S20-S24). A date randomisation test (Ho and Shapiro 2011) was not performed because the temporal distribution of dated sequences is too narrow, thus the test may not perform well (Duchêne, et al. 2015)

**Whole genome sequence read mapping and processing.** Reads sequenced from shotgun and whole genome enriched libraries were processed in the same way as mitochondrial enriched libraries but mapping was done to the NCBI Build GRCh37 assembly (hg19) with the mtDNA sequence replaced by the rCRS (Andrews, et al. 1999). Sequence data for the two samples with highest read depth and endogenous content (PI-420a and PI-51) were produced across multiple sequencing runs and combined for analysis. Damage patterns for these two samples are shown in Figure S6. Mitochondrial reads for these two samples were subset from the genome-wide BAM files and re-mapped to the rCRS. Variant calling, haplogroup estimation and schmutzi contamination estimates were repeated for these two samples as described above. Chromosomal sex for these two individuals was determined based on the number of reads mapping to the X and Y chromosomes following (Skoglund, et al. 2013). For individual PI-51, sexed as a chromosomal male, we estimated nuclear contamination on the haploid X-chromosome with ANGSD v. 0.928 (Korneliussen, et al. 2014) (Table S5).

**Autosomal genotypes statistical analyses.** For population genetics analyses, the newly sequenced genomes obtained for the individuals PI-420a and PI-51 from Paso del Indio were analyzed alongside one "Lucayan Taino" individual from the pre-contact Bahamas (PC537) previously sequenced in (Schroeder, et al. 2018). A reference panel composed of 967 individuals from 37 ancient and present-day worldwide populations, with 597,573 SNPs was compiled from the literature. The panel included 656 individuals from the Americas (Meyer, et al. 2012; Lazaridis, et al. 2014; 1000 Genomes Project, et al. 2015; Raghavan, et al. 2015; Mallick, et al. 2016; Gnecchi-Ruscone, et al. 2019) (Table S20).

We generated haploid genotype calls in the ancient individuals by randomly sampling one read per overlapping positions. A genotype likelihood approach was implemented for admixture analysis using FastNGSadmix (Jørsboe, et al. 2017). Genotype likelihoods of the ancient samples were estimated using ANGSD (Korneliussen, et al. 2014) for all overlapping positions with the reference panel. Regular runs of admixture were performed with ADMIXTURE v. 1.3.0 (Alexander, et al. 2009) using only the reference panel individuals and after pruning for linkage disequilibrium (LD) with PLINK v. 1.9.0 (Purcell, et al. 2007). LD pruning was performed using a window of 200 SNPs, excluding pairs if $r^2 > 0.4$ and advancing by 25 SNPs at the time A total of ten runs were estimated for K3 to K7. We kept the run with the highest likelihood per K, which was then used to calculate the proportion of the ancestral components in the ancient genomes (Figure S25-S26).

Principal component analyses was performed with smartpca from Eigensoft 6.0.1 (Patterson, et al. 2006) using the *lsqproject* option to project the ancient individuals onto components inferred using present-day populations (Figure S27). Similarly, an outgroup-$f_3$ analysis of the form $f_3$(ancient, X; Yoruba) (Raghavan, et al. 2014) was performed, where 'ancient' represented one of the two ancient genomes from PC-PR and X included a set of 22 Americas populations and the PC537 individual (Figure S28). This analysis was performed using Admixtools *qp3pop* (Patterson, et al. 2012). All analyses were performed with and without transitions to account for damage in the ancient samples (Table S21).

**Computational resources.** This research was completed using the ASU Saguaro High Performance Computing (HPC) environment, the Stanford University SCG Genomics Clusters and the computational resources of the Center for Research Informatics' Gardner HPC cluster at the University of Chicago.

### References

1000 Genomes Project C, Auton A, Brooks LD, Durbin RM, Garrison EP, Kang HM, Korbel JO, Marchini JL, McCarthy S, McVean GA, et al. 2015. A global reference for human genetic variation. *Nature* 526:68-74.

Achilli A, Perego UA, Bravi CM, Coble MD, Kong QP, Woodward SR, Salas A, Torroni A, Bandelt HJ. 2008. The phylogeny of the four pan-American MtDNA haplogroups: implications for evolutionary and disease studies. *PLoS One* 3:e1764.

Achilli A, Perego UA, Lancioni H, Olivieri A, Gandini F, Hooshiar Kashani B, Battaglia V, Grugni V, Angerhofer N, Rogers MP, et al. 2013. Reconciling migration models to the Americas with the variation of North American native mitogenomes. *Proc Natl Acad Sci U S A* 110:14308-14313.

Akaike H. 1974. A new look at the statistical model identification. *IEEE T Automat Contr* 19:716-723.

Aktas C. 2019. Haplotypes: Manipulating DNA Sequences and Estimating Unambiguous Haplotype Network with Statistical Parsimony. R package version 1.1.

Alexander DH, Novembre J, Lange K. 2009. Fast model-based estimation of ancestry in unrelated individuals. *Genome Res* 19:1655-1664.

Anderson CN, Ramakrishnan U, Chan YL, Hadly EA. 2005. Serial SimCoal: a population genetics model for data from multiple populations and points in time. *Bioinformatics* 21:1733-1734.

Andrews RM, Kubacka I, Chinnery PF, Lightowlers RN, Turnbull DM, Howell N. 1999. Reanalysis and revision of the Cambridge reference sequence for human mitochondrial DNA. *Nat Genet* 23:147-147.

Arias L, Barbieri C, Barreto G, Stoneking M, Pakendorf B. 2018. High-resolution mitochondrial DNA analysis sheds light on human diversity, cultural interactions, and population mobility in Northwestern Amazonia. *Am J Phys Anthropol* 165:238-255.

Auguie B. 2017. gridExtra: Miscellaneous Functions for "Grid" Graphics. R package version 2.3.

Bandelt H-J, Forster P, Rohl A. 1999. Median-Joining Networks for Inferring Intraspecific Phylogenies. *Mol Biol Evol* 16:37-48.

Barbieri C, Heggarty P, Castri L, Luiselli D, Pettener D. 2011. Mitochondrial DNA variability in the Titicaca basin: Matches and mismatches with linguistics and ethnohistory. *Am J Hum Biol* 23:89-99.

Beaumont MA, Zhang W, Balding DJ. 2002. Approximate Bayesian computation in population genetics. *Genetics* 162:2025-2035.

Benn-Torres J, Vilar MG, Torres GA, Gaieski JB, Bharath Hernandez R, Browne ZE, Stevenson M, Walters W, Schurr TG, Genographic C. 2015. Genetic Diversity in the Lesser Antilles and Its Implications for the Settlement of the Caribbean Basin. *PLoS One* 10:e0139192.

Bodner M, Perego UA, Huber G, Fendt L, Rock AW, Zimmermann B, Olivieri A, Gomez-Carballa A, Lancioni H, Angerhofer N, et al. 2012. Rapid coastal spread of First Americans: novel insights from South America's Southern Cone mitochondrial genomes. *Genome Res* 22:811-820.

Brandini S, Bergamaschi P, Cerna MF, Gandini F, Bastaroli F, Bertolini E, Cereda C, Ferretti L, Gómez-Carballa A, Battaglia V, et al. 2018. The Paleo-Indian Entry into South America According to Mitogenomes. *Mol Biol Evol* 35:299-311.

Cardoso S, Alfonso-Sanchez MA, Valverde L, Sanchez D, Zarrabeitia MT, Odriozola A, Martinez-Jarreta B, de Pancorbo MM. 2012. Genetic uniqueness of the Waorani tribe from the Ecuadorian Amazon. *Heredity* 108:609-615.

Carpenter ML, Buenrostro JD, Valdiosera C, Schroeder H, Allentoft ME, Sikora M, Rasmussen M, Gravel S, Guillen S, Nekhrizov G, et al. 2013. Pulling out the 1%: whole-genome capture for the targeted enrichment of ancient DNA sequencing libraries. *Am J Hum Genet* 93:852-864.

Cooper A, Poinar H. 2000. Ancient DNA: Do It Right or Not at All. *Science* 289:1139.

Crespo-Torres E. 2000. Estudio Comparativo Biocultural entre dos Poblaciones Prehistóricas en la Isla Puerto Rico: Punta Candelero y Paso del Indio. [PhD Thesis]: Universidad Nacional Autónoma de México.

Cui Y, Lindo J, Hughes CE, Johnson JW, Hernandez AG, Kemp BM, Ma J, Cunningham R, Petzelt B, Mitchell J, et al. 2013. Ancient DNA analysis of mid-holocene individuals from the Northwest Coast of North America reveals different evolutionary paths for mitogenomes. *PLoS One* 8:e66948.

Curet LA, Stringer LM. 2010. Tibes: People, Power, and Ritual at the Center of the Cosmos. Tuscaloosa: University of Alabama Press.

Dabney J, Knapp M, Glocke I, Gansauge MT, Weihmann A, Nickel B, Valdiosera C, Garcia N, Pääbo S, Arsuaga JL, et al. 2013. Complete mitochondrial genome sequence of a Middle Pleistocene cave bear reconstructed from ultrashort DNA fragments. *Proc Natl Acad Sci U S A* 110:15758-15763.

de Saint Pierre M, Bravi CM, Motti JM, Fuku N, Tanaka M, Llop E, Bonatto SL, Moraga M. 2012. An alternative model for the early peopling of southern South America revealed by analyses of three mitochondrial DNA haplogroups. *PLoS One* 7:e43486.

Drummond AJ, Suchard MA, Xie D, Rambaut A. 2012. Bayesian Phylogenetics with BEAUti and the BEAST 1.7. *Mol Biol Evol* 29:1969-1973.

Duchêne S, Duchêne D, Holmes EC, Ho SYW. 2015. The performance of the date-randomization test in phylogenetic analyses of time-structured virus data. *Mol Biol Evol* 32:1895-1906.

Duggan AT, Harris AJT, Marciniak S, Marshall I, Kuch M, Kitchen A, Renaud G, Southon J, Fuller B, Young J, et al. 2017. Genetic Discontinuity between the Maritime Archaic and Beothuk Populations in Newfoundland, Canada. *Curr Biol* 27:3149-3156 e3111.

Excoffier L, Novembre J, Schneider S. 2000. SIMCOAL: a general coalescent program for the simulation of molecular data in interconnected populations with arbitrary demography. *J Hered* 91:506-509.

Excoffier L, Smouse PE, Quattro JM. 1992. Analysis of molecular variance inferred from metric distances among DNA haplotypes: application to human mitochondrial DNA restriction data. *Genetics* 131:479.

Fagundes NJ, Kanitz R, Eckert R, Valls AC, Bogo MR, Salzano FM, Smith DG, Silva WA, Jr., Zago MA, Ribeiro-dos-Santos AK, et al. 2008. Mitochondrial population genomics supports a single pre-Clovis origin with a coastal route for the peopling of the Americas. *Am J Hum Genet* 82:583-592.

Fontanez R. 1991. Restos Faunisticos y Explotacion del Medioambiente en Punta Candelero, Puerto Rico. Informe Preliminar. Proceedings of the 13th International Congress for Caribbean Archaeology; Curaçao.

Fu Q, Mittnik A, Johnson PL, Bos K, Lari M, Bollongino R, Sun C, Giemsch L, Schmitz R, Burger J, et al. 2013. A revised timescale for human evolution based on ancient mitochondrial genomes. *Curr Biol* 23:553-559.

Gilbert MTP, Hansen AJ, Willerslev E, Turner-Walker G, Collins M. 2006. Insights into the processes behind the contamination of degraded human teeth and bone samples with exogenous sources of DNA. *Int J Osteoarchaeol* 16:156-164.

Ginolhac A, Rasmussen M, Gilbert MTP, Willerslev E, Orlando L. 2011. mapDamage: testing for damage patterns in ancient DNA sequences. *Bioinformatics* 27:2153-2155.

Gnecchi-Ruscone GA, Sarno S, De Fanti S, Gianvincenzo L, Giuliani C, Boattini A, Bortolini E, Di Corcia T, Mellado CS, Dávila Francia TJ, et al. 2019. Dissecting the Pre-Columbian Genomic Ancestry of Native Americans along the Andes–Amazonia Divide. *Mol Biol Evol* 36:1254-1269.

Heled J, Drummond AJ. 2008. Bayesian inference of population size history from multiple loci. *BMC Evol Biol* 8:289.

Ho SY, Endicott P. 2008. The crucial role of calibration in molecular date estimates for the peopling of the Americas. *Am J Hum Genet* 83:142-146; author reply 146-147.

Ho SY, Shapiro B. 2011. Skyline-plot methods for estimating demographic history from nucleotide sequences. *Mol Ecol Resour* 11:423-434.

Hutt S, Blanco CM, Varmer O. 1999. Heritage Resources Law: Protecting the Archeological and Cultural Environment. USA: John Wiley & Sons.

Ingman M, Kaessmann H, Paabo S, Gyllensten U. 2000. Mitochondrial genome variation and the origin of modern humans. *Nature* 408:708-713.

Jónsson H, Gonolhac A, Schubert M, Johnson P, Orlando L. 2013. mapDamage2.0: fast approximate Bayesian estimates of ancient DNA damage parameters. *Bioinformatics* 29:1682-1684.

Jørsboe E, Hanghøj K, Albrechtsen A. 2017. fastNGSadmix: admixture proportions and principal component analysis of a single NGS sample. *Bioinformatics* 33:3148-3150.

Kashani BK, Perego UA, Olivieri A, Angerhofer N, Gandini F, Carossa V, Lancioni H, Semino O, Woodward SR, Achilli A, et al. 2011. Mitochondrial haplogroup C4c: A rare lineage entering America through the ice-free corridor? *Am J Phys Anthropol* 147:35-39.

Kassambara A, Mundt F. 2017. factoextra: Extract and Visualize the Results of Multivariate Data Analyses. R package version 1.0.5.

Katoh K, Standley DM. 2013. MAFFT Multiple Sequence Alignment Software Version 7: Improvements in Performance and Usability. *Mol Biol Evol* 30:772-780.

Kimura M. 1980. A simple method for estimating evolutionary rates of base substitutions through comparative studies of nucleotide sequences. *J Mol Evol* 16:111-120.

Kircher M, Sawyer S, Meyer M. 2012. Double indexing overcomes inaccuracies in multiplex sequencing on the Illumina platform. *Nucleic Acids Res* 40:e3.

Korneliussen TS, Albrechtsen A, Nielsen R. 2014. ANGSD: Analysis of next generation sequencing data. *BMC Bioinform* 15.

Kumar S, Bellis C, Zlojutro M, Melton PE, Blangero J, Curran JE. 2011. Large scale mitochondrial sequencing in Mexican Americans suggests a reappraisal of Native American origins. *BMC Evol Biol* 11.

Lalueza-Fox C, Calderón FL, Calafell F, Morera B, Bertranpetit J. 2001. MtDNA from extinct Tainos and the peopling of the Caribbean. *Ann Hum Genet* 65:137-151.

Lalueza-Fox C, Gilbert MTP, Martinez-Fuentes AJ, Calafell F, Bertranpetit J. 2003. Mitochondrial DNA from pre-Columbian Ciboneys from Cuba and the prehistoric colonization of the Caribbean. *Am J Phys Anthropol* 121:97-108.

Lanfear R, Calcott B, Ho SY, Guindon S. 2012. Partitionfinder: combined selection of partitioning schemes and substitution models for phylogenetic analyses. *Mol Biol Evol* 29:1695-1701.

Lazaridis I, Patterson N, Mittnik A, Renaud G, Mallick S, Kirsanow K, Sudmant PH, Schraiber JG, Castellano S, Lipson M, et al. 2014. Ancient human genomes suggest three ancestral populations for present-day Europeans. *Nature* 513:409-413.

Lee EJ, Merriwether DA. 2015. Identification of Whole Mitochondrial Genomes from Venezuela and Implications on Regional Phylogenies in South America. *Hum Biol* 87:29-38.

Leigh J, Bryant D. 2015. PopART: Full-feature software for haplotype network construction. *Methods Ecol Evol* 6:1110-1116.

Li H, Durbin R. 2009. Fast and accurate short read alignment with Burrows–Wheeler transform. *Bioinformatics* 25:1754-1760.

Li H, Handsaker B, Wysoker A, Fennell T, Ruan J, Homer N, Marth G, Abecasis G, Durbin R, Genome Project Data Processing S. 2009. The Sequence Alignment/Map format and SAMtools. *Bioinformatics* 25:2078-2079.

Lippold S, Xu H, Ko A, Li M, Renaud G, Butthof A, Schroder R, Stoneking M. 2014. Human paternal and maternal demographic histories: insights from high-resolution Y chromosome and mtDNA sequences. *Investig Genet* 5:13.

Lischer HEL, Excoffier L. 2012. PGDSpider: an automated data conversion tool for connecting population genetics and genomics programs. *Bioinformatics* 28:298-299.

Llamas B, Fehren-Schmitz L, Valverde G, Soubrier J, Mallick S, Rohland N, Nordenfelt S, Valdiosera C, Richards SM, Rohrlach A, et al. 2016. Ancient mitochondrial DNA provides high-resolution time scale of the peopling of the Americas. *Sci Adv* 2:e1501385.

Llamas B, Valverde G, Fehren-Schmitz L, Weyrich LS, Cooper A, Haak W. 2017. From the field to the laboratory: Controlling DNA contamination in human ancient DNA research in the high-throughput sequencing era. *Sci Tech Arch Resear* 3:1-14.

Mallick S, Li H, Lipson M, Mathieson I, Gymrek M, Racimo F, Zhao M, Chennagiri N, Nordenfelt S, Tandon A, et al. 2016. The Simons Genome Diversity Project: 300 genomes from 142 diverse populations. *Nature* 538:201-206.

Maricic T, Whitten M, Paabo S. 2010. Multiplexed DNA sequence capture of mitochondrial genomes using PCR products. *PLoS One* 5:e14004.

Mendisco F, Pemonge MH, Leblay E, Romon T, Richard G, Courtaud P, Deguilloux MF. 2015. Where are the Caribs? Ancient DNA from ceramic period human remains in the Lesser Antilles. *Philos Trans R Soc Lond B Biol Sci* 370:20130388.

Mendizabal I, Sandoval K, Berniell-Lee G, Calafell F, Salas A, Martinez-Fuentes A, Comas D. 2008. Genetic origin, admixture, and asymmetry in maternal and paternal human lineages in Cuba. *BMC Evol Biol* 8:213.

Meyer M, Kircher M. 2010. Illumina sequencing library preparation for highly multiplexed target capture and sequencing. *Cold Spring Harb Protoc* 6:1-10.

Meyer M, Kircher M, Gansauge MT, Li H, Racimo F, Mallick S, Schraiber JG, Jay F, Prufer K, de Filippo C, et al. 2012. A high-coverage genome sequence from an archaic Denisovan individual. *Science* 338:222-226.

Meyer S, Weiss G, Von Haeseler A. 1999. Pattern of nucleotide substitution and rate heterogeneity in the hypervariable regions I and II of human mtDNA. *Genetics* 152:1103-1110.

Milne I, Stephen G, Bayer M, Cock PJA, Pritchard L, Cardle L, Shaw PD, Marshall D. 2013. Using Tablet for visual exploration of second-generation sequencing data. *Brief Bioinform* 14:193-202.

Mizuno F, Wang L, Sugiyama S, Kurosaki K, Granados J, Gomez-Trejo C, Acuna-Alonzo V, Ueda S. 2017. Characterization of complete mitochondrial genomes of indigenous Mayans in Mexico. *Ann Hum Biol* 44:652-658.

Molak M, Lorenzen ED, Shapiro B, Ho SYW. 2013. Phylogenetic Estimation of Timescales Using Ancient DNA: The Effects of Temporal Sampling Scheme and Uncertainty in Sample Ages. *Mol Biol Evol* 30:253-262.

Molak M, Suchard MA, Ho SYW, Beilman DW, Shapiro B. 2015. Empirical calibrated radiocarbon sampler: a tool for incorporating radiocarbon-date and calibration error into Bayesian phylogenetic analyses of ancient DNA. *Mol Ecol Resour* 15:81-86.

Nei M. 1987. Molecular Evolutionary Genetics. New York: Columbia University Press.

Nei M, Miller J. 1990. A simple method for estimating average number of nucleotide substitutions within and between populations from restriction data. *Genetics* 125:873-879.

Nieves-Colón MA. 2012. The Contribution of sub-Saharan African and Eurasian maternal (mtDNA) lineages to the genetic heritage of the Dominican Republic. [M.A. Thesis]: Arizona State University.

Nieves-Colón MA, Ozga AT, Pestle WJ, Cucina A, Tiesler V, Stanton TW, Stone AC. 2018. Comparison of two ancient DNA extraction protocols for skeletal remains from tropical environments. *Am J Phys Anthropol* 166:824-836.

O'Fallon BD, Fehren-Schmitz L. 2011. Native Americans experienced a strong population bottleneck coincident with European contact. *Proc Natl Acad Sci U S A* 108:20444-20448.

Okonechnikov K, Conesa A, García-Alcalde F. 2016. Qualimap 2: advanced multi-sample quality control for high-throughput sequencing data. *Bioinformatics* 32:292-294.

Oksanen J, Blanchet FG, Friendly M, Kindt R, Legendre P, McGlinn D, Minchin PR, O'Hara RB, Simpson GL, Solymons P, et al. 2018. vegan: Community Ecology Package. R package version 2.5-2.

Ousley SD, Billeck WT, Hollinger RE. 2005. Federal repatriation legislation and the role of physical anthropology in repatriation. *Am J Phys Anthropol* Suppl 41:2-32.

Ozga AT, Nieves-Colón MA, Honap TP, Sankaranarayanan K, Hofman CA, Milner GR, Lewis CM, Jr., Stone AC, Warinner C. 2016. Successful enrichment and recovery of whole mitochondrial genomes from ancient human dental calculus. *Am J Phys Anthropol* 160:220-228.

Paradis E. 2010. pegas: an R package for population genetics with an integrated-modular approach. *Bioinformatics* 26:419-420.

Paradis E, Claude J, Strimmer K. 2004. APE: analyses of phylogenetics and evolution in R language. *Bioinformatics* 20:289-290.

Patterson N, Moorjani P, Luo Y, Mallick S, Rohland N, Zhan Y, Genschoreck T, Webster T, Reich D. 2012. Ancient Admixture in Human History. *Genetics* 192:1065.

Patterson N, Price AL, Reich D. 2006. Population structure and eigenanalysis. *PLoS Genet* 2:e190.

Perego UA, Achilli A, Angerhofer N, Accetturo M, Pala M, Olivieri A, Hooshiar Kashani B, Ritchie KH, Scozzari R, Kong QP, et al. 2009. Distinctive Paleo-Indian migration routes from Beringia marked by two rare mtDNA haplogroups. *Curr Biol* 19:1-8.

Perego UA, Angerhofer N, Pala M, Olivieri A, Lancioni H, Hooshiar Kashani B, Carossa V, Ekins JE, Gomez-Carballa A, Huber G, et al. 2010. The initial peopling of the Americas: a growing number of founding mitochondrial genomes from Beringia. *Genome Res* 20:1174-1179.

Pestle WJ. 2010. Diet and Society in Prehistoric Puerto Rico: An Isotopic Approach. [Ph.D. Thesis]: University of Illinois at Chicago.

Pestle WJ, Colvard M. 2012. Bone collagen preservation in the tropics: a case study from ancient Puerto Rico. *J Archaeol Sci* 39:2079-2090.

Posada D, Buckley TR. 2004. Model selection and model averaging in phylogenetics: advantages of akaike information criterion and bayesian approaches over likelihood ratio tests. *Syst Biol* 53:793-808.

Purcell S, Neale B, Todd-Brown K, Thomas L, Ferreira MAR, Bender D, Maller J, Sklar P, de Bakker PIW, Daly MJ, et al. 2007. PLINK: A Tool Set for Whole-Genome Association and Population-Based Linkage Analyses. *Am J Hum Genet* 81:559-575.

Raghavan M, Skoglund P, Graf KE, Metspalu M, Albrechtsen A, Moltke I, Rasmussen S, Stafford TW, Jr., Orlando L, Metspalu E, et al. 2014. Upper Palaeolithic Siberian genome reveals dual ancestry of Native Americans. *Nature* 505:87-91.

Raghavan M, Steinrucken M, Harris K, Schiffels S, Rasmussen S, DeGiorgio M, Albrechtsen A, Valdiosera C, Avila-Arcos MC, Malaspinas AS, et al. 2015. Genomic evidence for the Pleistocene and recent population history of Native Americans. *Science* 349:aab3884.

Renaud G, Slon V, Duggan AT, Kelso J. 2015. Schmutzi: estimation of contamination and endogenous mitochondrial consensus calling for ancient DNA. *Genome Biol* 16:224.

Rodríguez López M. 1991. Arqueología de Punta Candelero, Puerto Rico. Proceedings of the 13th International Congress for Caribbean Archaeology; Curaçao. p. 605-627.

Rohland N, Hofreiter M. 2007. Ancient DNA extraction from bones and teeth. *Nat Protoc* 2:1756-1762.

Rousset F. 2008. genepop'007: a complete re-implementation of the genepop software for Windows and Linux. *Mol Ecol Resour* 8:103-106.

Sarkar D. 2008. Lattice: Multivariate Data Visualization with R. New York: Springer.

Schroeder H, Sikora M, Gopalakrishnan S, Cassidy LM, Maisano Delser P, Sandoval Velasco M, Schraiber JG, Rasmussen S, Homburger JR, Avila-Arcos MC, et al. 2018. Origins and genetic legacies of the Caribbean Taino. *Proc Natl Acad Sci U S A* 115:2341-2346.

Schubert M, Ermini L, Der Sarkissian C, Jonsson H, Ginolhac A, Schaefer R, Martin MD, Fernandez R, Kircher M, McCue M, et al. 2014. Characterization of ancient and modern genomes by SNP detection and phylogenomic and metagenomic analysis using PALEOMIX. *Nat Protoc* 9:1056-1082.

Schuenemann VJ, Bos K, DeWitte S, Schmedes S, Jamieson J, Mittnik A, Forrest S, Coombes BK, Wood JW, Earn DJD, et al. 2011. Targeted enrichment of ancient pathogens yielding the pPCP1 plasmid of Yersinia pestis from victims of the Black Death. *Proc Natl Acad Sci U S A* 108:E746–E752.

Seguin-Orlando A, Hoover CA, Vasiliev SK, Ovodov ND, Shapiro B, Cooper A, Rubin EM, Willerslev E, Orlando L. 2015. Amplification of TruSeq ancient DNA libraries with AccuPrime Pfx: consequences on nucleotide misincorporation and methylation patterns. *Sci Tech Arch Resear* 1:1-9.

Siegel P. 2011. Puerto Rico. In: Siegel P, Righter E, editors. Protecting Heritage in the Caribbean. Tuscaloosa: University of Alabama Press. p. 46-57.

Simbolo M, Gottardi M, Corbo V, Fassan M, Mafficini A, Malpeli G, Lawlor RT, Scarpa A. 2013. DNA qualification workflow for next generation sequencing of histopathological samples. *PLoS One* 8:e62692.

Skoglund P, Northoff BH, Shunkov MV, Derevianko AP, Paabo S, Krause J, Jakobsson M. 2014. Separating endogenous ancient DNA from modern day contamination in a Siberian Neandertal. *Proc Natl Acad Sci U S A* 111:2229-2234.

Skoglund P, Storå J, Götherström A, Jakobsson M. 2013. Accurate sex identification of ancient human remains using DNA shotgun sequencing. *J Archaeol Sci* 40:4477-4482.

Soares P, Ermini L, Thomson N, Mormina M, Rito T, Rohl A, Salas A, Oppenheimer S, Macaulay V, Richards MB. 2009. Correcting for purifying selection: an improved human mitochondrial molecular clock. *Am J Hum Genet* 84:740-759.

Söchtig J, Álvarez-Iglesias V, Mosquera-Miguel A, Gelabert-Besada M, Gómez-Carballa A, Salas A. 2015. Genomic insights on the ethno-history of the Maya and the 'Ladinos' from Guatemala. *BMC Genomics* 16:131.

Tamm E, Kivisild T, Reidla M, Metspalu M, Smith DG, Mulligan CJ, Bravi CM, Rickards O, Martinez-Labarga C, Khusnutdinova EK, et al. 2007. Beringian Standstill and Spread of Native American Founders. *PLoS One* 2:e829.

Tascón Peñaranda EP. 2014. Relación entre los principales linajes indígenas del ADN mitochondrial en las islas de Puerto Rico y República Dominicana. [M.A. Thesis]: Universidad de Puerto Rico, Recinto de Mayagüez.

van Oven M, Kayser M. 2009. Updated comprehensive phylogenetic tree of global human mitochondrial DNA variation. *Hum Mutat* 30:E386-E394.

Vilar MG, Melendez C, Sanders AB, Walia A, Gaieski JB, Owings AC, Schurr TG, Genographic C. 2014. Genetic diversity in Puerto Rico and its implications for the peopling of the Island and the West Indies. *Am J Phys Anthropol* 155:352-368.

Walker J. 2005. The Paso del Indio Site, Vega Baja, Puerto Rico: A Progress Report. In: Siegel P, editor. Ancient Borinquen: Archaeology and Ethnohistory of Native Puerto Rico. Tuscaloosa: University of Alabama Press. p. 55-87.

Warinner C, Rodrigues JF, Vyas R, Trachsel C, Shved N, Grossmann J, Radini A, Hancock Y, Tito RY, Fiddyment S, et al. 2014. Pathogens and host immunity in the ancient human oral cavity. *Nat Genet* 46:336-344.

Warnes GR, Bolker B, Bonebakker L, Gentleman R, Huber W, Liaw A, Lumley T, Maechler M, Magnusson A, Moeller S, et al. 2016. gplots: Various R Programming Tools for Plotting Data. R package version 3.0.1.

Wei T, Simko V. 2017. "corrplot": Visualization of a Correlation Matrix. R package version 0.84.

Weissensteiner H, Pacher D, Kloss-Brandstätter A, Forer L, Specht G, Bandelt H-J, Kronenberg F, Salas A, Schönherr S. 2016. HaploGrep 2: mitochondrial haplogroup classification in the era of high-throughput sequencing. *Nucleic Acids Res* 44:W58-W63.

Wickham H. 2009. ggplot2: Elegant Graphics for Data Analysis. New York: Springer-Verlag.

### Supplementary Figure Legends

**Supplementary Figure S1. Permits for destructive analysis.**

**Supplementary Figure S2. Documentation of human skeletal remains prior to destructive analysis.**

**Supplementary Figure S3. Simplified timeline of the pre-contact history of the Antilles demonstrating occupation periods of studied sites**. Site chronologies based on published sources. Dashed lines represent the non-continuous occupation of Paso del Indio. Red boxes encompass minimum and maximum median calibrated radiocarbon dates (cal A.D.) for sampled human skeletal remains.

**Supplementary Figure S4. Per site comparison of recovery rates for complete mtDNA data after enrichment.**

**Supplementary Figure S5. Fragment misincorporation plots for 45 mtDNA enriched libraries selected for analysis.** Samples where mapped and filtered BAM files were analyzed after combining across multiple sequencing runs are noted with the "comb" notation.

**Supplementary Figure S6. Fragment misincorporation plots for two whole genome enriched libraries selected for analysis of autosomal genotypes.** Mapped and filtered BAM files were analyzed after combining across multiple sequencing runs.

**Supplementary Figure S7. Fragment misincorporation plots for nine mtDNA enriched libraries before and after contamination filtering with PMDtools.**

**Supplementary Figure S8. Pairwise $\Phi_{ST}$ measures calculated with complete mtDNA comparing communities from Paso del Indio, Punta Candelero and Tibes.**

**Supplementary Figure S9. Mantel test z-statistic plot.** Mantel test comparing genetic versus temporal distance for 37 radiocarbon dated skeletal remains from PC-PR.

**Supplementary Figure S10. Molecular diversity measures calculated for PC-PR and comparative populations.** Complete mtDNA data (Americas): A) Nucleotide diversity and variance, B) Haplotype diversity and variance. HVR-1 mtDNA data (Caribbean): A) Nucleotide diversity and variance, B) Haplotype diversity and variance.

**Supplementary Figure S11. Pairwise $\Phi_{ST}$ measures calculated with complete mtDNA comparing PC-PR to 46 ancient and present-day populations from the Americas.**

**Supplementary Figure S12. Pairwse $\Phi_{ST}$ measures calculated with HVR-1 data comparing PC-PR to eight ancient and present-day Caribbean populations.**

**Supplementary Figure S13. Shepard diagram of linear fit for two non-metric multidimensional scaling (MDS) analyses plotting pairwise $\Phi_{ST}$ distances**. A) Complete mtDNA MDS, B) HVR-1 MDS.

**Supplementary Figure S14. Correspondence analysis of haplogroup frequencies between Caribbean populations (HVR-1 data).**

**Supplementary Figure S15. Median joining network of complete mtDNA diversity for haplogroup A.** Includes data from 519 ancient and present-day individuals from across the Americas. PC-PR shown in red. Puerto Ricans shown in lilac.

**Supplementary Figure S16. Median joining network of complete mtDNA diversity for haplogroup C.** Includes data from 456 ancient and present-day individuals from across the Americas. PC-PR shown in red. Puerto Ricans shown in lilac.

**Supplementary Figure S17. Median joining network of complete mtDNA diversity for haplogroup D.** Includes data from 423 ancient and present-day individuals from across the Americas. PC-PR shown in red. D1 sequences from present-day Puerto Ricans were not publicly available for inclusion in this analysis.

**Supplementary Figure S18. Median joining network of HVR-1 mtDNA diversity across the Caribbean.** Includes data from ancient and present-day individuals sampled in Puerto Rico, Cuba, Dominican Republic, Trinidad First People's Community, St. Vincent Garifuna and Guadeloupe. A) Haplogroup A: n=236, B) Haplogroup C: n=41, C) Haplogroup D: n=37.

**Supplementary Figure S19. Demographic models of Caribbean population history tested with BayeSSC simulations.** Models simulate the available HVR-1 data from four pre-contact Caribbean populations under eight possible demographic scenarios varying the amount and direction of inter-island gene flow. C = Cuba, DR = Dominican Republic, PR = Puerto Rico, G = Guadeloupe.

**Supplementary Figure S20. Median tree calculated with BEAST and including 127 complete mtDNA sequences from ancient and present-day Puerto Rico (Native American lineages only).** Branches leading to the main clades Hg A2, B2, C1, and D1 have support values of 1, all other branches have support values < 0.85.

**Supplementary Figure S21. Median tree with rate along branches highlighted.**

**Supplementary FigureS22: Posterior population size changes for All Sequences dataset (median: 3; 95% HPD: 2–5).** Constant population can be confidently rejected (0 is not in the 95% HPD).

**Supplementary Figure S23. BEAST analysis results for Haplogroup A dataset.** A) Posterior population size changes (median: 2; % HPD: 1–4) for haplogroup A. Constant population can be confidently rejected (0 is not in the 95% HPD). B) Extended Bayesian skyline plot of female effective population size for haplogroup A, based on a generation time of 25 years.

**Supplementary Figure S24 BEAST analysis results for Haplogroup C dataset.** Posterior population size changes (median: 3; % HPD: 0–4) for haplogroup C. Constant population cannot be confidently rejected (0 is in the 95% HPD), therefore the skyline reconstruction is meaningless.

**Supplementary Figure S25. ADMIXTURE analysis (K3-K7) with PI-420a, PI-51 (PC-PR) and PC537 (Bahamas) autosomal genotypes.** A) Including all overlapping sites, B) including only transitions.

**Supplementary Figure S26. Cross validation errors associated with clustering analysis for runs K3=K7.**

**Supplementary Figure S27. Principal components analyses (PCA) with autosomal genotypes.** Ancient samples are projected on eigenvectors calculated using reference panel individuals. Top panels: PCA comparing PI-420a, PI-51 (PC-PR) and PC537 (Bahamas) to world populations. A) Including all overlapping positions, and B) restricted to transitions only. Bottom panels: PCA comparing PI-420a, PI-51 (PC-PR) and PC537 (Bahamas) to Native American populations. C) Including all overlapping positions, and D) restricted to transitions only.

**Supplementary Figure S28. Outgroup $f_3$ statistics for two Paso del Indio individuals.** Statistics calculated in the form $f_3$(ancient, X; Yoruba)**.** Error bars correspond to 95% standard error estimates. Populations are coloured by region: Blue = North America and Mesoamerica, Green = South America, Red = Caribbean. A) $f_3$(PI-420A, X; Yoruba) with all overlapping sites, B) $f_3$(PI-420A, X; Yoruba) with just transitions, C) $f_3$(PI-51, X; Yoruba) with all overlapping sites, D) $f_3$(PI-51, X; Yoruba) with just transitions.

**CONSEJO PARA LA PROTECCION DEL PATRIMONIO ARQUEOLOGICO TERRESTRE DE PUERTO RICO**

*Adscrito al*
*Instituto de Cultura Puertorriqueña*

5 de mayo de 2004

Dr. Antonio Curet Salim
Field Museum
1400 S. Lake Shore Dr.
Chicago, IL 60605

Estimado doctor Curet:

El Consejo para al Protección del Patrimonio Arqueológico Terrestre de Puerto Rico en su Reunión Ordinaria del 4 de mayo de 2004, tuvo ante su consideración su petición para el traslado de muestras de esqueletos humanos con propósito de análisis.

El Consejo determinó conceder su petición, por lo que se autoriza el traslado de las muestras. Debe informar al Consejo la cantidad de procedencia de estas una vez haya realizado la selección.

Atentamente

Marisol Rodríguez Miranda
Directora

c. Miembros del Consejo

Calle San Sebastián #1 Apartado 9024184 San Juan de Puerto Rico 00902-4184 Tel: 721-6828 / 723-1798

**Supplementary Figure S1. Permits for destructive analysis.**

ESTADO LIBRE ASOCIADO DE PUERTO RICO

**AUTORIDAD DE CARRETERAS Y TRANSPORTACION**

29 de abril de 2004

Dr. L. Antonio Curet, Curador Auxiliar
Department of Anthropology
Field Museum of Natural History
1400 S Lake Shore Drive
Chicago IL 60605-2496

**PROYECTO ARQUEOLÓGICO PASO DEL INDIO**
**AC-220115**

Estimado doctor Curet:

Hago referencia a su carta del 29 de marzo de 2004. Incluyo, debidamente firmada, una copia de su carta autorizándole a realizar los estudios solicitados. Debe quedar claro que esta autorización está condicionada a que usted obtenga la autorización del Consejo para la Protección del Patrimonio Arqueológico Terrestre de Puerto Rico antes de poder transportar los materiales arqueológicos fuera del Estado Libre Asociado de Puerto Rico y que cumpla con todas las disposiciones legales y reglamentarias aplicables.

Agradeceré se mantenga en coordinación con el Plan. Roberto Vélez, de nuestra Oficina de Estudios Ambientales (teléfono (787) 764-9692), y con el Dr. Jeff Walker, del Servicio Forestal Federal.

Cordialmente,

Irma M. García
Directora
Área de Programación
y Estudios Especiales

6704/CGA/RVB/egn

Anexo

Autoridad de Carreteras y Transportación • Departamento de Transportación y Obras Públicas

Estado Libre Asociado de Puerto Rico
*Municipio Autónomo de Ponce*

**Secretaría de Cultura y Turismo**

Sra. Vangie Rivera
Directora

10 de mayo de 2004

Dr. L. Antonio Curet
Departamento de Antropologia
Field Museum
Chicago, Ill.

Estimado Dr. Curet:

Permítame saludarle en nombre de todos los que trabajamos en la Secretaría de Cultura y Turismo. Es para nosotros un gran orgullo el que haya considerado el Centro Ceremonial Indígena de Tibes como uno de los yacimientos que serán incluidos en el estudio arqueológico

**"Proposal for the Study of Migration and Subsistence in Four Ancient Sites in Puerto Rico through Bone Chemistry and Genetic Analysis"**

Sabe que puede contar con nuestros recursos, nuestra colección arqueológica y la ayuda incondicional de todo el personal que labora en el Centro Ceremonial Indígena de Tibes para el desarrollo de este proyecto.Estamos seguros que el mismo pasará a ser parte de las investigaciones arqueológicas de importancia dentro de la prehistoria de nuestra isla.

En nombre de nuestra Alcaldesa, Hon. Delis Castillo de Santiago, y de nuestros ciudadanos, le damos las gracias por el interés que siempre ha mostrado por el Centro Ceremonial Indígena de Tibes y esperamos darle la bienvenida al comienzo del proyecto. Favor de manifestarle nuestro saludo y respeto a los Doctores Edwin Crespo, Sloan Williams y al Sr. William Pestle, co-investigadores de este proyecto.

Nuevamente, gracias por su compromiso con nuestra cultura y especialmente con Tibes.

Cordialmente,

Vangie Rivera
Directora
Cultura y Turismo

**Página Cibernética: www.ponceweb.org Correo Electrónico:**

A) Tibes (4 samples): T-257, T-3, T-27, T-268

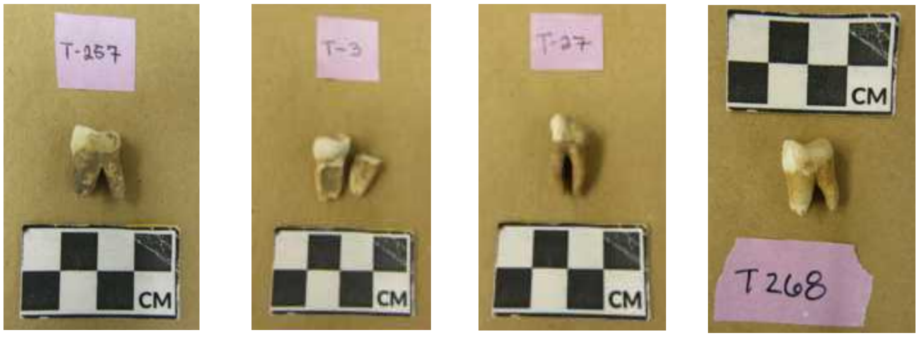


B) Paso del Indio (4 samples): PI-429, PI-410, PI-41, PI-433

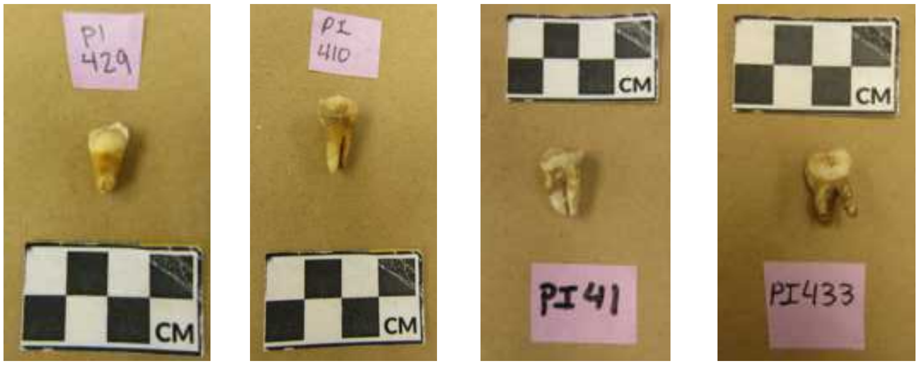


C) Punta Candelero (4 samples): PC-E36, PC-129, PC-E23, PC-A6E1

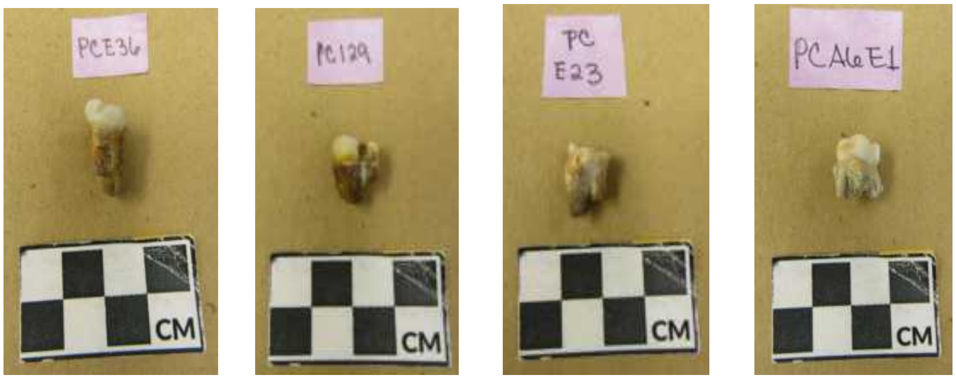


**Supplementary Figure S2. Documentation of human skeletal remains prior to destructive analysis.**

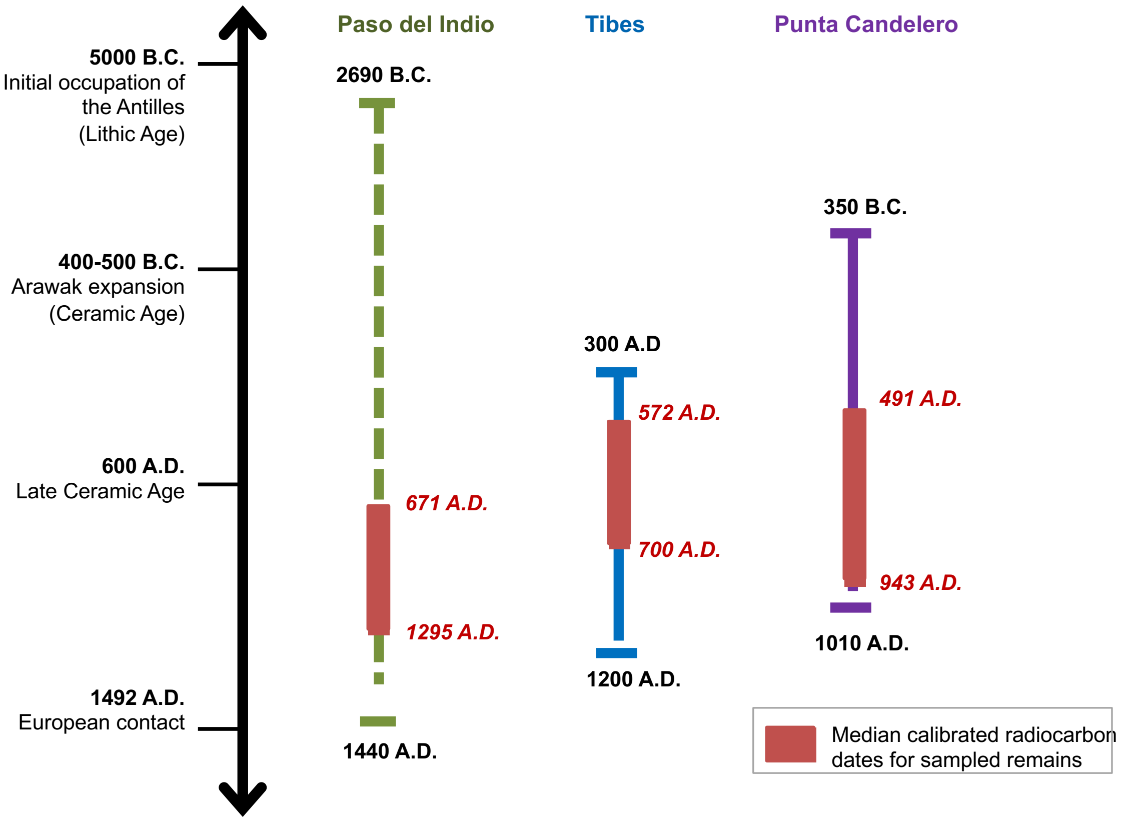


**Supplementary Figure S3. Simplified timeline of the pre-contact history of the Antilles demonstrating occupation periods of studied sites.** Site chronologies based on published sources. Dashed lines represent the non-continuous occupation of Paso del Indio. Red boxes encompass minimum and maximum median calibrated radiocarbon dates (cal A.D.) for sampled human skeletal remains.

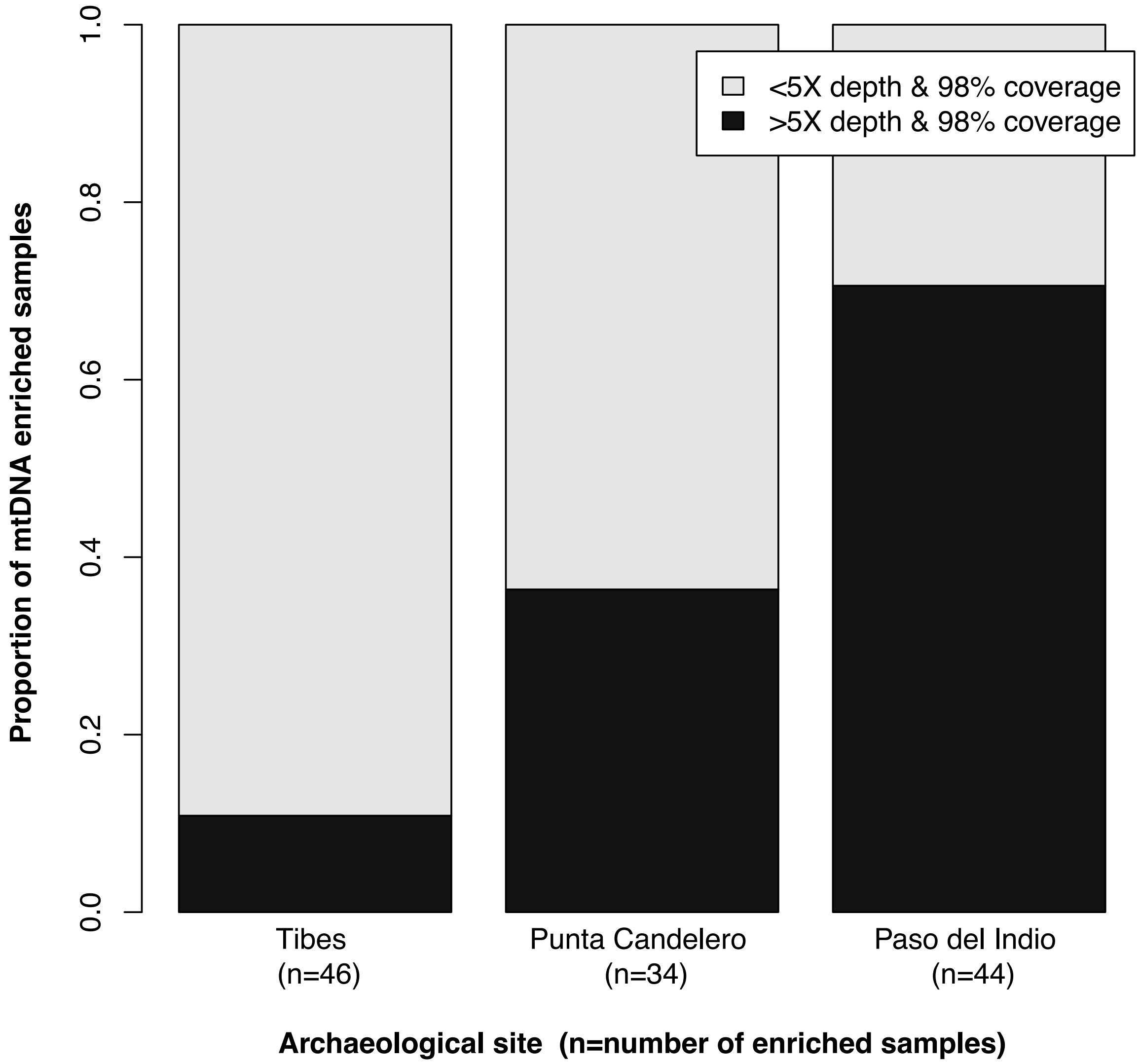


**Supplementary Figure S4. Per site comparison of recovery rates for complete mtDNA data after enrichment.**

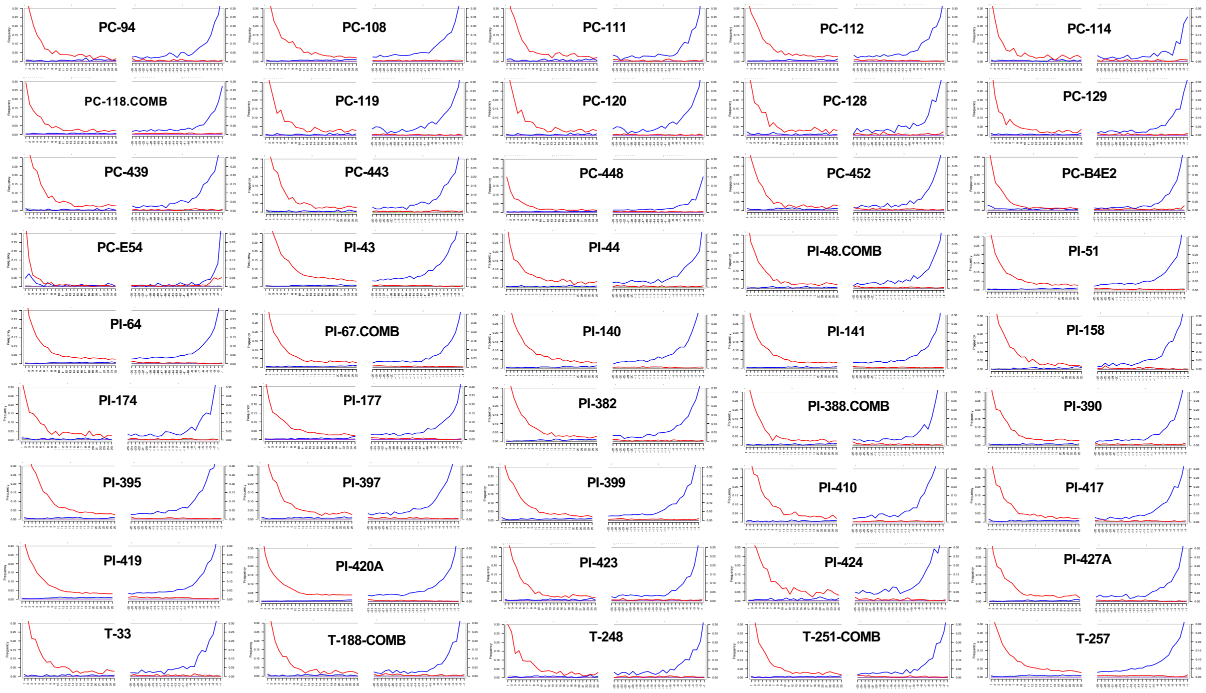


**Supplementary Figure S5.** Fragment misincorporation plots for 45 mtDNA enriched libraries selected for analysis. Samples where mapped and filtered BAM files were analyzed after combining across multiple sequencing runs are noted with the "comb" notation.

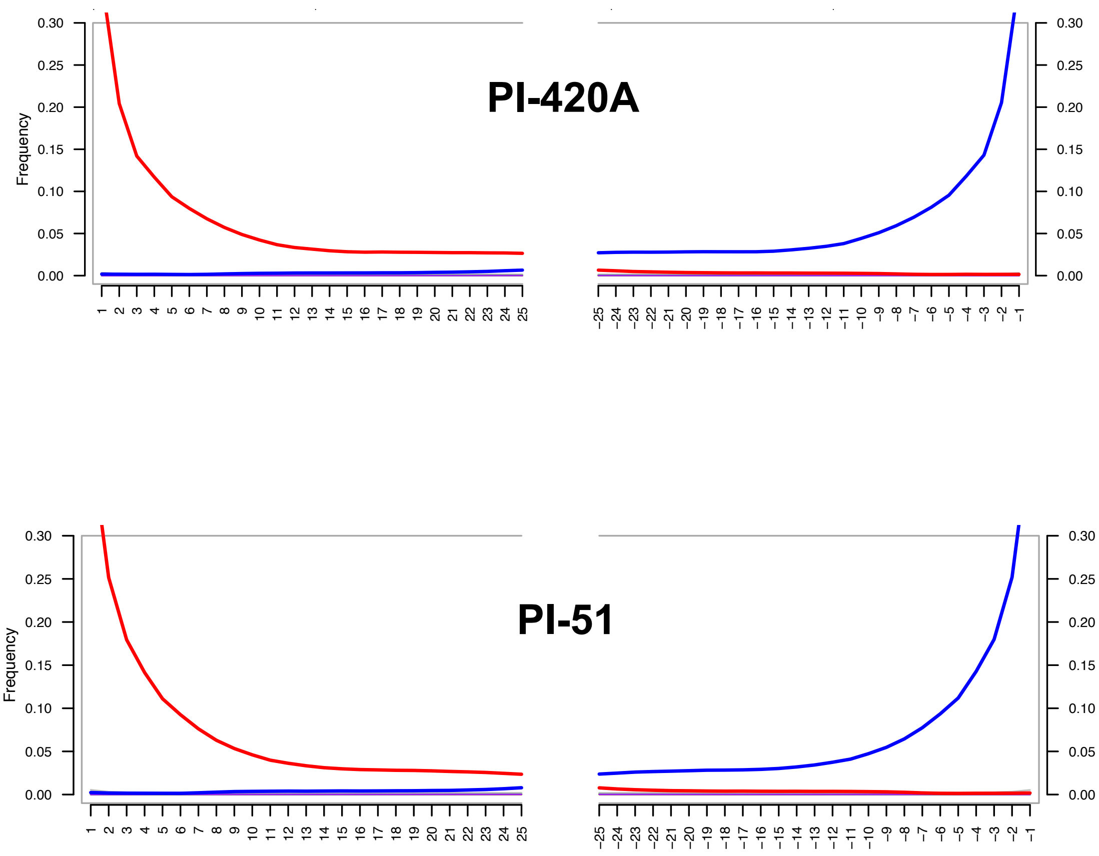


**Supplementary Figure S6.** Fragment misincorporation plots for two whole genome enriched libraries selected for analysis of autosomal genotypes. Mapped and filtered BAM files were analyzed after combining across multiple sequencing runs.

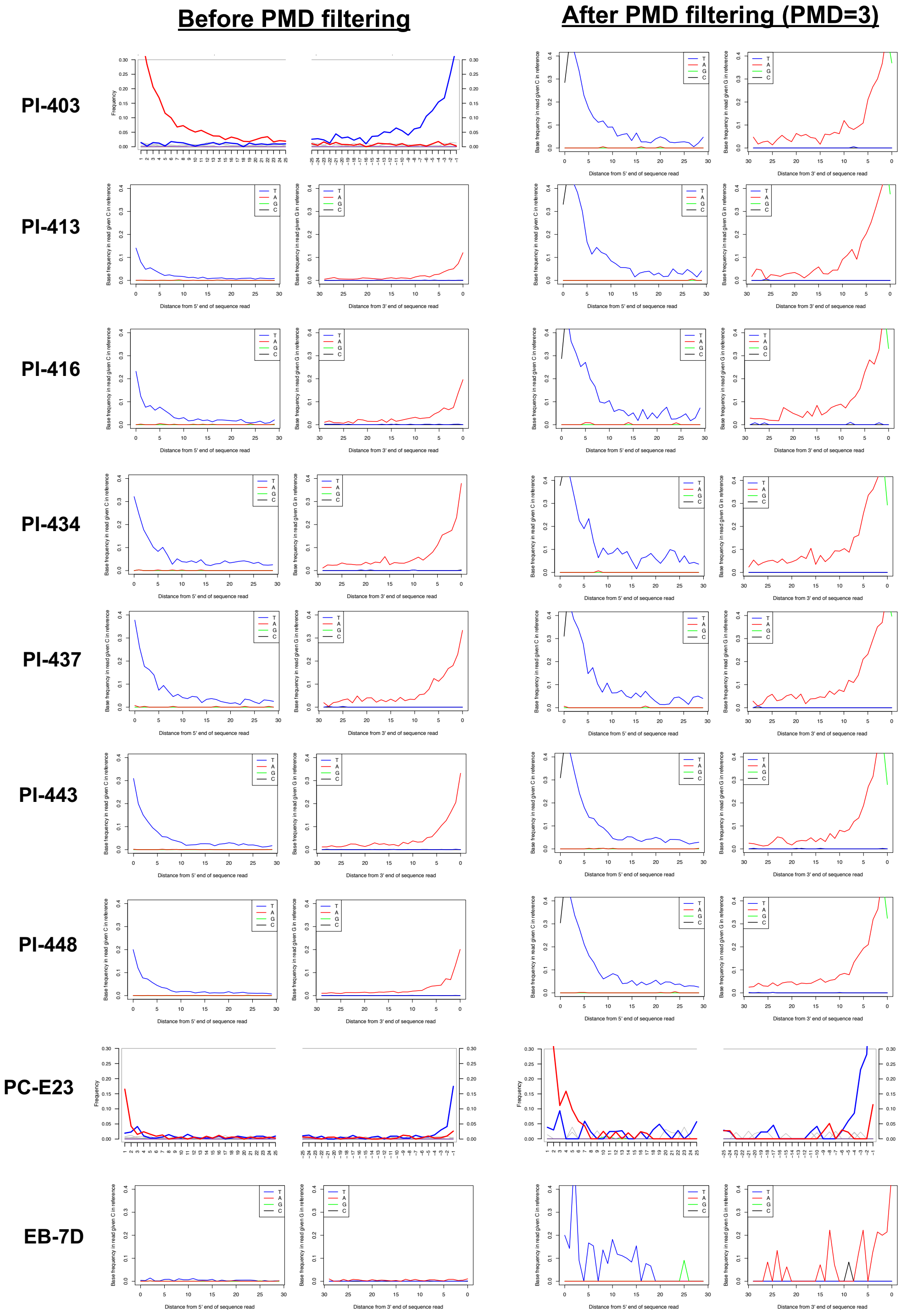


**Supplementary Figure S7. Fragment misincorporation plots for nine mtDNA enriched libraries before and after contamination filtering with PMDtools.**

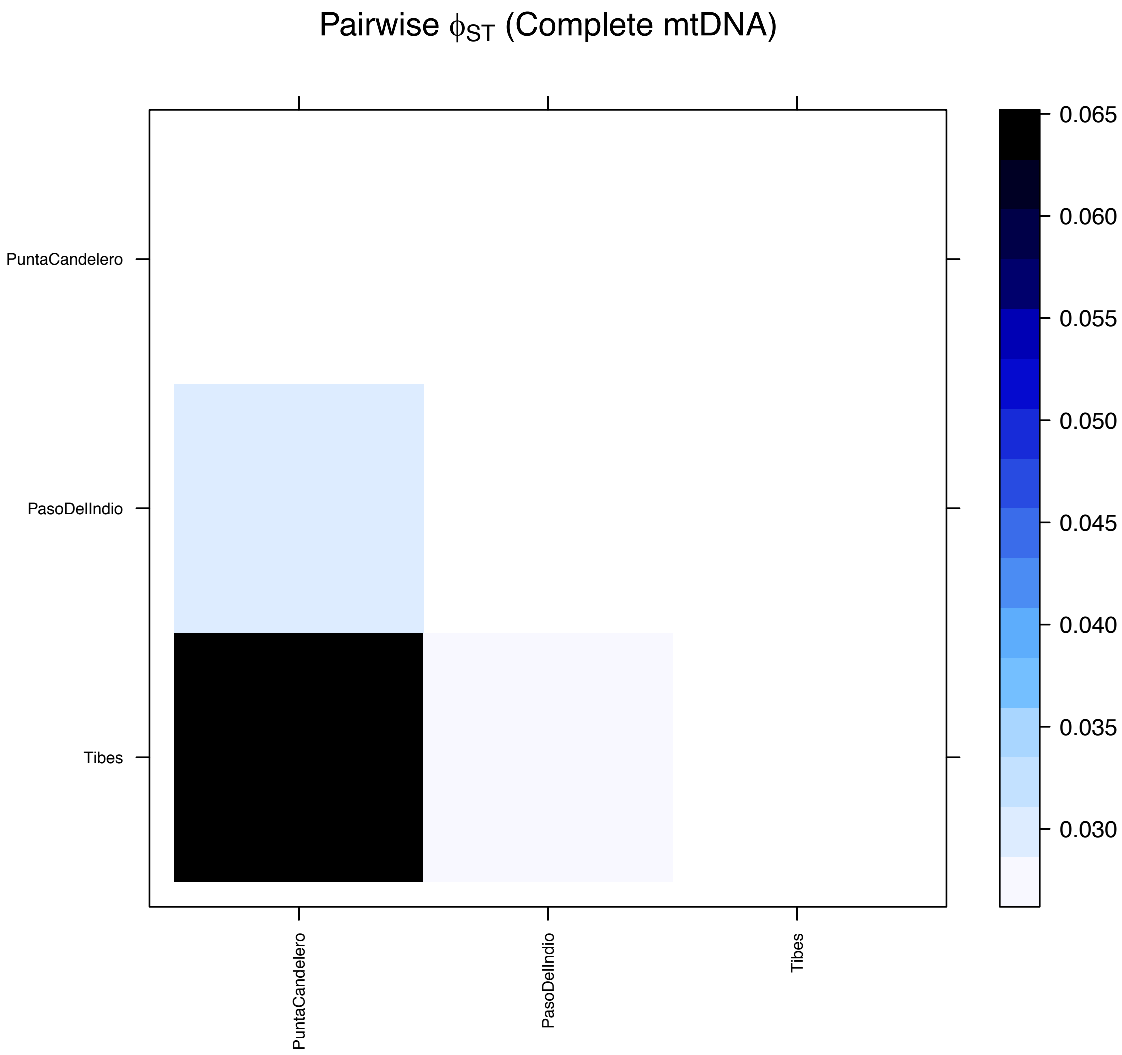


**Supplementary Figure S8. Pairwise $\Phi_{ST}$ measures calculated with complete mtDNA comparing communities from Paso del Indio, Punta Candelero and Tibes.**

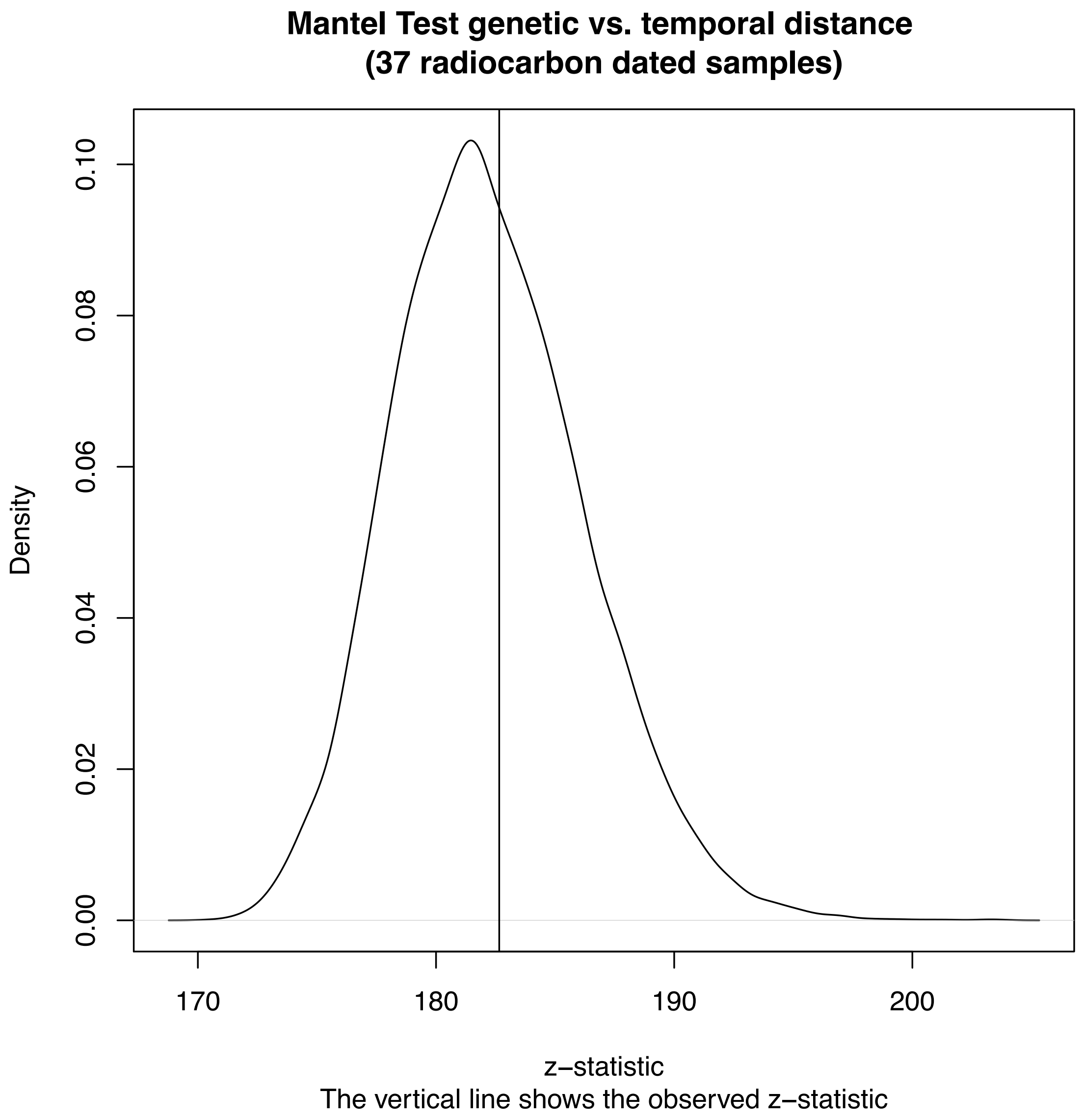


**Supplementary Figure S9. Mantel test z-statistic plot.** Mantel test comparing genetic versus temporal distance for 37 radiocarbon dated skeletal remains from PC-PR.

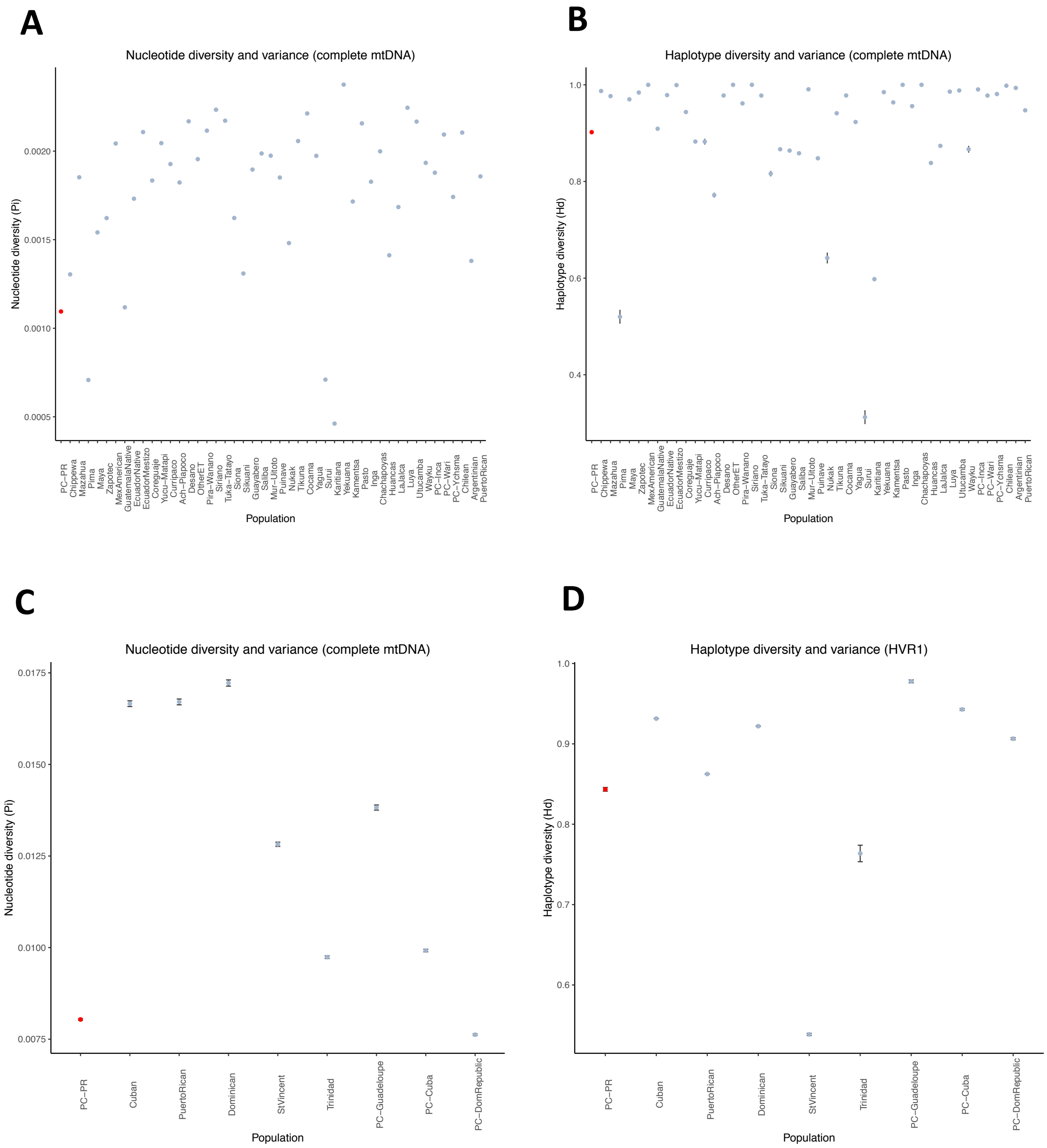


**Supplementary Figure S10. Molecular diversity measures calculated for PC-PR and comparative populations.** Complete mtDNA data (Americas): A) Nucleotide diversity and variance, B) Haplotype diversity and variance. HVR-1 mtDNA data (Caribbean): C) Nucleotide diversity and variance, D) Haplotype diversity and variance.

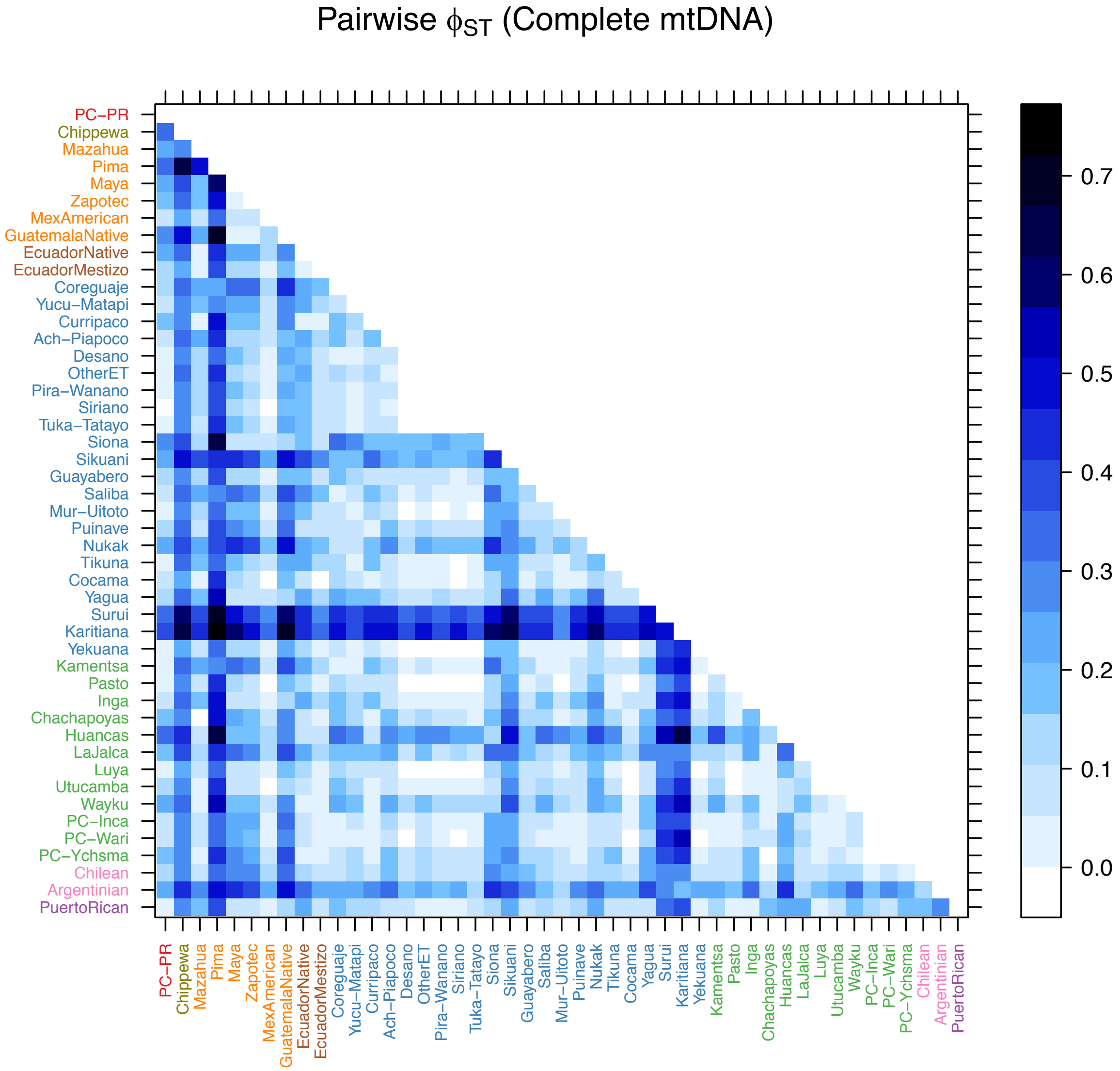


**Supplementary Figure S11. Pairwise $\Phi_{ST}$ measures calculated with complete mtDNA comparing PC-PR to 46 ancient and present-day populations from the Americas.**

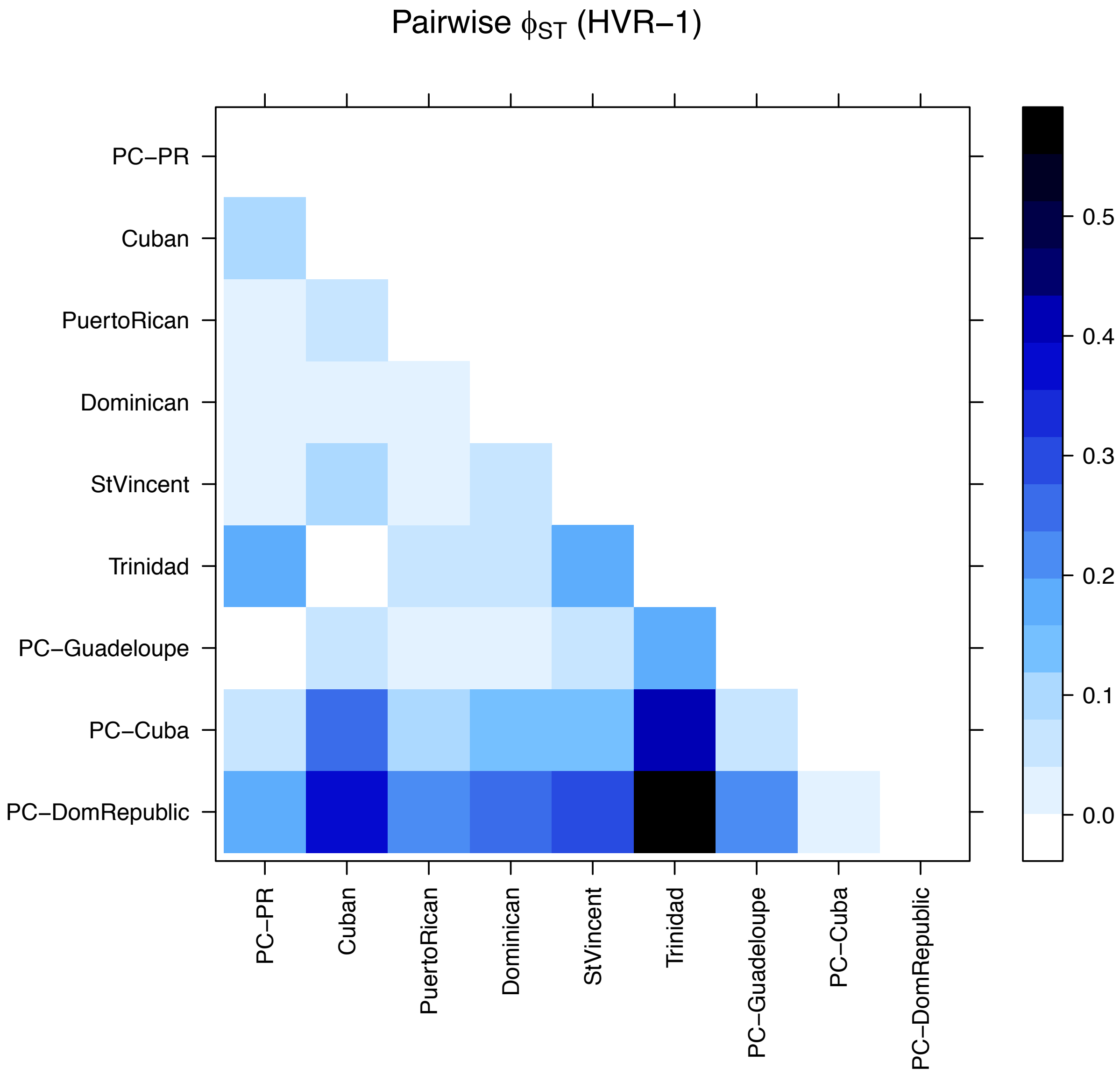


**Supplementary Figure S12. Pairwise $\Phi_{ST}$ measures calculated with HVR-1 data comparing PC-PR to eight ancient and present-day Caribbean populations.**

A

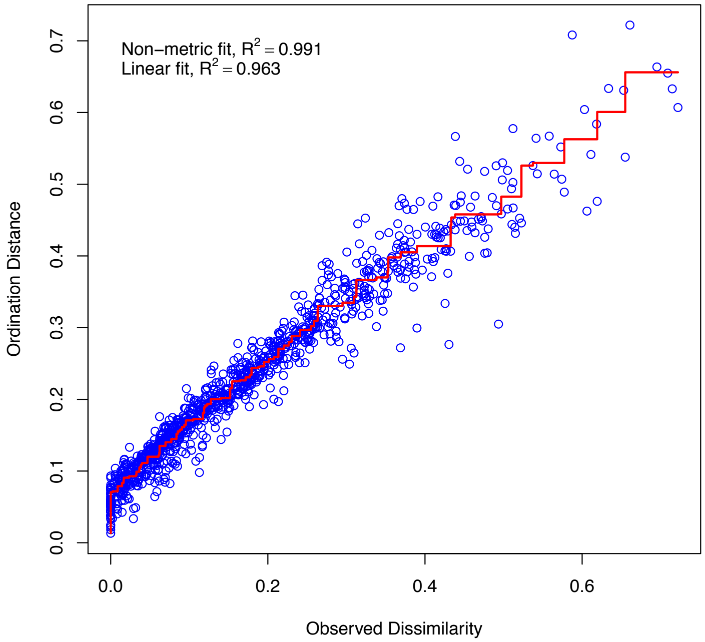


B

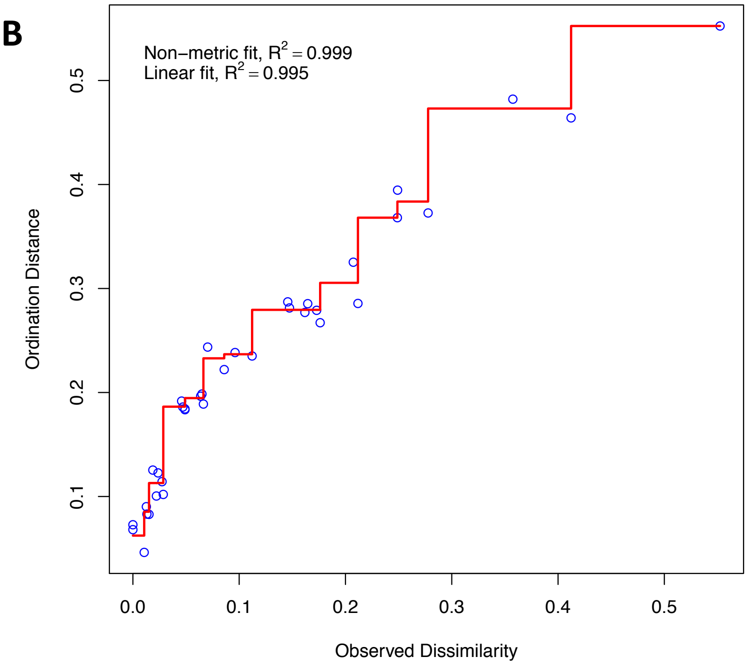


**Supplementary Figure S13. Shepard diagram of linear fit for two non-metric multidimensional scaling (MDS) analyses plotting pairwise $\Phi_{ST}$ distances.** A) Complete mtDNA MDS, B) HVR-1 MDS.

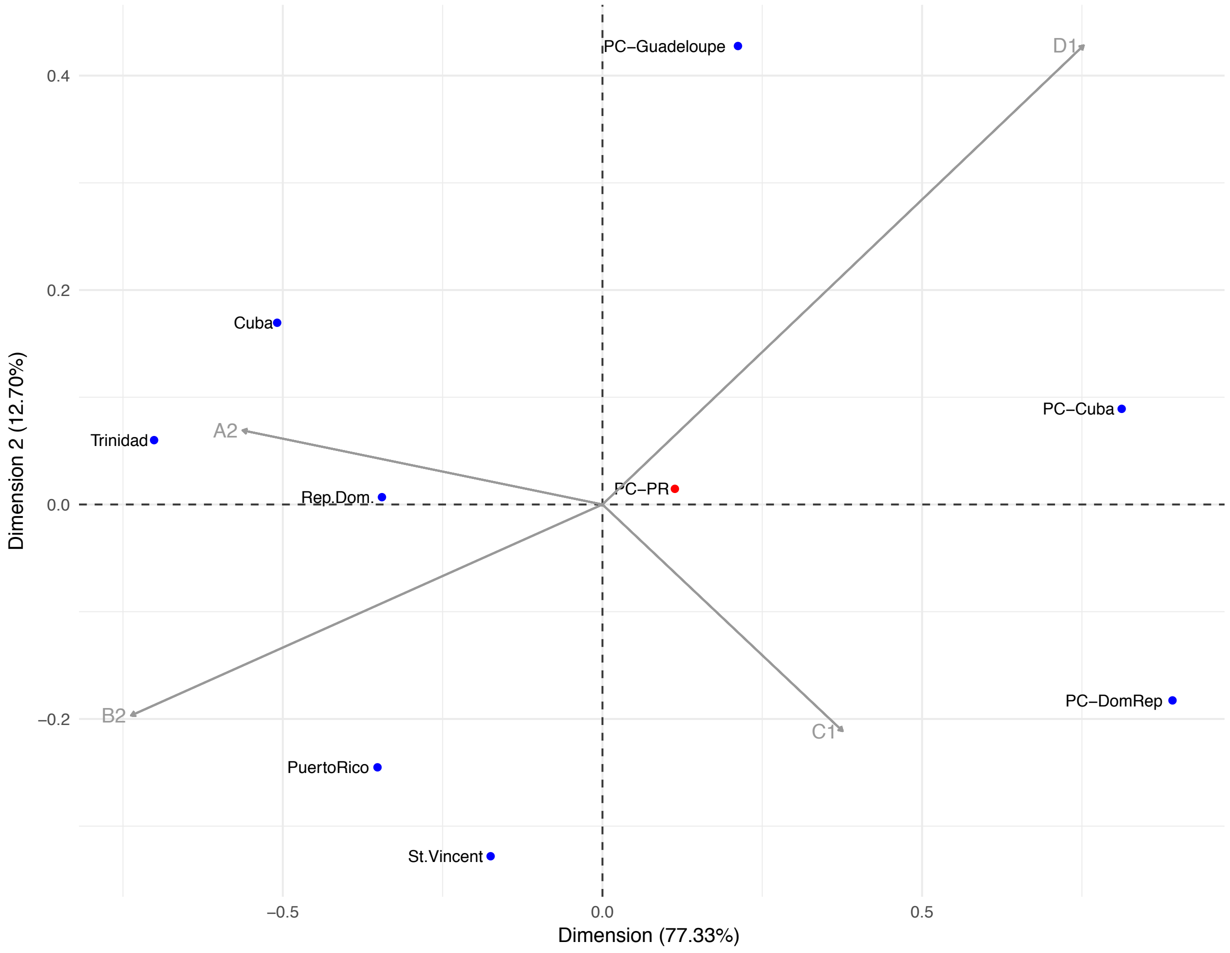


**Supplementary Figure S14. Correspondence analysis of haplogroup frequencies between Caribbean populations (HVR-1 data).**

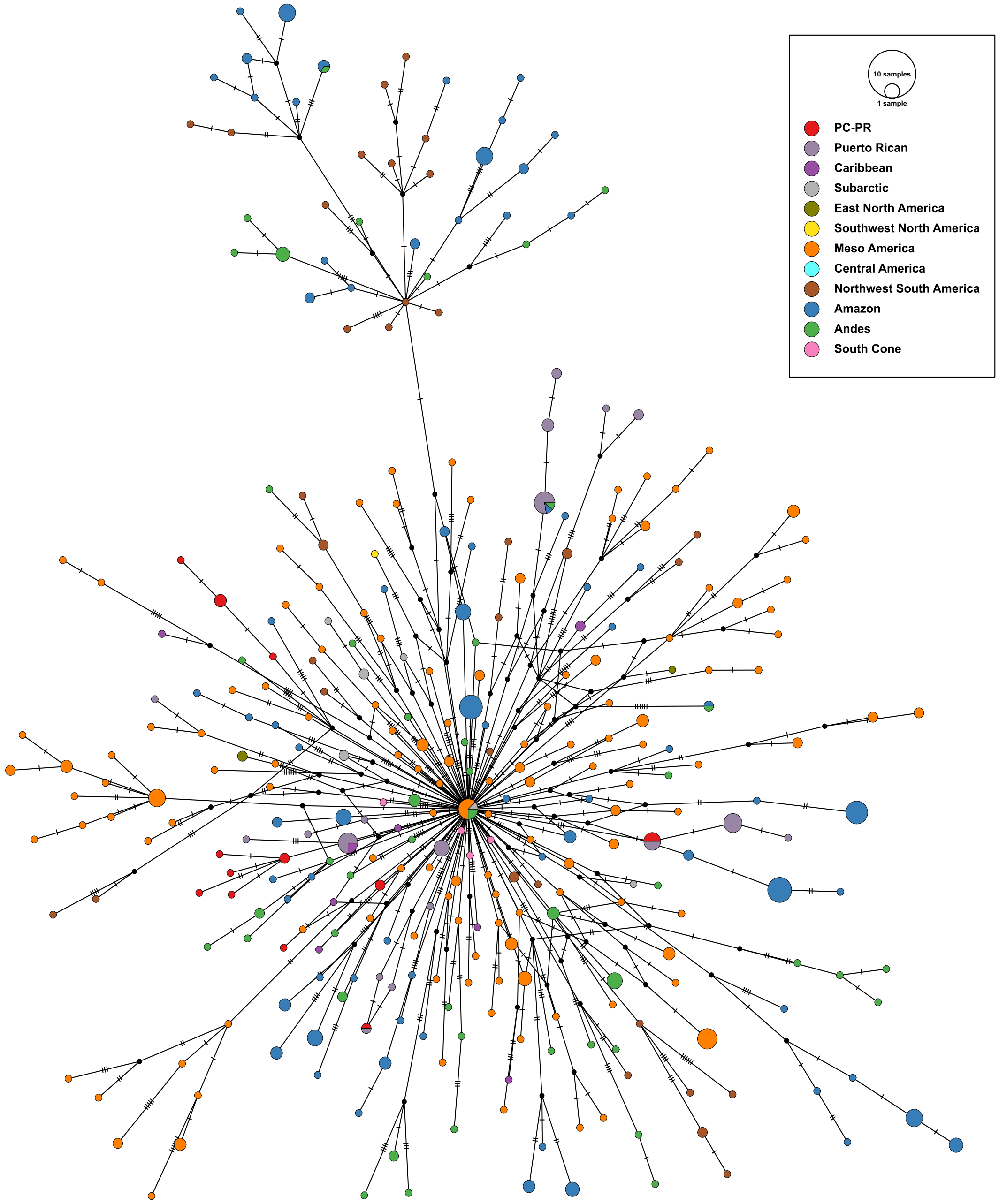


**Supplementary Figure S15. Median joining network of complete mtDNA diversity for haplogroup A.** Includes data from 519 ancient and present-day individuals from across the Americas. PC-PR shown in red, Puerto Ricans shown in lilac.

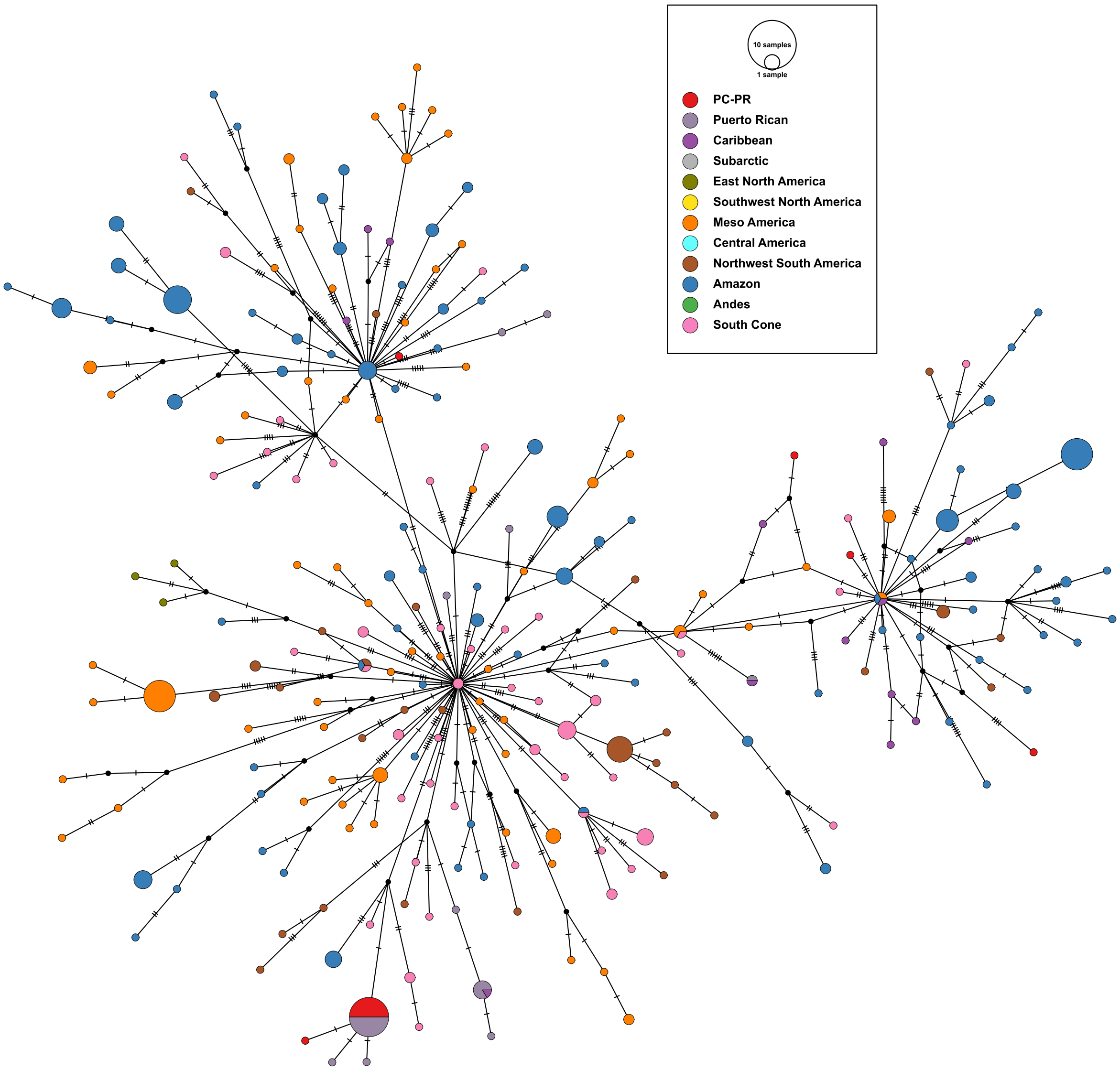


**Supplementary Figure S16. Median joining network of complete mtDNA diversity for haplogroup C.** Includes data from 456 ancient and present-day individuals from across the Americas. PC-PR shown in red, Puerto Ricans shown in lilac.

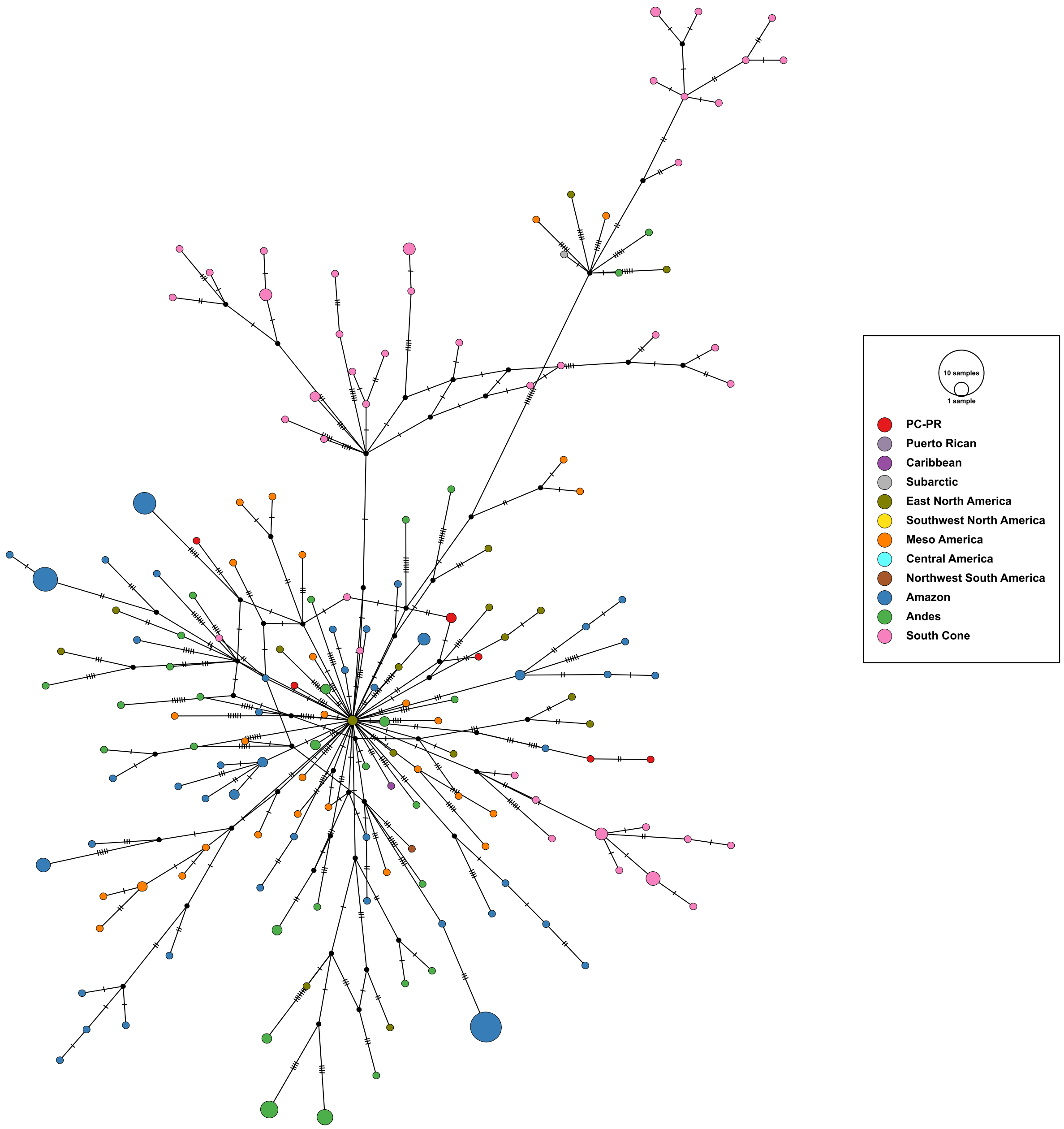


**Supplementary Figure S17. Median joining network of complete mtDNA diversity for haplogroup D.** Includes data from 423 ancient and present-day individuals from across the Americas. PC-PR shown in red. D1 sequences from present-day Puerto Ricans were not publicly available for inclusion in this analysis.

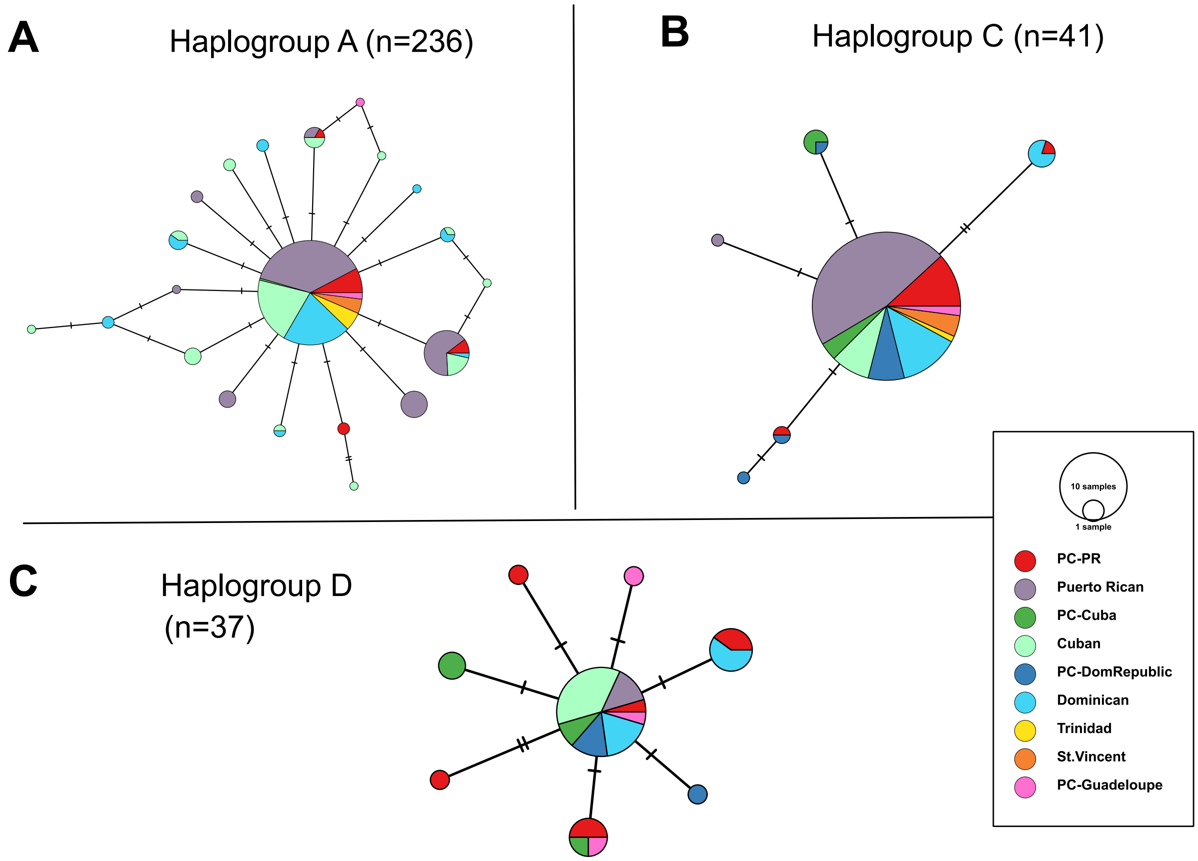


**Supplementary Figure S18. Median joining network of HVR-1 mtDNA diversity across the Caribbean.** Includes data from ancient and present-day individuals sampled in Puerto Rico, Cuba, Dominican Republic, Trinidad First People's Community, St. Vincent Garifuna and Guadeloupe. A) Haplogroup A: n=236, B) Haplogroup C: n=41, C) Haplogroup D: n=37.

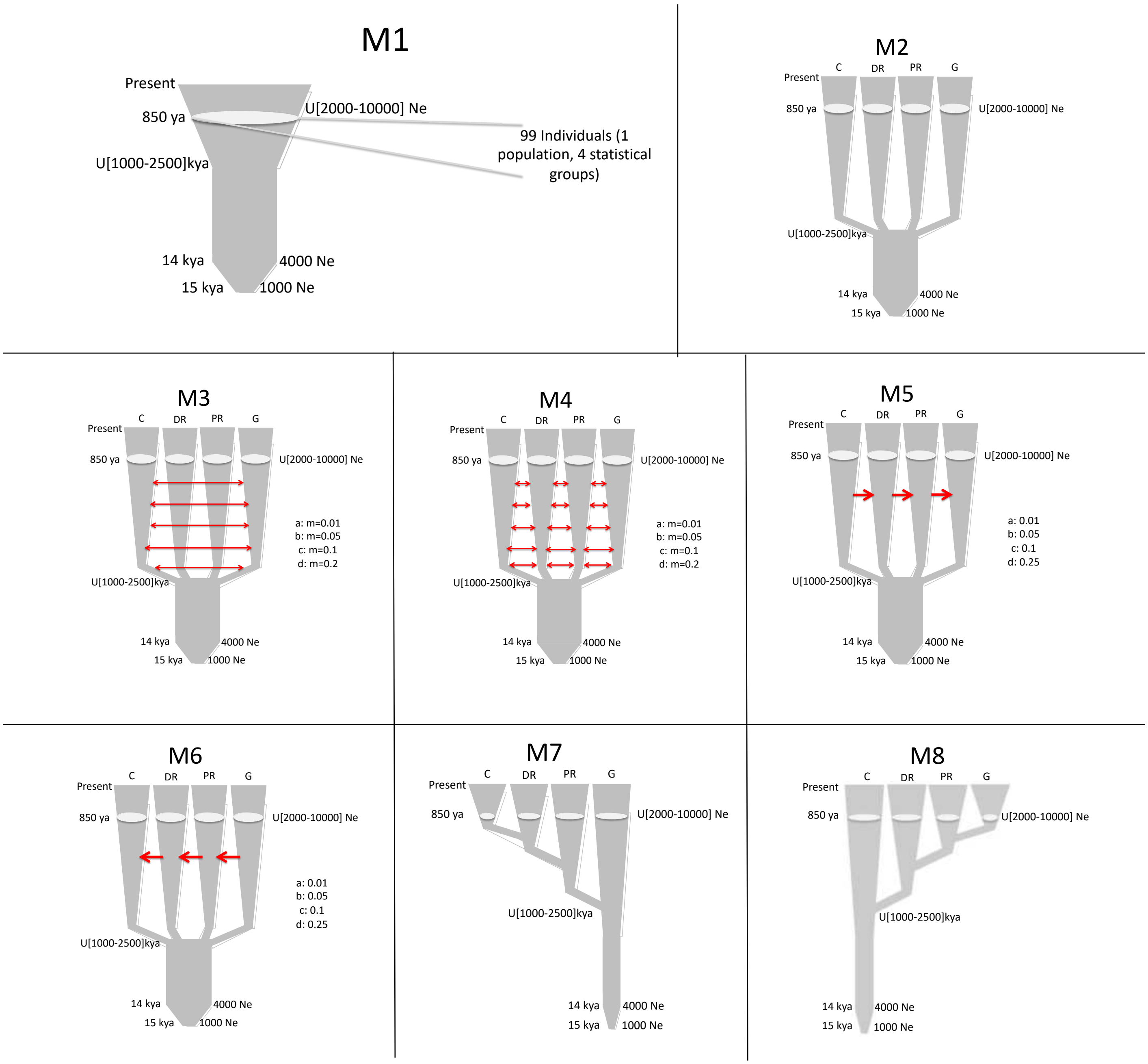


**Supplementary Figure S19. Demographic models of Caribbean population history tested with BayeSSC simulations.** Models simulate the available HVR-1 data from four pre-contact Caribbean populations under eight possible demographic scenarios varying the amount and direction of inter-island gene flow. C = Cuba, DR = Dominican Republic, PR = Puerto Rico, G = Guadeloupe.

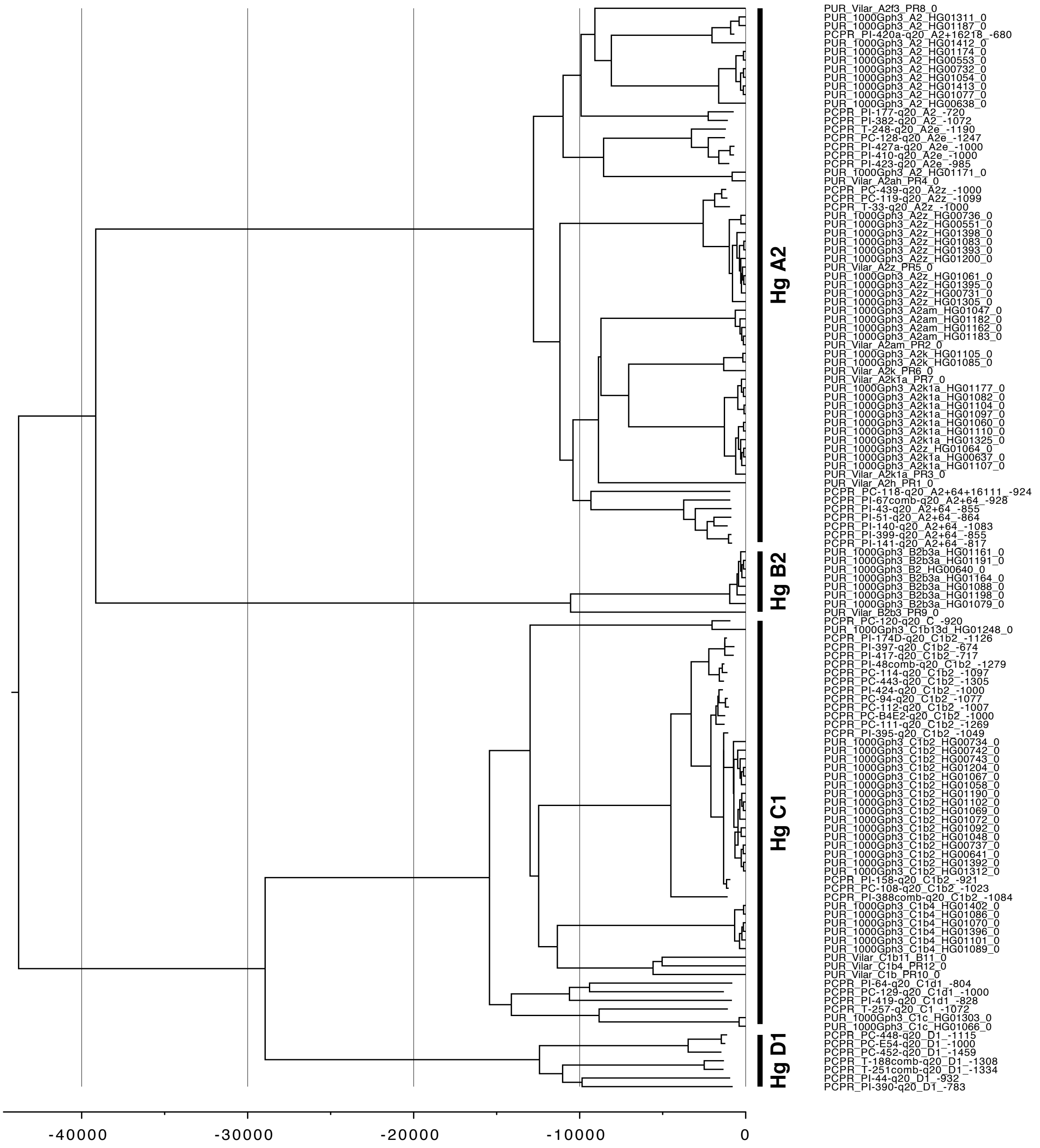


**Supplementary Figure S20. Median tree calculated with BEAST and including 127 complete mtDNA sequences from ancient and present-day Puerto Rico (Native American lineages only).** Branches leading to the main clades Hg A2, B2, C1, and D1 have support values of 1, all other branches have support values < 0.85.

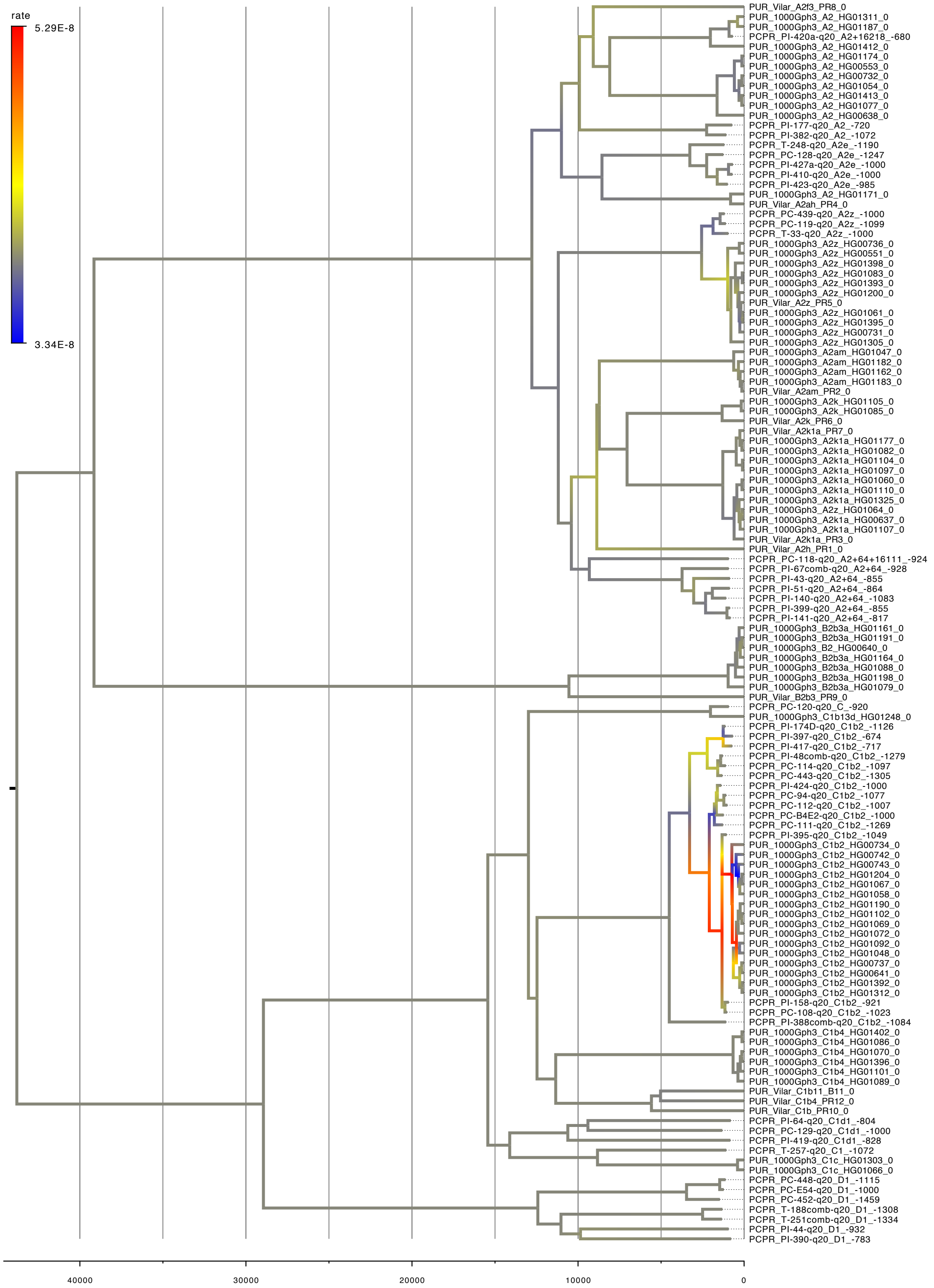


**Supplementary Figure S21. Median tree with rate along branches highlighted.**

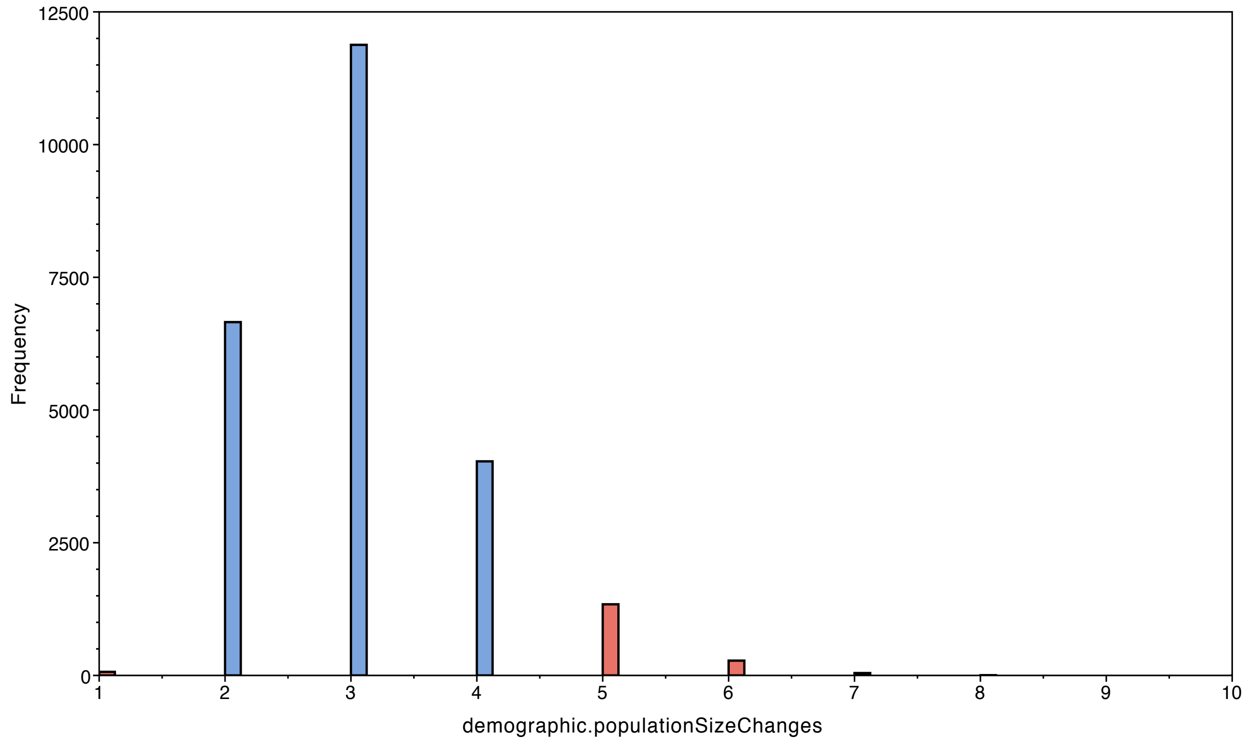


**Supplementary FigureS22: Posterior population size changes for All Sequences dataset (median: 3; 95% HPD: 2–5).** Constant population can be confidently rejected (0 is not in the 95% HPD).

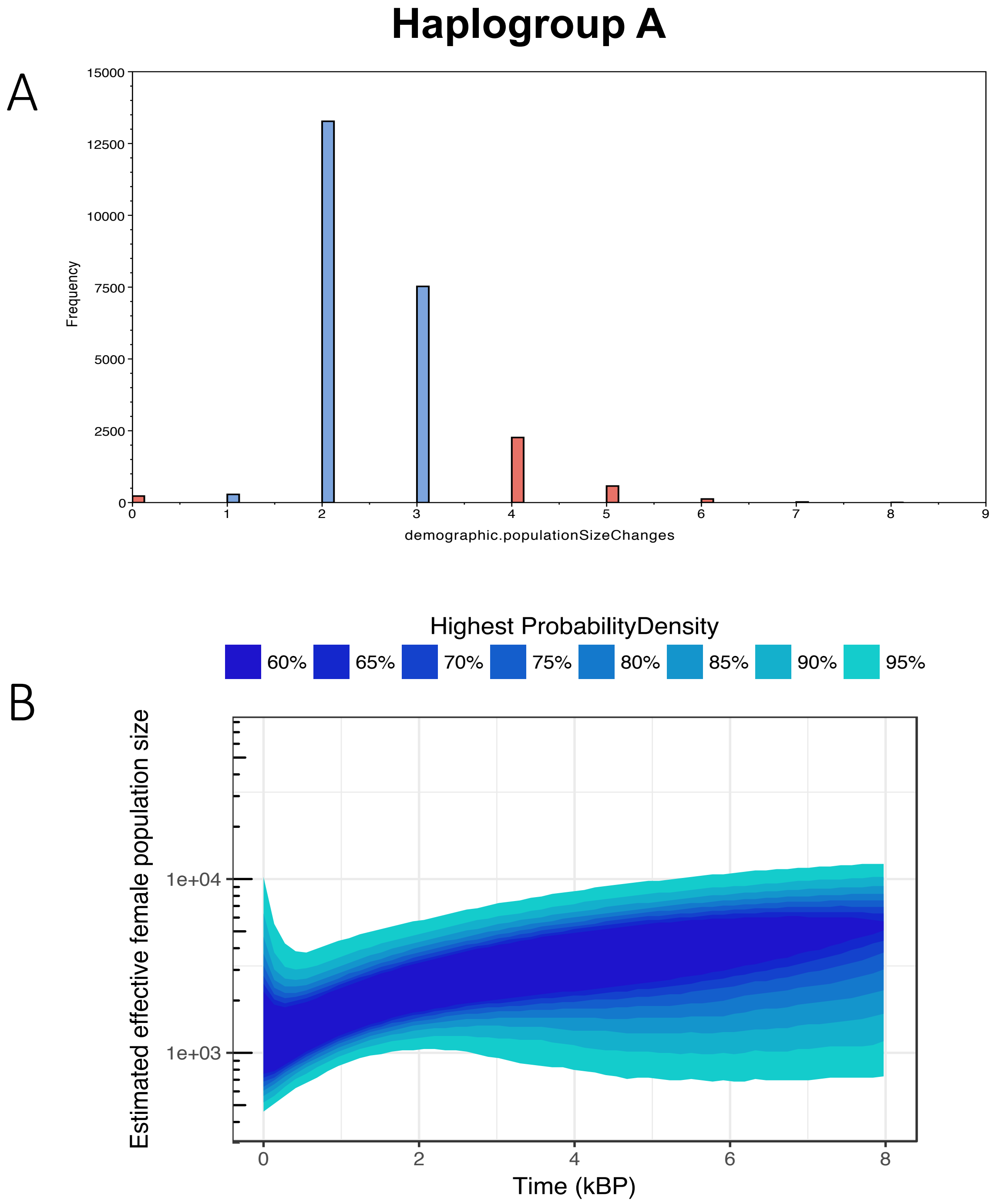


**Supplementary Figure S23. BEAST analysis results for Haplogroup A dataset.** A) Posterior population size changes (median: 2; % HPD: 1–4) for haplogroup A. Constant population can be confidently rejected (0 is not in the 95% HPD). B) Extended Bayesian skyline plot of female effective population size for haplogroup A, based on a generation time of 25 years.

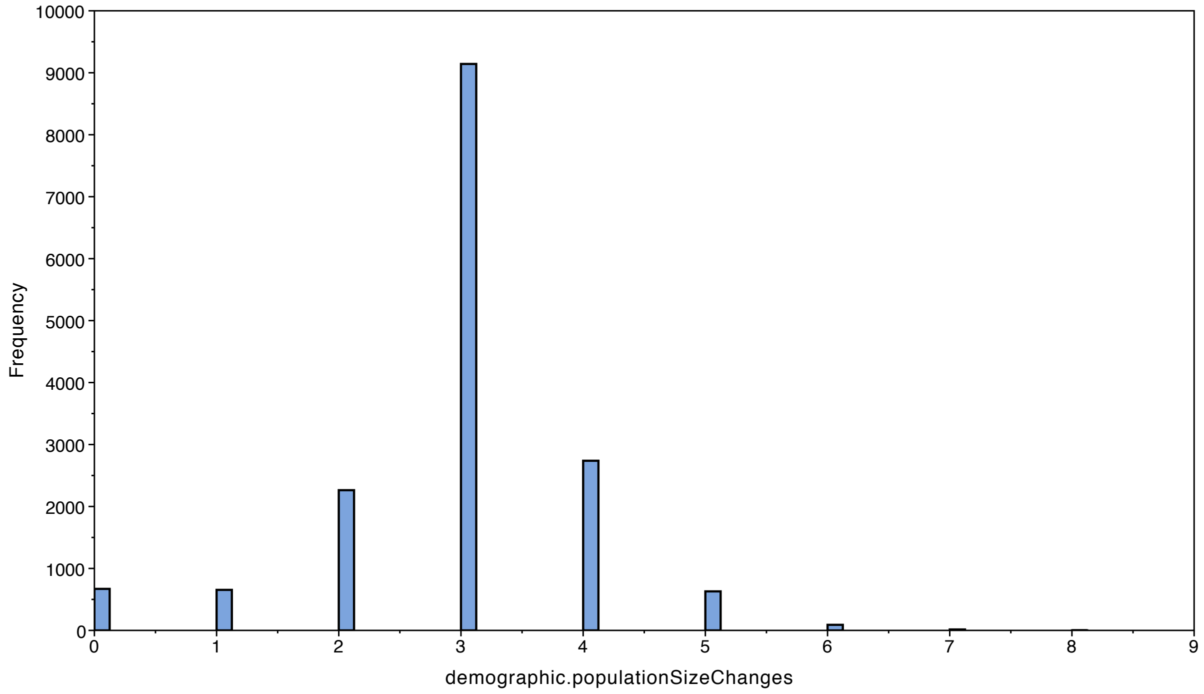


**Supplementary Figure S24 BEAST analysis results for Haplogroup C dataset.** Posterior population size changes (median: 3; % HPD: 0–4) for haplogroup C. Constant population cannot be confidently rejected (0 is in the 95% HPD), therefore the skyline reconstruction is meaningless.

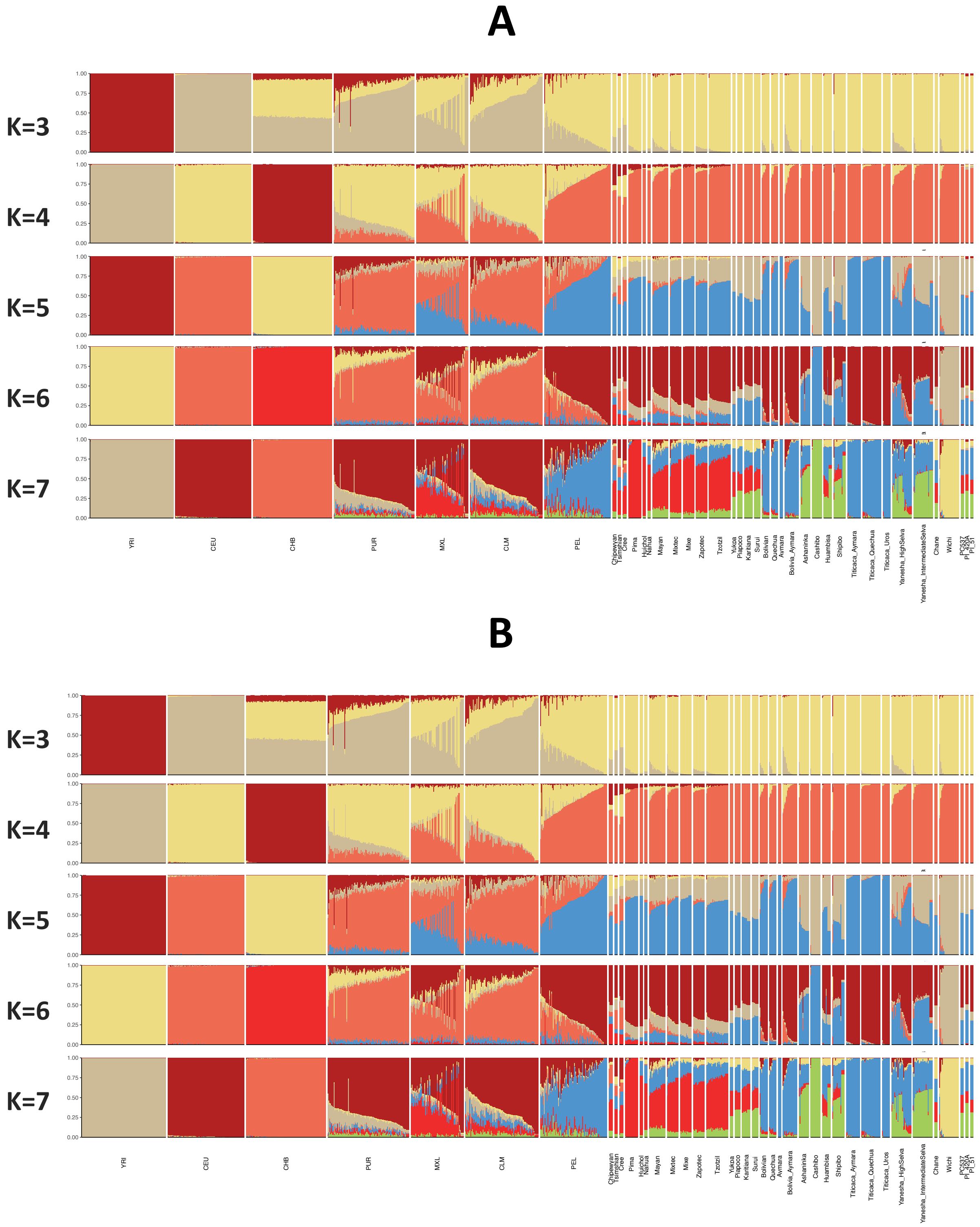


**Supplementary Figure S25. ADMIXTURE analysis (K3-K7) with PI-420a, PI-51 (PC-PR) and PC537 (Bahamas) autosomal genotypes.** A) Including all overlapping sites, B) including only transitions.

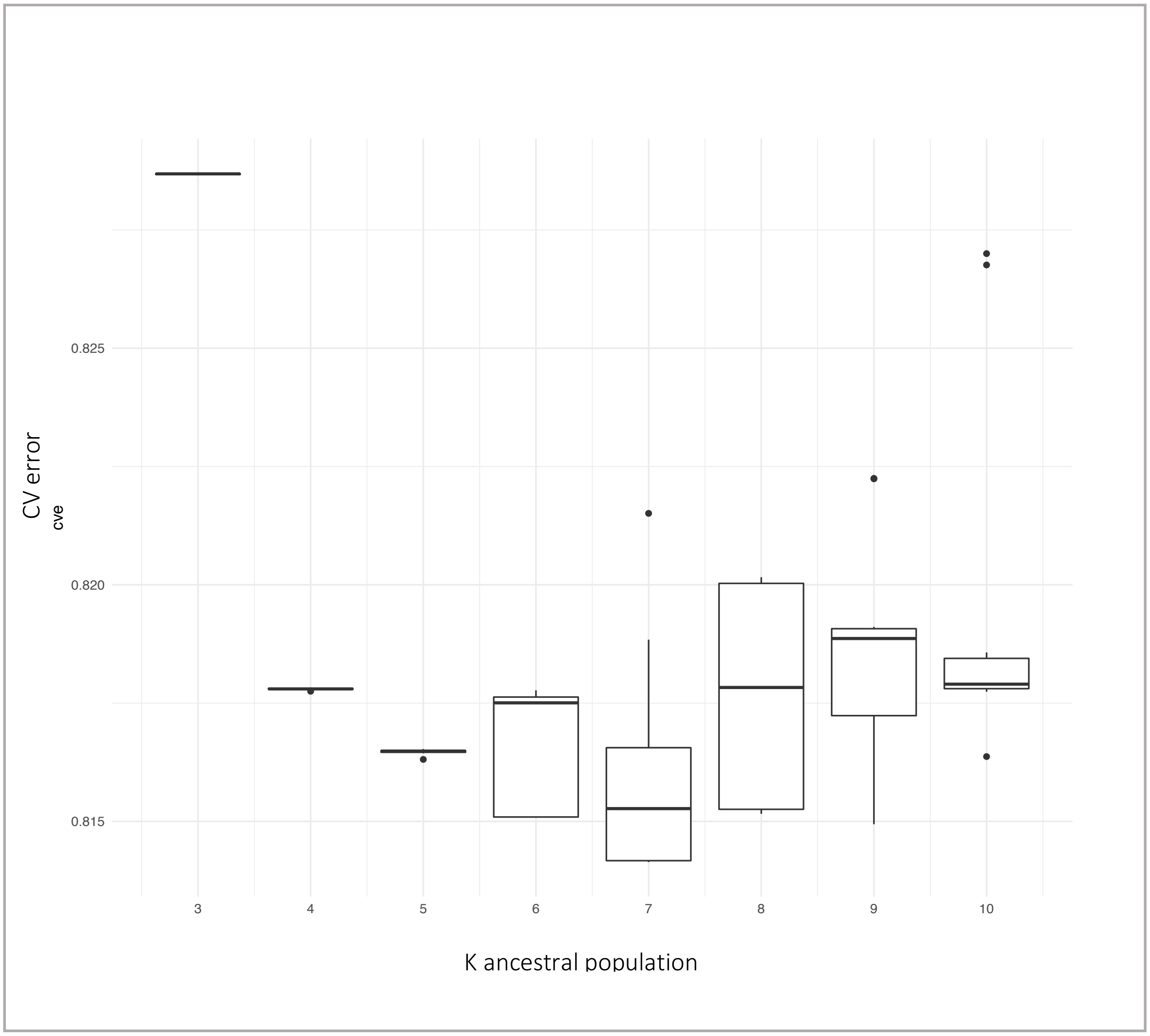


**Supplementary Figure S26. Cross validation errors associated with clustering analysis for runs K3=K7.**

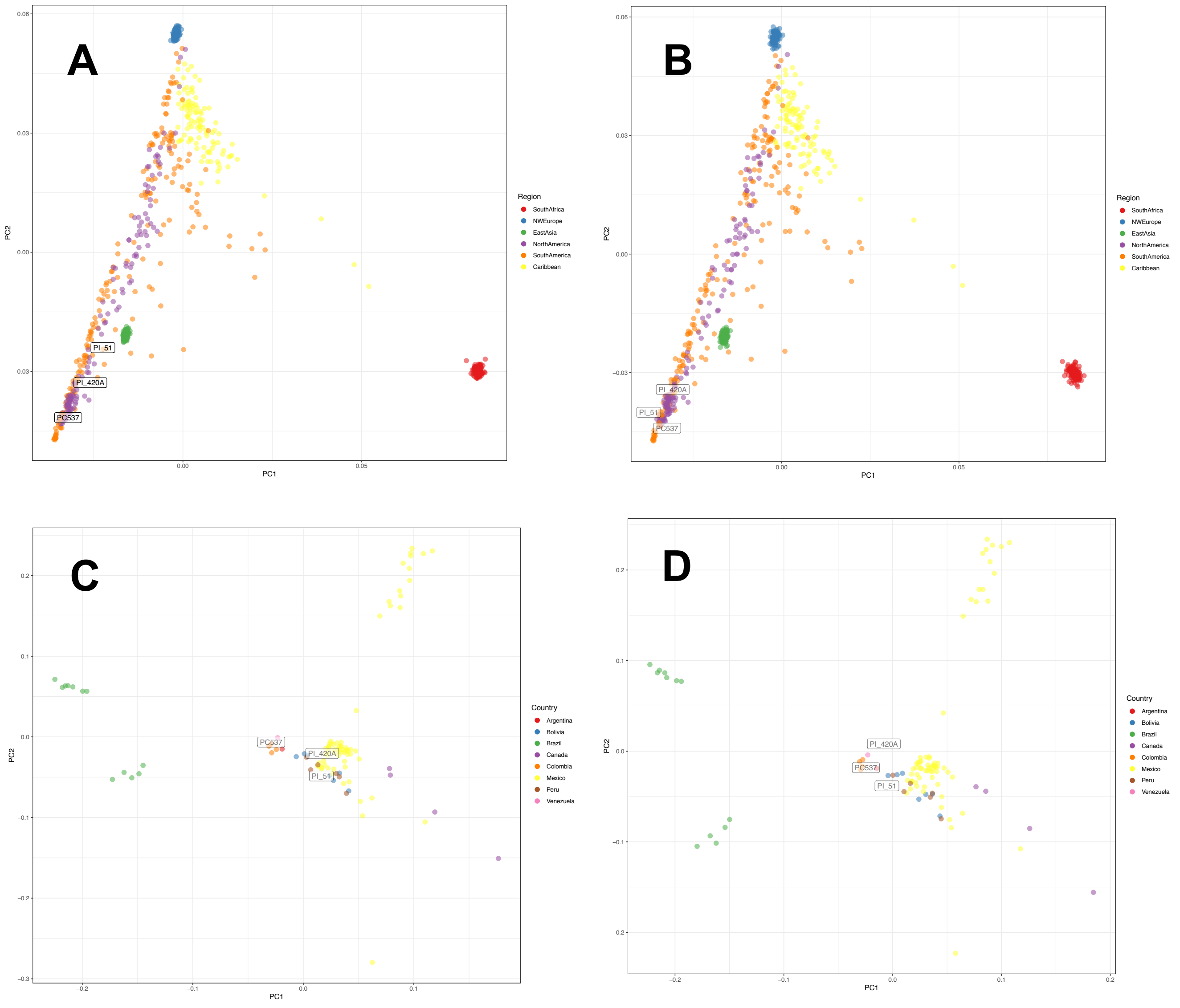


**Supplementary Figure S27. Principal components analyses (PCA) with autosomal genotypes.** Ancient samples are projected on eigenvectors calculated using reference panel individuals. Top panels: PCA comparing PI-420a, PI-51 (PC-PR) and PC537 (Bahamas) to world populations. A) Including all overlapping positions, and B) restricted to transitions only. Bottom panels: PCA comparing PI-420a, PI-51 (PC-PR) and PC537 (Bahamas) to Native American populations. C) Including all overlapping positions, and (D) restricted to transitions only.

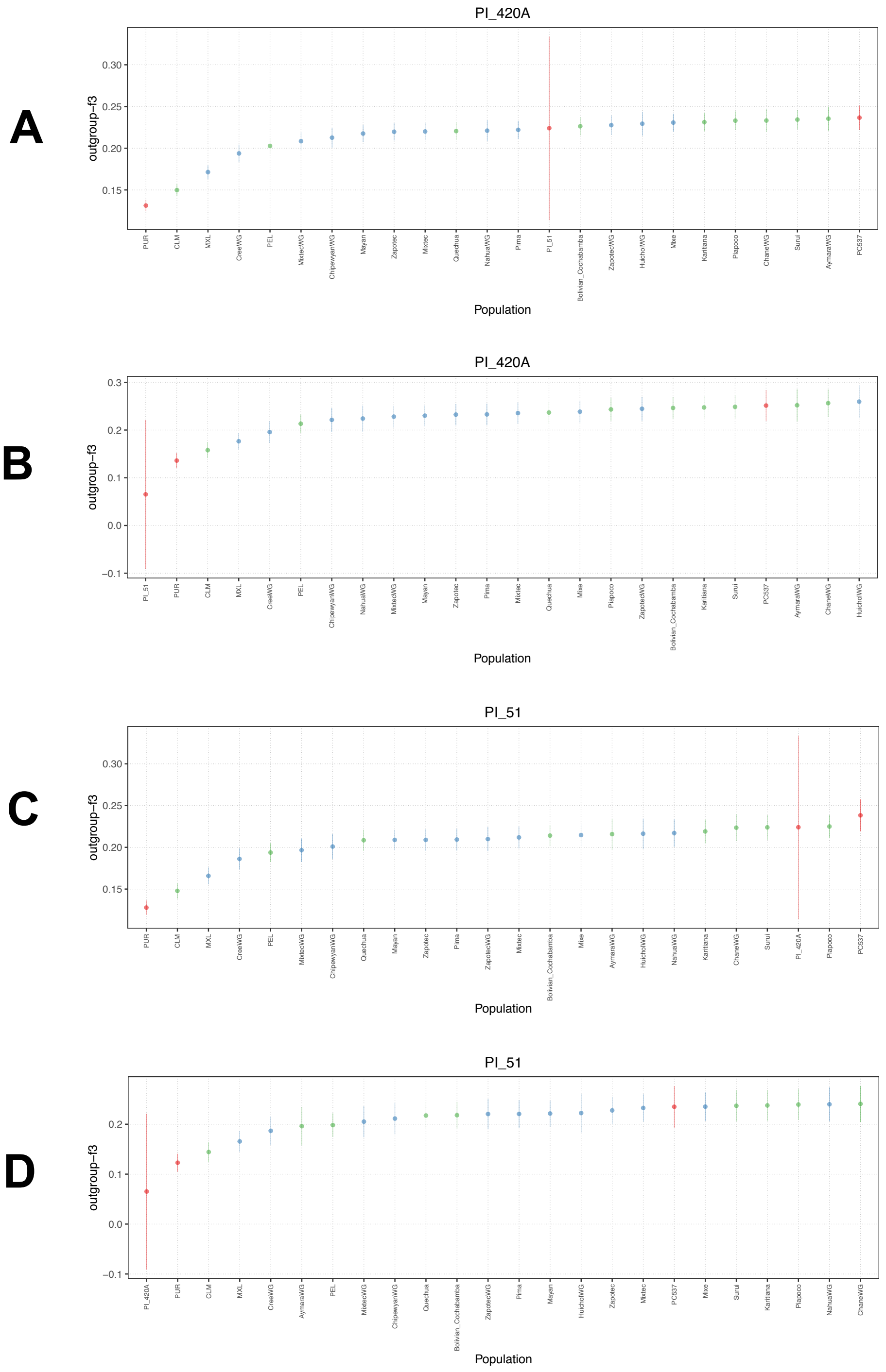


**Supplementary Figure S28. Outgroup $f_3$ statistics for two Paso del Indio individuals.** Statistics calculated in the form $f_3$(ancient, X; Yoruba). Error bars correspond to 95% standard error estimates. Populations are coloured by region: Blue = North America and Mesoamerica, Green = South America, Red = Caribbean. A) $f_3$(PI-420A, X; Yoruba) with all overlapping sites, B) $f_3$(PI-420A, X; Yoruba) with just transitions, C) $f_3$(PI-51, X; Yoruba) with all overlapping sites, D) $f_3$(PI-51, X; Yoruba) with just transitions.

### Supplementary Tables

**Supplementary Table S1. Ancient DNA sample information.**

**Supplementary Table S2. Extraction and library preparation information for ancient samples.**

**Supplementary Table S3. Post-mtDNA enrichment sequencing, haplogroup assignment and contamination analyses results.**

**Supplementary Table S4. Shotgun and whole genome enrichment sequencing statistics for ancient samples**

**Supplementary Table S5. Estimates of chromosomal sex and X-chromosome contamination calculated with ANGSD.**

**Supplementary Table S6. Results of contamination filtering on mitochondrial enriched samples.**

**Supplementary Table S7. Unique haplotypes in PC-PR sample.**

**Supplementary Table S8. Results from Fisher's Exact Test for genetic differentiation calculated with complete mtDNA data comparing the three study sites.**

**Supplementary Table S9. Pairwise PhiST matrix calculated with complete mtDNA data comparing the three study sites.**

**Supplementary Table S10. Comparative mtDNA data collected from the literature.**

**Supplementary Table S11. Complete mtDNA summary statistics for PC-PR and comparative populations from across the Americas.**

**Supplementary Table S12. MtDNA HVR1 summary statistics for PC-PR and comparative Caribbean populations (positions 16056 to 16391).**

**Supplementary Table S13. Results from Fisher's Exact Test for genetic differentiation calculated with complete mtDNA data comparing PC-PR to 46 ancient and modern populations from the Americas.**

**Supplementary Table S14. Pairwise PhiST matrix calculated with complete mtDNA data comparing PC-PR to 46 ancient and modern populations from the Americas.**

**Supplementary Table S15. Results from Fisher's Exact Test for genetic differentiation calculated with complete mtDNA data comparing PC-PR to eight ancient and modern populations from the Caribbean.**

**Supplementary Table S16. Pairwise PhiST matrix calculated with HVR-1 data comparing PC-PR to ancient and modern populations from the Caribbean.**

**Supplementary Table S17. AIC and likelihood goodness-of-fit results for demographic models simulated with BayeSSC.**

**Supplementary Table S18. TMRCA estimated from the BEAST analysis using all individuals.**

**Supplementary Table S19. Date estimates for individuals with no radiocarbon dates used in BEAST analysis.**

**Supplementary Table S20. Reference panel for autosomal DNA analyses. Data collected from the literature.**

**Supplementary Table S21. Number of overlapping positions included in autosomal data analyses.**
